# Multi-omics reveals a monocyte-macrophage-fibroblast axis in post-COVID-19 fibroinflammatory lung remodelling

**DOI:** 10.64898/2026.09.11.750491

**Authors:** Puja Mehta, Connar M. A. Mawer, Gordon Beattie, David J. F. Smith, Fadi Nikola, Hugh Selway-Clarke, Amy Y. Zhao, Taylor Adams, Annabel Kunzemann-Martinez, Stefan C. Stanel, Kaylee B. Worlock, Masahiro Yoshida, Rachel Walters, Sian Goldsworthy, Imran Uddin, Josephine L. Barnes, Gabrielle J. Rockett, Marko Z. Nikolić, Paolo Piazza, Manuela Platé, Iain Stewart, Joanna C. Porter, Arjun Nair, Joseph Jacob, Xiting Yan, Naftali Kaminski, Ahmed S. N. Alhendi, Rachel C. Chambers

**Affiliations:** Section of Rheumatology, Allergy, and Immunology, Department of Internal Medicine, Yale School of Medicine, New Haven, CT, USA; Department of Biomedical Informatics and Data Science (BIDS), Yale School of Medicine, New Haven, CT, USA; UCL Respiratory, Division of Medicine, University College London, London, UK; Department of Medicine, Imperial College London; Section of Pulmonary, Critical Care and Sleep Medicine, Department of Internal Medicine, Yale School of Medicine, New Haven, CT, USA; CRUK City of London Centre Single Cell and Spatial Genomics Facility, UCL Cancer Institute, University College London, London, UK; Bioinformatics Hub, UCL Cancer Institute, University College London, London, UK; National Heart and Lung Institute (NHLI), Imperial College London, London, UK; Department of Respiratory Medicine, University of Exeter, Exeter, UK; Great Ormond Street Institute of Child Health, Faculty of Population Health Sciences, University College London, London, UK; Genomics Translational Technology Platform, UCL Cancer Institute, University College London, London, UK; Nuffield Department of Medicine, Centre for Human Genetics, University of Oxford, Oxford, UK; Department of Radiology, University College London Hospitals NHS Foundation Trust, London, UK; Satsuma Lab, Hawkes Institute, University College London, London, UK; Department of Biostatistics, Yale School of Public Health, New Haven, CT, USA

## Abstract

Post-COVID-19 residual lung abnormalities (RLA) are associated with persistent respiratory symptoms and radiological changes, yet the underlying mechanisms remain unclear. We performed integrated multi-omic profiling of paired bronchoalveolar lavage and blood samples from patients with post-COVID-19 RLA and healthy controls, combining single-cell RNA sequencing, CITE-seq, single-cell T cell receptor sequencing, bronchoalveolar lavage fluid proteomics and functional fibroblast assays. In post-COVID-19 RLA lungs, we identified an increased abundance of profibrotic SPP1hi monocyte-derived alveolar macrophages, arising from an expanded circulating HLA-DRlowCD163+PDE4Dhi classical monocyte progenitor population, supporting a blood-lung myeloid axis. Cell-cell communication modelling positioned macrophages as central hubs of immune-stromal crosstalk, promoting monocyte recruitment with profibrotic priming, and fibroblast activation. Proteomic analysis of bronchoalveolar lavage fluid from post-COVID-19 RLA and idiopathic pulmonary fibrosis, compared with healthy controls, revealed shared and distinct signatures. These alveolar proteins in post-COVID-19 RLA were predominantly attributed to myeloid cells and predicted to engage fibroblast receptors. Bronchoalveolar lavage fluid induced fibroblast proliferation, differentiation and collagen deposition in vitro, with proliferation attenuated by the antifibrotic drug nintedanib. We also identified compartment-specific lymphoid dysregulation, including depletion of mucosal-associated invariant T (MAIT) cells in both the lung and blood, decreased natural killer (NK) cells with oligoclonal T cell expansion in the lung, and expansion of regulatory and cytotoxic T cells in the blood. These findings support a persistent monocyte-macrophage-fibroblast axis linking immune dysregulation to fibroproliferative remodelling after COVID-19 and highlights candidate therapeutic targets for post-viral lung fibrosis. We provide a publicly available atlas at x (TBA).

## Main

Acute lung injury may culminate in fibrosis, characterised by aberrant tissue repair, immune dysregulation and fibroblast activation. The COVID-19 pandemic, caused by SARS-CoV-2, provided a tractable human context to study the fibroinflammatory consequences of severe viral pneumonitis. Post-acute sequelae of SARS-CoV-2 infection (PASC) describes persistent symptoms and organ dysfunction following COVID-19^1^. A subset of these patients have post-COVI D-19 residual lung abnormalities (RLA), with persistent respiratory symptoms and radiological changes. However, the alveolar and systemic immune landscape is not well-defined and understanding this biology is important because therapeutic interventions remain elusive^2^.

Recent studies that assessed the immune profile of acute severe COVID-19^3,4^ and post-COVI D lung disease^5,6^, focused mainly on altered transcriptional states of circulating monocytes^7-10^, as well as the presence of transcriptionally distinct profibrotic SPP1⁺ monocyte-derived alveolar macrophages (MoAMs) previously observed in patients with idiopathic pulmonary fibrosis (IPF)^11^ and in fibrotic hypersensitivity pneumonitis^12^. These findings suggest conserved mechanisms with potential therapeutic implications across fibrotic lung disease. However, in post-COVID-19 RLA, the relationship between systemic and local pulmonary immunopathology, their overlap, and their contribution to tissue phenotype remain poorly understood. Defining these relationships could identify blood-accessible signatures of persistent lung immune activity and fibrotic remodeling.

In this study we sought to understand the innate and adaptive immune cell landscape in the lung and blood and the immune-stromal cross-talk with fibroblasts that may promote lung remodelling. We performed single-cell RNA sequencing (scRNA-seq), T cell receptor sequencing (scTCR-seq) and cellular indexing of transcriptomes and epitopes by sequencing (CITE-seq), using paired bronchoalveolar lavage (BAL) and blood samples from patients with post-COVID-19 RLA and healthy controls (Figure 1a), as well as BAL proteomics and functional *in vitro* bioassays, from the same patients. Among myeloid cell populations we found enrichment of *SPP1*⁺ profibrotic alveolar macrophages, likely arising from an expanded circulating HLA-DR^low^CD163^+^PDE4D^hi^ classical monocyte precursor population. BAL fluid (BALF) proteomic profiling revealed shared and distinct inflammatory cytokine and chemokine networks in post-COVID-19 RLA and IPF. BALF proteins and alveolar macrophage ligands were predicted to engage with fibroblast receptors. Functional bioassays revealed that BALF induced fibroblast proliferation (which was attenuated by the antifibrotic drug, nintedanib), as well as fibroblast differentiation and collagen deposition. Among lymphoid populations, we found compartment-specific dysregulation including depletion of MAIT cells in both the lung and blood, decreased NK cells with oligoclonal T cell expansion in the lung, and expansion of regulatory and cytotoxic T cells in the blood.

**Figure 1.**
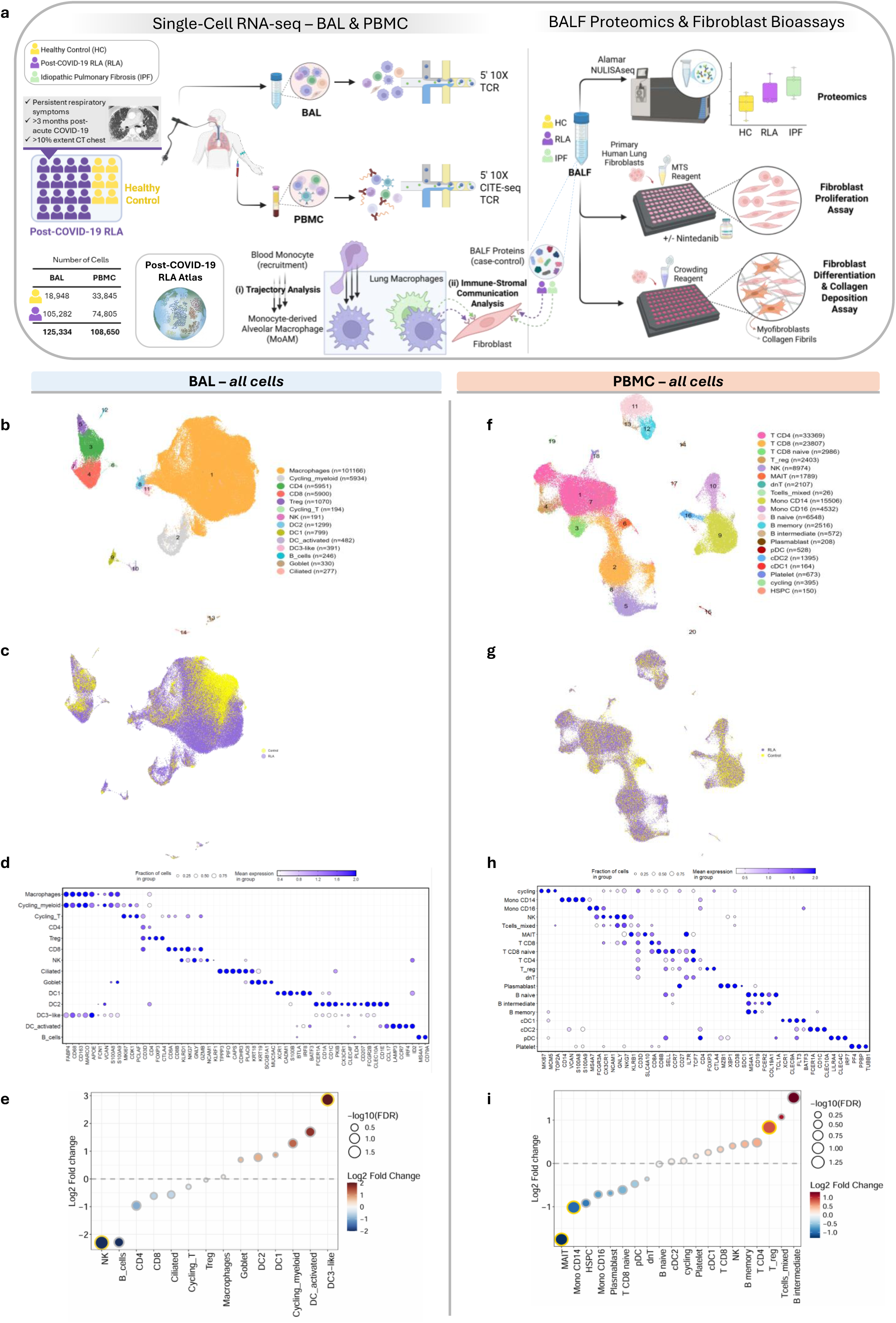
Study design and single cell clustering and cell compositional results of the bronchoalveolar and systemic cellular landscape in patients with post-COVID-19 RLA. **a** Study overview. Paired bronchoalveolar lavage (BAL) and peripheral blood mononuclear cells (PBMCs) were obtained from patients with post-COVID-19 residual lung abnormalities (RLA; n=16) and healthy controls (HC; n=6). Single-cell RNA sequencing (scRNA-seq) with 5⍰ chemistry and T cell receptor (TCR) sequencing was performed on BAL and PBMC samples, with CITE-seq applied to PBMCs. Trajectory analysis was used to investigate blood monocyte transition to monocyte-derived alveolar macrophages (MoAM). Immune–stromal communication analyses were conducted using macrophage–fibroblast interactions from post-COVID-19 and idiopathic pulmonary fibrosis (IPF) data and differentially expressed BAL fluid (BALF) proteomics-fibroblast. BALF proteomics was performed using the Alamar Nucleic Acid Linked Immuno-Sandwich Assay (NULISA)-seq platform. BALF was applied to primary human lung fibroblasts to assess proliferation (MTS and DAPI cell counts) and the effect of the anti-fibrotic agent, nintedanib, and fibroblast differentiation and collagen deposition using macromolecular crowding assays. **b-e** BAL and **f-I** PBMC Single-cell clustering and compositional analysis **b-c** BAL and PBMC **f–g** Uniform manifold approximation and projection (UMAP) visualization of transcriptomes, annotated by major cell types (**b,f**) and coloured by condition (**c,g**). post-COVID-19 RLA = purple; Healthy Control = yellow. Cell plotting order was randomized between conditions to improve visualization **d,h**. Dot plots show canonical marker genes used for cell-type annotation for BAL (**d**) and PBMC (**h**). **e** BAL and **i** PBMC Differential cell abundance analysis comparing post-COVID-19 RLA and HC samples Log₂ fold change (LFC) i s shown, with red indicating increased and blue decreased abundance in RLA. Circle size reflects –log₁₀(FDR) from Wilcoxon rank-sum test with Bonferroni correction (FDR < 0.05, indicated by yellow border). Dashed line indicates LFC = 0. RLA = post-COVID-19 Residual Lung Abnormalities, T_reg_= regulatory T cells; MAIT = mucosal-associated invariant T cells; NK= natural killer cells; pDCs = plasmacytoid dendritic cells; DC2 = dendritic cells (type 2); DC1 = dendritic cells (type 1); DC activated = activated dendritic cells; dNT = double negative T cells; Mono = monocytes; HSPC = haematopoetic progenitor cells

Together, these findings reveal persistent fibroinflammatory immune remodelling and implicate dysregulated monocyte–macrophage–fibroblast interactions and immune-stromal cross-talk in post-COVID-19 RLA. We provide a publicly available atlas (TBA).

## Results

### Persistent bronchoalveolar and systemic immune perturbations in post-COVID-19 RLA

The prospectively recruited study cohort included sixteen patients with post-COVID-19 RLA and six healthy control subjects (Table 1,, Extended Data Table 1; Extended Data Table 2), with paired BAL and blood sampling. Patients with post-COVID-19 RLA had persistent respiratory symptoms, >10% interstitial abnormalities on high-resolution computed tomography (HRCT) chest scans and were sampled at a median of 8.5 months after initial (first-episode) COVID-19 symptom onset. Healthy control participants were asymptomatic with normal chest CT imaging and had no prior infection with SARS-CoV-2 in the previous 24 months. Patients with pre-existing lung disease or receiving steroids of any formulation within 4 weeks were excluded from the study. Full eligibility criteria are outlined in Extended Data Table 3, Extended Data Table 4. All BAL samples underwent microbiological screening, to exclude bacterial infection.

**Table 1.** Clinical features ofthe cohort. Data are median (range), n (%), or n/N (%). Percentages might not total 100 where expected owing to rounding. HV = Healthy volunteers; BAL = Broncholveolar lavage; BMI = Body Mass Index; ECMO = Extracorporeal Membrane Oxygenation; I&V = Intubated and ventilated; CPAP = Continuous Positive Airways Pressure; HFNO = High-flow nasal oxygen;

|  | post-COVID-19 RLA All<br>(n=16) | Controls<br>(n=6) | p |
| --- | --- | --- | --- |
| Age |  |  |  |
| Years | 60 (25-79) | 47.5 (28-71) | 0.112 |
| Sex |  |  |  |
| Male | 10 (63%) | 5 (83%) | 0.674 |
| Female | 6 (38%) | 1 (17%) |  |
| Race |  |  |  |
| White | 9 (56%) | 5 (83%) | 0.055 |
| Asian | 7 (44%) | 0 |  |
| Black | 0 | 1 (17%) |  |
| BMI |  |  |  |
| kg/m<sup>2</sup> | 30 (21-49) | 27 (21-30) | 0.161 |
| Smoking status |  |  |  |
| Current | 1 (6%) | 1 (17%) | 0.341 |
| Former | 2 (13%) | 2 (33%) |  |
| Never | 13 (81%) | 3 (50%) |  |
| Respiratory support in acute COVID-19 |  |  |  |
| ECMO | 3 (19%) | NA |  |
| I&V | 8 (50%) | NA |  |
| CPAP | 2 (13%) | NA |  |
| HFNO | 2 (13%) | NA |  |
| Nil | 1 (6%) | NA |  |
| WHO Severity Score |  |  |  |
|  | 8 (3-9) | NA |  |
| Treatment for acute COVID-19 |  |  |  |
| Steroid | 14 (88%) | NA | 1 * |
| Tocilizumab | 4 (25%) | NA | 0 * |
| Timing of bronchoscopy post-acute COVID-19 |  |  |  |
| Months | 8.5 (3-13) | NA |  |

We generated a dataset of 233,984 single cells across BAL and PBMC compartments (Figure 1a-h, Supplementary Figure 1). There were disease-associated shifts in immune cell composition between patients with post-COVID-19 RLA and healthy controls (Figure 1 c, g, Supplementary Figure 2). In the BAL, DC3 cells were significantly increased whereas NK cells were decreased (Figure 1e), and MAIT cells were significantly reduced in both compartments (Figure 1i, Figure 4i). In the blood, CD14⁺ monocytes were significantly decreased, while Tregs and CD8⁺ cytotoxic T cells were enriched (Figure 1g, Figure 4i). These findings suggest compartment-specific immune perturbations months after acute infection.

### Profibrotic SPP1^hi^ alveolar and cycling macrophages are enriched in post-COVID-19 RLA

Myeloid reclustering revealed disease-specific shifts in post-COVID-19 RLA compared with healthy controls (Figure 2a-d; f-i). In the BAL, post-COVID-19 RLA SPP1^hi^ macrophages were significantly enriched (Figure 2d) and shared transcriptional features with profibrotic MoAMs previously described in acute COVI D-19^4,13-15^ and IPF^11,16-18^ (Supplementary Figure 3a,b). These cells co-expressed genes implicated in matrix remodelling and fibrosis (MMP9, CHIT1, and RETN), consistent with a fibrotic effector phenotype (Supplementary Figure 3c).

**Figure 2.**
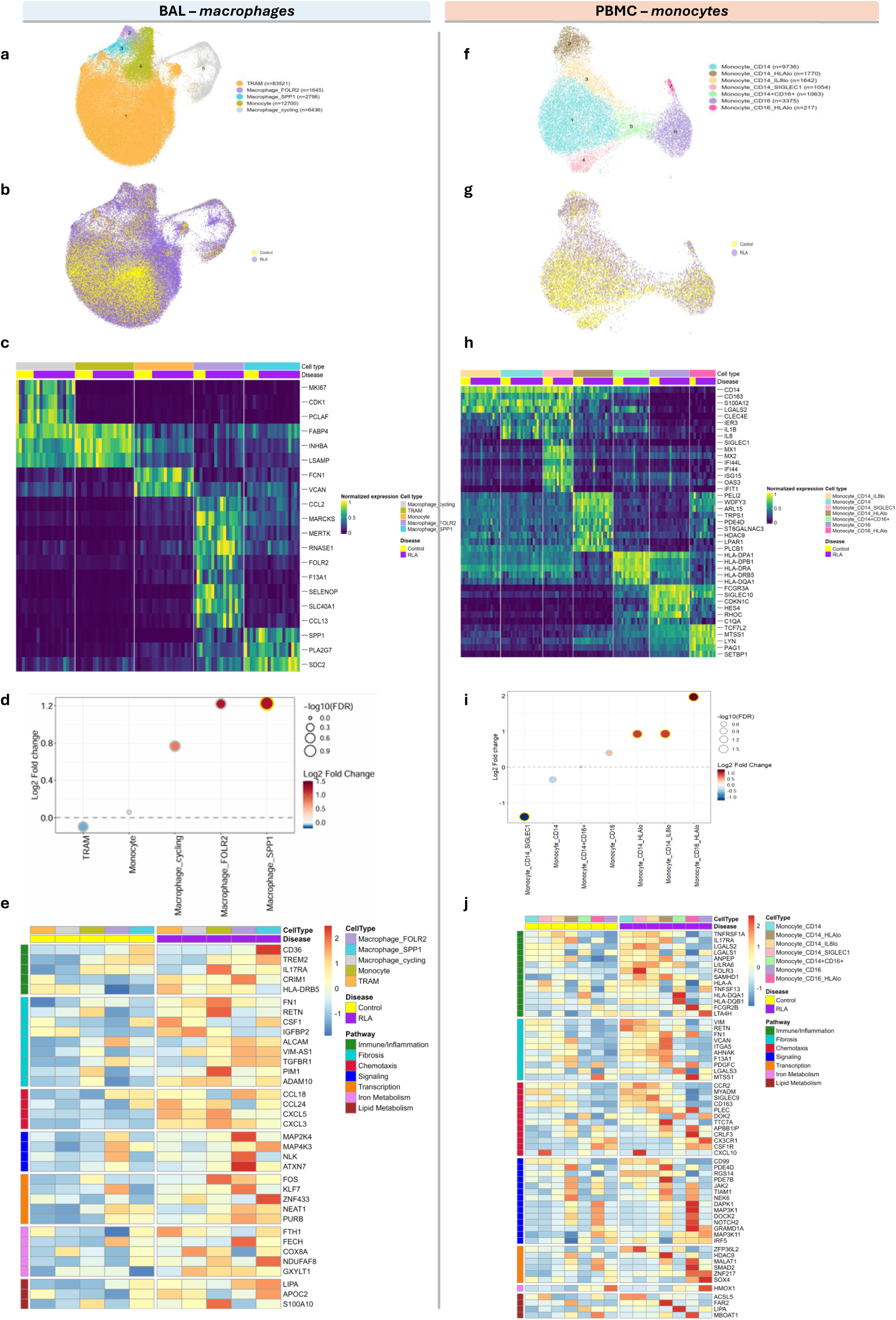
Myeloid single-cell analysis of bronchoalveolar and blood compartments in patients with post-COVID-19 Residual Lung Abnormalities. **a–e** BAL and **f-j** PBMC single-cell clustering and differential abundance analysis of myeloid cells. **a, b** BAL and **d, e** PBMC uniform manifold approximation and projection (UMAP) visualization of myeloid labelled by cell type (**a, f**); cell numbers in parentheses) or condition (**b, e)**; Healthy Control = yellow, post-COVID-19 RLA = purple **c, f** Differential cell abundance analysis comparing post-COVID-19 RLA vs Healthy Control in BAL (**c**) and PBMC (**f**). Log₂ fold change (LFC) is shown, with red indicating increased and blue decreased abundance in RLA. Circle size reflects –log₁₀(FDR) from Wilcoxon rank-sum test with Bonferroni correcti on (FDR < 0.05, indicated by yellow border). Dashed line indicates LFC = 0. **g, h** Heatmaps of macrophage/monocyte subset marker genes in BAL ( **g**) and PBMC (**h**). Genes ranked by FDR-adjusted p-value from Wilcoxon rank-sum test (per subject average expression), comparing each subset against all others. Gene expression values are unity normalized from 0 to 1 across rows within each subset. Each column represents the average expression value for one individual. **i, j,** Differentially expressed genes in BAL (**i**) and PBMC (**j**) myeloid cells comparing post-COVID-19 RLA vs Healthy Control. Gene expression values represent Z-score normalized log-normalized counts, where each gene’s expression is mean-centred and scaled across all samples. The colour scale reflects row-wise Z-scored expression, with red indicating upregulation and blue indicating downregulation in RLA rel ative to controls. DEGs were identified using Seurat’s FindMarkers function (Wilcoxon test; log₂ fold change > 0.25; FDR < 0.05; Bonferroni correction), excluding ribosomal genes. Top annotation indicates myeloid cell type and disease group (Healthy Control = yellow, post-COVID-19 RLA = purple); right annotation categorizes DEGs by pathway. RLA = post-COVID-19 Residual Lung Abnormalities, TRAM = tissue-resident alveolar macrophages.

Subclustering of cycling alveolar macrophages (defined by *MKI67, UBE2C, and TOP2A*) identified two distinct *SPP1*⁺ cycling macrophage subsets (Supplementary Figure 4 a-c). One subset (cluster 9) was present exclusively in post-COVID-19 RLA and co-expressed matrix remodelling and inflammatory genes, including *MMP9, CHIT1* and *CXCL10*, consistent with a profibrotic transcriptional programme, previously described in fibrotic regions of IPF lungs^16^.

### Circulating monocyte remodelling in post-COVID-19 RLA

In peripheral blood, monocyte composition was markedly altered in post-COVID-19 RLA compared with controls (Figure 2f). Classical CD14⁺ monocytes were reduced, alongside a type I interferon–activated subset characterised by expression of the interferon-inducible receptor *SIGLEC1* (sialoadhesin / *CD169*), *IFIT1,* and *IFI44*.

In contrast, several monocyte states were enriched in post-COVID-19 RLA, including an HLA-DR^lo^PDE4D^hi^CD163^+^ classical monocyte population, CD14^+^IL8^low^ classical monocytes and CD16^+^HLA-DR^low^ non-classical monocytes. The HLA-DR^lo^PDE4D^hi^CD163^+^ classical monocyte subset exhibited reduced MHC class II expression (HLA-DPA1, HLA-DRB5, HLA-DRA) and upregulation of PELI2, consistent with an altered innate immune activation state.

### Transcriptional programmes indicate profibrotic alveolar macrophages and systemic monocyte priming

Differential gene expression of post-COVID-19 RLA and healthy controls revealed compartmentalised myeloid activation programs. In the BAL (Figure 2e), alveolar macrophages exhibited fibrosis- and inflammation-associated transcriptional signatures, including *TGFBR1*, *TREM2*, *RETN*, *CSF1*, *VIM-AS1*, and *PIM1*, particularly within SPP1⁺, FOLR2⁺ and cycling macrophage subsets. Chemotactic mediators (*CCL18*, *CCL24*, *CXCL3*, and *CXCL5*) and lipid and iron metabolism genes (*LIPA*, *APOC2*, *FTH1*) were enriched, suggesting immune recruitment and metabolic reprogramming. In peripheral blood (Figure 2j), classical monocytes demonstrated transcriptional features of migratory priming, including increased expression of *CCR2* and *CX3CR1*.

Together, these findings suggest coordinated myeloid remodelling across compartments, with profibrotic macrophage expansion in the lung and circulating monocyte states transcriptionally primed for recruitment.

### Circulating HLA-DR^low^PDE4D^hi^ monocytes give rise to profibrotic SPP1^hi^MoAMs in post-COVI D-19 RLA

To investigate the relationship between circulating monocytes and lung macrophages, we integrated and reclustered PBMC (n=18,857) and BAL (n=107,132) myeloid cells, identifying previously described HLA-DR^lo^PDE4D^hi^ classical monocytes and SPP1^hi^MoAMs(Figure 3a-c, Supplementary Figure 6a).

**Figure 3.**
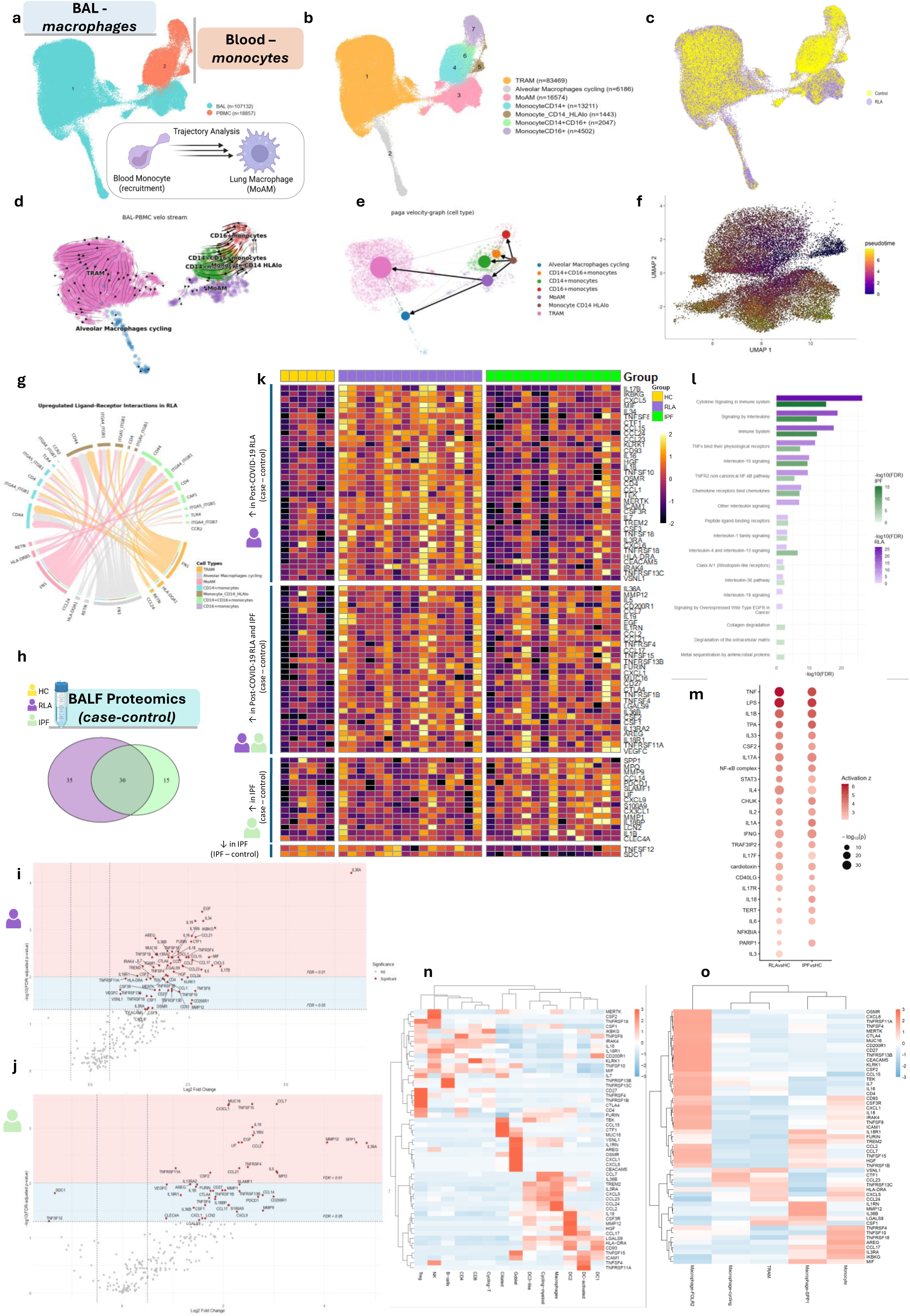
Blood monocyte – bronchoalveolar macrophage trajectory analysis and Bronchoalveolar Lavage Fluid Proteomics. **a–g** Integrated analysis of myeloid cells from BAL and PBMC compartments. UMAP of Harmony-integrated BAL and PBMC myeloid cells, labelled by sample type (**a**), cell type (**b**; cell numbers in parentheses), and disease (**c**; Healthy Control = yellow; RLA = purple;). NB The plotting order of cells in (**c**) was randomly shuffled to improve visual clarity. **d** RNA velocity vector fields projected onto the UMAP embedding and coloured by latent time, demonstrating velocity vectors pointing from HLA-DR^lo^PDE4D^hi^ classical monocytes to profibrotic SPP1+ MoAMs **e** Partition-based graph abstraction (PAGA) velocity graph showing directed connectivity between HLA-DR^lo^PDE4D^hi^ classical monocytes and MoAMs **f** Pseudotime trajectories across integrated blood CD14+ monocytes and BAL MoAMs, coloured by pseudotime **g** Upregulated ligand–receptor signalling interactions in post-COVID-19 RLA using CellChat. BAL macrophages were defined as signalling senders and circulating monocytes as receivers. The chord diagram illustrates significantly enriched signalling pathways in RLA compared with healthy controls (log₂ fold change > 0.2; FDR < 0.05). Interactions are grouped by cell type and colour-coded to indicate the direction and source of signalling, with ribbons representing ligand–receptor pairs contributing to enhanced macrophage–monocyte communication in RLA. **h** Bronchoalveolar lavage fluid (BALF) proteomics across disease groups. Area-proportional Venn diagram showing the number of significantly differentially expressed proteins identified when comparing cases versus controls in the RLA and IPF cohorts using Alamar NULISAseq. Shared (n = 30) and disease-specific proteins (n = 35 unique to RLA; n = 15 unique to IPF) are shown. **i,j** Differential protein abundance in bronchoalveolar lavage fluid (BALF) from RLA (**i**) and IPF (**j**) patients compared with healthy controls. Volcano plot showing log₂ fold change and statistical significance for proteins measured by Alamar NULISAseq. Differential expression was assessed using linear regression (lm) with Benjamini–Hochberg (BH) correction for multiple testing. Proteins meeting significance thresholds are highlighted (FDR < 0.01, pink; FDR < 0.05, blue). **k** Heatmap showing relative abundance of selected BALF proteins across healthy controls (HC), RLA, and IPF samples measured by Alamar NULISAseq. Data are shown at the patient level, with each column representing an individual patient and each row a protein. Proteins are organised into sections based on differential abundance patterns: increased in post-COVID-19 RLA compared with controls, increased in both post-COVID-19 RLA and IPF compared with controls, increased in IPF compared with controls and decreased in IPF compared with controls. Proteins are ordered by log₂ fold change in RLA (first and second sections) and IPF (third and fourth sections). Values are row-normali sed and displayed as z-scores, with samples grouped by disease status. **l** Reactome pathway enrichment of BALF proteins enriched in post-COVID-19 RLA and IPA, compared to healthy controls. Significantly increased proteins (FDR < 0.05; log₂FC > 0.5; linear regression with Benjamini–Hochberg correction) from RLA vs healthy controls and IPF vs healthy controls were analysed for pathway enrichment using g:Profiler2 (Reactome; with FDR correction). Bar plots show the top 15 enriched Reactome pathways per comparison, plotted as −log₁₀(FDR). **m** Ingenuity Pathway Analysis (IPA) upstream regulator analysis of BALF proteomic signatures. Proteins significantly differentially abundant (FDR < 0.05) in RLA versus healthy controls and IPF versus healthy controls (Alamar NULISAseq) were used as input. Only regulators predicted to be activated (positive activation z-score; | z| ≥ 2 with nominal p < 0.05) are shown, ordered by activation z-score in RLA. Dot colour denotes activation z-score and dot size denotes significance (−log₁₀(p)). **n–o** Cellular sources of differentially abundant BALF proteins in RLA mapped to BAL single-cell transcriptomic data, all BAL cells (**n**) and BAL myeloid (**o**). Proteins significantly differentially abundant in RLA versus healthy controls (FDR < 0.05; | log₂FC| > 0.5; Alamar NULISAseq) were mapped to gene expression in the BAL scRNA-seq atlas. Heatmaps show average log-normalised expression per cell type displayed as row-normali sed z-scores with hierarchical clustering (Ward’s method).

RNA velocity and partition-based graph abstraction (PAGA) analysis suggested directional trajectories from HLA-DR^lo^PDE4D^hi^ classical monocyte toward MoAMs, but not toward tissue resident alveolar macrophages (TRAMs) (Figure 3d, e), in post-COVID-19 RLA, but not in healthy controls (Supplementary Figure 6c). These monocytes exhibited the highest proportion of unspliced transcripts, consistent with a progenitor-like transcriptional state (Supplementary Figure 6b). Pseudotime analysis confirmed this trajectory identifying the HLA-DR^lo^PDE4D^hi^ classical monocytes as the root population, using an unbiased, bootstrapped connectivity approach (Figure 3f, Supplementary Figure 6d, e). Pseudotime further demonstrated a transcriptional cascade consistent with macrophage differentiation, described in fibrotic lung disease^12^. Early markers (CCR2, PDE4D, PELI2) decreased, whereas fibrotic effectors (SPP1, CCL2, MMP9) peaked at intermediate stages. Late macrophage markers (CD68, MERTK, CCL18) increased progressively, suggesting differentiation toward a profibrotic MoAM phenotype (Supplementary Figure 6f,g).

### Lung macrophages promote recruitment and profibrotic priming of circulating monocytes

To investigate cell–cell communication across the blood and lung myeloid compartments, we applied CellChat and NicheNet to integrated PBMC-BAL myeloid data. I n post-COVI D-19 RLA, MoAMs emerged as a central signalling hub, with increased outgoing and incoming signalling across inflammation- and fibrosis-associated pathways, including FN1, TNF, TGF-β and MHC-I (Supplementary Figure 7a, b). Cycling alveolar macrophages participated as both senders and receivers Supplementary Figure 7a,b), of fibrosis-associated (FN1–ITGA5) and chemotaxis (CCL23–CCR1/CCR2) pathways, with greater interaction strength observed in post-COVID-19 RLA compared with controls. Furthermore, the HLA-DR^lo^PDE4D^hi^ classical monocytes were prominent receivers of signalling from alveolar macrophages (Figure 3h), including pathway such as FN1–ITGAV, CD99-CD99L2, RETN–TLR4, TGFβ–TGFBR1 and CSF1–CSF1R (Figure 3g, Supplementary Figure 7d-g), suggesting monocyte-to-MoAM differentiation in the lung.

We identified a *CCL24–CCR2* chemokine axis linking lung macrophages to blood monocytes in RLA (Supplementary Figure 7c), alongside established (*CCL2–CCR2*, *CCL3/CCL23–*CCR1) recruitment signals. This *CCL24–CCR2* axis was predominantly derived from TRAMs and cycling alveolar macrophages and targeted CD14⁺ and CD14⁺CD16⁺ monocytes, suggesting selective monocyte trafficking to the fibrotic milieu.

NicheNet analysis also suggested that macrophage-derived cytokines and chemokines (TGFβ1, TNF, IL1B, CCL2, CCL3, CCL4, CCL5, CCL24) may regulate monocyte transcriptional programmes associated with inflammation, antigen presentation and fibrosis, with substantial overlap with genes upregulated along the monocyte-to-MoAM differentiation trajectory (Supplementary Figure 7h).

Together, these findings support a lung-blood signalling axis in which macrophage-derived cues recruit and reprogramme circulating monocytes toward profibrotic MoAM states.

### Proteomic profiling of bronchoalveolar lavage fluid reveals fibroinflammatory signalling networks in post-COVID-19 RLA

Proteomic profiling of BALF from patients with post-COVID-19 RLA (n=16), IPF (n=15) and healthy controls (n=6) revealed distinct signatures (Figure 4h-m), with principal component analysis showing clear separation across disease groups (Supplementary Figure 8a).

Comparison of differentially abundant proteins compared with healthy controls revealed both shared and disease-specific signatures; 30 proteins were significantly increased in both post-COVID-19 RLA and IPF, whereas 35 and 15 proteins were uniquely enriched in post-COVI D-19 RLA and IPF respectively (Figure 4h). These patterns were consistent across individual samples (Figure 4k).

**Figure 4.**
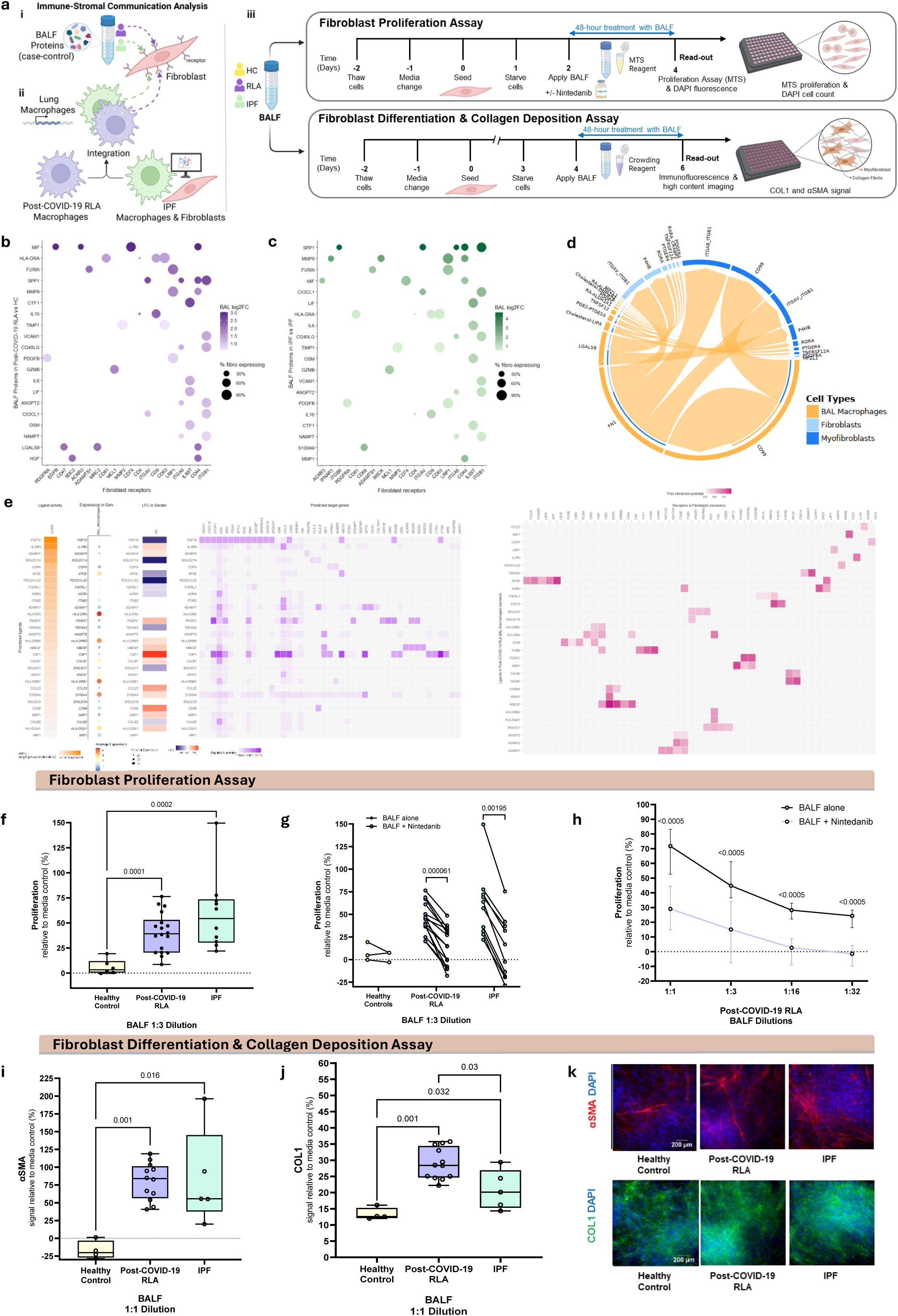
Immune-Stromal Cross Talk and Bronchoalveolar Lavage Fluid (BALF)- induced fibroblast proliferation, differentiation and collagen deposition. **a** Overview of the experimental design and immune–stromal communication analysis. **i**, Significantly differentially abundant bronchoalveolar lavage fluid (BALF) proteins from case–control comparisons with healthy controls (HC) in post-COVID-19 RLA and IPF were analysed to identify ligands with potential activity on fibroblast receptors using NicheNet. **ii**, Integrati on of single-cell macrophage transcriptional profiles from RLA (this study) with published IPF macrophage and fibroblast datasets to predict macrophage–fibroblast signalling interactions using NicheNet. **iii**, Functional validation using BALF stimulation assays in primary fibroblasts. Fibroblast proliferation was assessed after 48-hour BALF exposure using MTS assays and DAPI-based cell counts, with or without nintedanib treatment. Fibroblast differentiation and collagen deposition were assessed by immunofluorescence imaging of COL1 and αSMA following 48-hour BALF exposure. **b–c** Prioritisation of BALF ligands and cognate fibroblast receptors in post-COVID-19 RLA (**b**) and IPF (**c**) using the NicheNet ligand–receptor network. Significantly increased BALF proteins (Alamar NULISAseq) were treated as candidate ligands and linked to receptors expressed in fibroblasts. Dot colour indicates BALF protein log₂ fold change and dot size indicates the fraction of fibroblasts expressing the receptor. Ligands are ordered by their maximum ligand–receptor score (BALF log₂FC × fibroblast receptor expression × fraction of fibroblasts expressing the receptor), and receptors are ordered by the number of ligands predicted to signal through them, with ties resolved by score. **d** Macrophage–fibroblast signalling interactions in post-COVID-19 RLA inferred using CellChat. Single-cell transcriptomic data were analysed to predict ligand–receptor signalling from macrophages to fibroblasts. The chord diagram shows predicted signalling pathways, with ribbon width indicating interaction strength and colours denoting the signalling cell population. **e** NicheNet analysis predicting macrophage-derived ligands regulating fibroblast transcriptional responses in post-COVID-19 RLA. **Left**, Ligand-receptor interactions between BAL macrophages (senders) and fibroblasts (receivers) inferred using NicheNet, integrating post-COVID-19 BAL myeloid data (this study) with IPF dataset. Dot plot shows macrophage ligand expression (dot size) and log₂ fold change (LFC; colour intensity), ranked by regulatory potential (AUPR score; area under the preci sion-recall curve). **Middle**, heatmap showing predicted ligand–target gene regulatory potential. **Right**, Heatmap of ligand-receptor interactions between BAL macrophage ligands (y-axis) and predicted fibroblast receptors (x-axis), inferred using NicheNet. Colour intensity reflects predicted interaction strength. **f–h** Fibroblast proliferation following exposure to BALF with and without nintedanib measured by MTS assay. Proliferation was quantified relative to media control (Plasmax™ with 0.1% FBS; dotted line, y = 0) **f** Proliferation induced by BALF at a 1:3 dilution from healthy controls (HC, n = 3), post-COVID-19 RLA (n = 15), and IPF patients (n = 10). Each data point represents the mean of 3-4 technical replicates per biological repeat (patient). Biological repeats are shown with median ± IQR. Mann-Whitney U test; *p*-values indicated. **g**, Effect of nintedanib on BALF-induced fibroblast proliferation. Each line represents BALF from an individual patient; coloured circles indicate BALF alone and white circles indicate BALF plus nintedanib. Wilcoxon matched-pairs signed rank test, multiple testing correction was applied using the two-stage linear step-up procedure of Benjamini, Krieger, and Yekutieli to control the false-discovery rate at 5%. P-values are indicated. **h**, Serial dilution analysis of post-COVID-19 RLA BALF showing the effect size of BALF-induced fibroblast proliferation across increasing dilutions, with and without nintedanib. Coloured lines represent BALF with nintedanib; black lines represent BALF alone. Wilcoxon matched-pairs signed-rank test, adjusted with the Benjamini, Krieger, and Yekutieli procedure to control the false-discovery rate at 5%. P-values are indicated. **i–k** Fibroblast differentiation and collagen deposition following BALF exposure. Primary fibroblasts were treated with BALF from healthy controls (HC, n = 4), post-COVID-19 RLA (n = 12), or IPF patients (n = 5) at a 1:1 dilution. **i**, αSMA signal and **j**, collagen I (COL1) signal quantified relative to Plasmax™ 0.1% FBS macromolecular crowding medium control (dotted line, y = 0). Each data point represents the mean of 3–4 technical replicates per biological sample; biological repeats are shown with median ± IQR. Mann–Whitney test, p values indicated. **k**, Representative immunofluorescence images of fibroblasts treated with BALF from HC, post-COVID-19 RLA, or IPF showing αSMA (red), COL1 (green), and DAPI (blue) staining. Scale bars, 200 μm.

Proteins uniquely increased in post-COVID-19 RLA included inflammatory mediators such as IL17B, MIF, TREM2 and IKBKG, whereas IPF was characterized by enrichment of markers associated with extracellular matrix remodelling and neutrophilic inflammation, including SPP1 (osteopontin), MMP7, MMP9, IL1B, MPO and S100A9. A subset of proteins were increased in both conditions, including inflammatory mediators (IL36A, TNFRSF1B) and chemokines involved in immune recruitment (e.g. CCL2), LGALS9, an immunoregulatory galectin linked to fibrosis, as well as mediators associated with epithelial injury and tissue remodelling (e.g. MMP12, EGF, AREG, FURIN, MUC16/CA125) (Figure 4i, j; Supplementary Figure 8b).

Pathway analysis showed enrichment of immune and cytokine signalling in both conditions, whereas extracellular matrix remodelling and collagen degradation pathways were more prominent in IPF (Figure 4l, Supplementary Figure 8c, d).

Upstream regulator analysis predicted activation of inflammatory signalling pathways in post-COVID-19 RLA and IPF, including TNF, IL1B, IL6 and NF-κB pathway components (Figure 4m).

To infer the cellular sources of these mediators, proteins significantly increased in post-COVI D-19 RLA were mapped onto gene expression profiles from the BAL. Many differentially abundant BALF proteins, including CCL2, IL1B, CSF1, and LGALS9, were predominantly associated with airway macrophage populations (Figure 4n, o).

Together, these findings suggest that airway macrophages contribute to inflammatory and chemotactic mediators detected in BALF, however other cells, including those not captured by BAL (epithelial, endothelial and stromal cells) may influence the pool of soluble mediators.

### Immune–stromal communication links BALF-derived mediators to lung fibroblasts in post-COVI D-19 RLA and IPF

To investigate whether soluble mediators in BALF and macrophage-derived signals influence fibroblast activation and because stromal cells (fibroblasts) were not captured in our BAL dataset, we integrated our BAL macrophage single-cell data with a published IPF dataset that included macrophages, fibroblasts and myofibroblasts^19^ (Supplementary Figure 9a). This integration enabled analysis of two complementary signalling axes: (i) interactions between BALF-derived ligands and fibroblast receptors, and (ii) lung macrophage–fibroblast cross-talk (Figure 4a).

NicheNet was used to predict ligand–receptor interactions between BALF proteins enriched in post-COVID-19 RLA and IPF and lung fibroblast receptors (Figure 4b,c). Several inflammatory and profibrotic ligands detected in BALF were predicted to engage fibroblast receptors in both conditions, including SPP1, PDGFB, IL6, MIF and CX3CL1. These interactions corresponded to signalling pathways associated with fibroblast activation and migration, including integrin/CD44 signalling (SPP1), PDGF receptor signalling (PDGFB), IL6-IL6R signalling, and MIF–CD74/CXCR4 signalling. Many ligand–receptor interactions were shared between post-COVID-19 RLA and IPF, although the overall signalling profiles differed, indicating partially overlapping but distinct fibroblast activation programmes. Together, these findings suggest that soluble mediators present in the BALF have the potential to directly stimulate fibroblast activation.

### Macrophage–fibroblast cross-talk reveals mitogenic and profibrotic programmes in post-COVI D-19 and IPF

To infer macrophage-fibroblast crosstalk, we applied NicheNet and CellChat, identifying a network of profibrotic and mitogenic signalling interactions (Figure 4d,e). We first applied this approach to IPF, to confirm that it recapitulated established macrophage–fibroblast signalling interactions (Supplementary Figure 9b-e).

Pooled BAL macrophages from post-COVID-19 RLA expressed mitogens (PDGFC, HBEGF) predicted to drive fibroblast proliferation and matrix production, alongside fibroinflammatory cues (TGFB1, TNF, CSF1) associated with fibroblast activation and myofibroblast differentiation (Figure 4e). CellChat further identified PDGFC, FN1, Galectin (LGALS9), and CD99 as key macrophage-derived ligands communicating with fibroblasts and myofibroblasts, including PDGFC–PDGFRA, FN1–integrin, Galectin–CD44, and CD99–PILRA signalling axes (Figure 4d).

Comparative analysis of macrophage–fibroblast signalling in RLA and IPF revealed both shared and disease-specific features (Supplementary Figure 9b,c). In post-COVID-19 RLA, macrophages preferentially engaged mitogenic (PDGFC–PDGFRA) and inflammatory (TNF) signalling pathways, whereas SPP1–integrin/CD44 signalling, prominent in I PF, was absent in post-COVID-19 RLA. Matrix-associated cues (FN1, LGALS9, CD99) were conserved across conditions, suggesting shared extracellular matrix remodelling programmes.

To further investigate how the fibrotic milieu may be maintained, we examined macrophage–myofibroblast and fibroblast–myofibroblast signalling interactions using NicheNet (Supplementary Figure 9f,g). Macrophages were predicted to signal directly to myofibroblasts through profibrotic ligands, including TGFB1 and PDGFC, while fibroblasts expressed additional profibrotic ligands such as TGFB1, BMP2 and FGF2, with extracellular matrix components (COL1A1, FN1). Together, these findings confirm the validity of our computational approach, suggesting that fibroblasts may sustain myofibroblast activation through feed-forward stromal signalling loops, potentially reinforcing fibrosis independently of immune input.

### BALF-induced fibroblast proliferation in post-COVID-19 RLA and IPF is attenuated by nintedanib

Next, we functionally investigated the fibroproliferative signals inferred from BALF protein and macrophage-fibroblast cross-talk analysis, using fibroblast bioassays to evaluate fibroblast proliferation, differentiation and collagen deposition (Figure 4a(iii)). BALF from healthy controls, patients with post-COVID-19 RLA and IPF (Extended Data Table 6), was concentrated, desalinated and added in serial dilutions to primary human lung fibroblasts (Supplementary Figure 10), that had been pre-incubated with or without nintedanib (a receptor tyrosine kinase (RTK) inhibitor, an FDA-approved antifibrotic). Proliferation was assessed using the MTS assay and validated using DAPI cell counts (Supplementary Figure 11). The concentration of nintedanib was optimised using a PDGF-BB concentration-response curve (Supplementary Figure 12). BALF from post-COVID-19 RLA significantly increased fibroblast proliferation compared with healthy control BALF, albeit to a lesser extent than IPF BALF (median increase 39.1% vs. 68.8% at 1:3 dilution respectively; *p*<0.001) (Figure 4f). Nintedanib significantly attenuated BALF-induced fibroblast proliferation across all dilutions (Figure 4f–g; Supplementary Figure 14), with a greater inhibitory effect size observed in IPF. Of note, healthy control BALF elicited modest fibroblast proliferation, consistent with previous reports^20,21^.

These findings suggest that fibroblast proliferation is partly dependent on RTK signalling (e.g., PDGF, FGF), with a greater effect in IPF than in post-COVID-19 RLA.

### BALF from post-COVID-19 RLA and IPF promotes fibroblast differentiation and collagen deposition

We next assessed the impact of BALF on fibroblast-to-myofibroblast differentiation and collagen deposition, using αSMA expression and collagen I (immunofluorescence). Compared with healthy controls, BALF from post-COVID-19 RLA significantly increased fibroblast αSMA and collagen I expression (Figure 4i-k). Similar effects were also observed with IPF BALF. Although collagen I deposition appeared higher in the post-COVID-19 RLA group, this difference may reflect the smaller number of IPF BALF samples. Across all conditions cell counts (DAPI) (Supplementary Figure 13f) and viability (LDH levels) Supplementary Figure 13g), were maintained, confirming confluent monolayer integrity.

These findings provide functional evidence linking soluble mediators in post-COVID-19 RLA and IPF BALF to fibroblast proliferation, differentiation and collagen deposition.

### Compartment-specific reprogramming of T and NK cells in post-COVID-19 lung disease

We recently demonstrated that the lymphoid compartment may contribute to the immunopathogenesis of post-COVID-19 radiological subphenotypes^22^. To define disease-specific lymphocyte remodelling in post-COVID-19 RLA, we subclustered BAL and peripheral blood T/NK cells (Figure 5a-i). In the lung, NK cells were significantly depleted (Figure 5b, d). MAIT cells were reduced in both lung and blood, whereas Tregs and cytotoxic CD8⁺ZEB2⁺ T cells were expanded in the circulation (Figure 5g, i).

**Figure 5.**
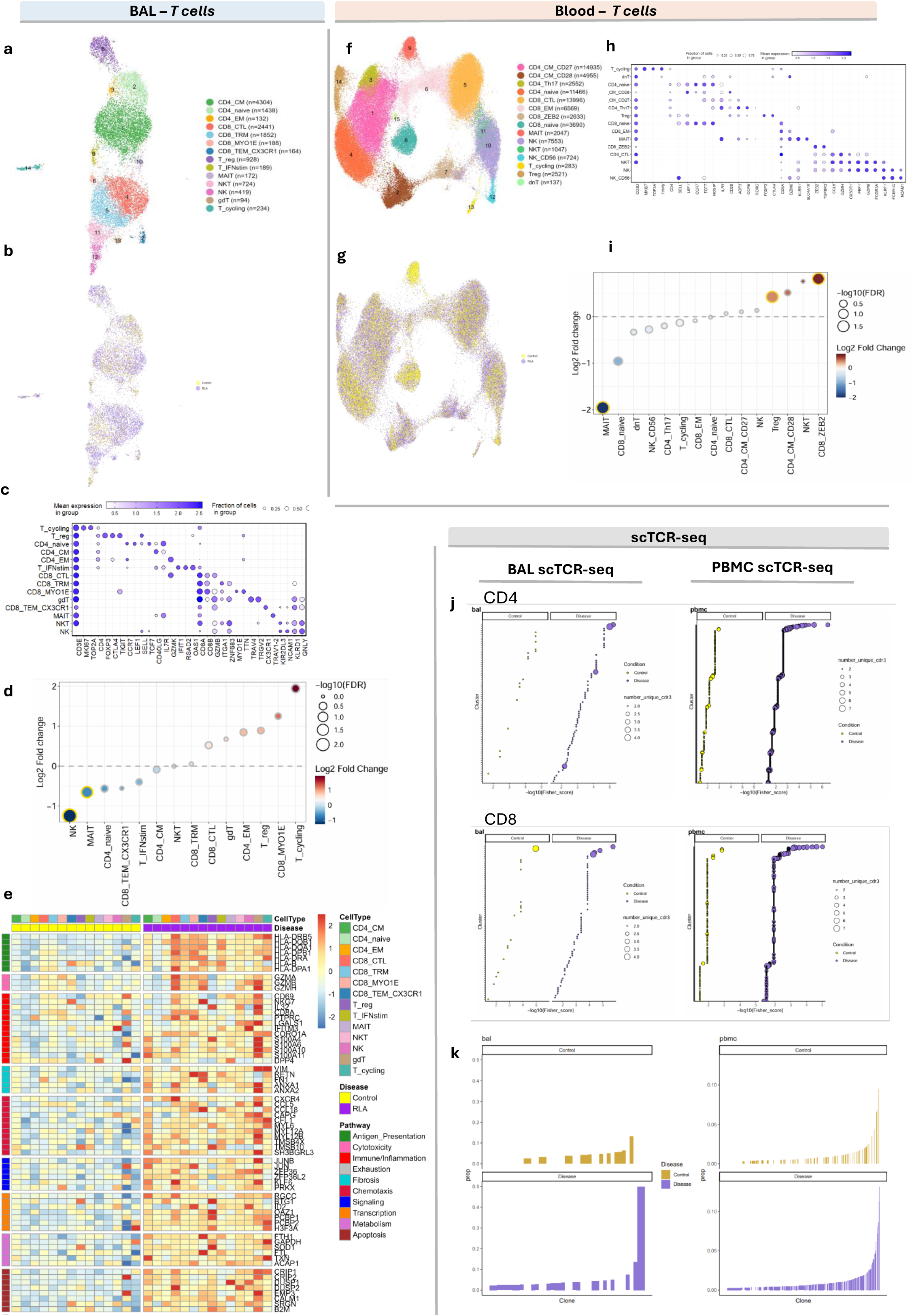
Single-cell and clonal analysis of T and NK cells in bronchoalveolar and blood compartments in post-COVID-19 RLA. **a–i** Single-cell transcriptomic analysis of T and NK cells in BAL (**a-e**) and blood (**f-i**) UMAP embedding of T and NK cells annotated by cell subtype (**a,f**) and (**b,g**) disease group (post-COVID-19 RLA = purple; Healthy Control = yellow). Cell plotting order was randomized between conditions to improve visualization **c,h**, Dot plot showing expression of canonical marker genes across T and NK cell subsets; colour indicates mean expression and dot size indicates the fraction of cells expressi ng each gene. **d,i**, Differential cell abundance analysis comparing post-COVID-19 RLA vs Healthy Control in BAL (**d**) and PBMC (**i**).Log₂ fold change (LFC) is shown, with red indicating increased and blue decreased abundance in RLA. Circle size reflects –log₁₀(FDR) from Wilcoxon rank-sum test with Bonferroni correction (FDR < 0.05, indicated by yellow border). Dashed line indicates LFC = 0. **e**, Heatmap of differentially expressed genes across BAL T and NK cell subsets, grouped by functional pathways. Genes ranked by FDR-adjusted p-value from Wilcoxon rank-sum test (per subject average expression), comparing each subset against all others. Gene expression values represent Z-score normalized log-normalized counts, where each gene’s expression is mean-centred and scaled across all samples. The colour scale reflects row-wise Z-scored expression, with red indicating upregulation and blue indicating downregulation in RLA relative to controls. DEGs were identified using Seurat’s FindMarkers function (Wilcoxon test; log₂ fold change > 0.25; FDR < 0.05; Bonferroni correction), excluding ribosomal genes. Top annotation indicates T cell type and disease group (Healthy Control = yellow, post-COVID-19 RLA = purple); right annotation categorizes DEGs by pathway. **j-k** scTCR seq analysis in BAL (left) and blood (right) **j** TCR repertoire analysis of CD4⁺ (top) and CD8⁺ (bottom) T cells in BAL (left) and PBMC (right) using GLIPH2 on each compartment separately. Each point represents a TCR cluster; point size indicates the number of unique CDR3 sequences per cluster. The x-axis denotes –log₁₀(Fisher score), indicating enrichment in RLA (purple) or Control (yellow). Clusters are ranked on the y-axis by Fisher score. **k** TCR clone frequency distribution in BAL (left) and PBMC (right) from post-COVID-19 RLA (purple) and Control (yellow) samples. The x-axis ranks individual clones by abundance, and the y-axis shows the proportion of total TCR sequences occupied by each clone (prop). Therefore each bar represents unique clones. RLA = post-COVID-19 Residual Lung Abnormalities, TRAM = tissue-resident alveolar macrophages.

In the lung differential gene expression analysis, revealed widespread transcriptional perturbation in post-COVID-19 RLA (Figure 5e). Multiple lung T-cell subsets, including CD4 effector-memory, CD8 tissue-resident memory, MAIT, and γδ T cells, exhibited upregulated cytotoxic effector genes such as *GZMB, GNLY*, and *PRF1*, consistent with enhanced activation. Several populations, including CD8+TRM, γδ T cells, IFN-stimulated T cells, and Tregs, also showed increased expression of MHC class II genes (*HLA-DRA*, *HLA-DRB1*, *HLA-DPB1*, *HLA-DQA1*, *HLA-DQB1*, *HLA-DRB5*), together with stress- and activation-associated genes including *DUSP1, ZFP36, JUNB,* and *BTG1,* suggesting a locally activated and transcriptionally remodelled bronchoalveolar T-cell compartment.

In peripheral blood, transcriptional changes were also evident across T cell multiple subsets, including signatures consistent with antigen presentation, immune activation, and apoptosis (Supplementary Figure 15e-g). CD8+ZEB2 cells expressed genes associated with cytotoxicity and terminal effector differentiation, consistent with sustained systemic immune activation.

### NK and MAIT cell depletion and transcriptional reprogramming in post-COVID-19 RLA lungs

NK cells were significantly reduced in the BAL of patients with post-COVID-19 RLA compared with healthy controls (Figure 5b,d). Residual NK cells exhibited transcriptional features of activation and cellular stress, including increased expression of cytotoxic genes (Figure 5e). In peripheral blood, NK cells showed increased expression of migration-associated genes (CXCR1, CXCR2, CX3CR1, CCL3 and CCL4) suggesting priming for tissue recruitment (Supplementary Figure 15e,f,g). However, their relative depletion in the BAL and reduced expression of homing receptors such as CCR1 and CCR7, suggesting altered trafficking or retention within the lung.

MAIT cells were significantly reduced in both the BAL and blood in post-COVID-19 RLA (Figure 5d,i). In the lung, MAIT cells exhibited increased expression of *TMSB4X* (encoding thymosin β4), a regulator of tissue repair and immune modulation, consistent with a transcriptional shift toward a reparative or regulatory phenotype. In peripheral blood, MAIT cells showed transcriptional features of activation and immune regulation, including *CD69, DPP4, HLA-B, INPP4B, GIMAP7* and *TRAF3IP3*, with downregulated chemokine receptors involved in inflammatory trafficking (*CX3CR1, CCR1, CCR2* and *CXCR4*) (Supplementary Figure 15e-g). Conversely, expression of the mucosal-homing receptor gene (*CCR6*) was increased, suggesting altered migratory behaviour and enhanced localisation to epithelial barrier sites. Together, these features indicate that MAIT cells in post-COVID-19 RLA may adopt an activated and dysregulated state, characterised by reduced abundance, and altered trafficking programmes.

### Oligoclonal T cell expansion in the lung contrasts with broader polyclonal expansion in the blood

To investigate T cell clonal dynamics in post-COVID-19 RLA, we performed TCR repertoire analysis of CD4⁺ and CD8⁺ T cells from BAL and PBMC samples(Figure 5h,i). TCR similarity was assessed using GLIPH2 clustering (Figure 5h), which identifies groups of TCRs with shared sequence motifs and potential antigen specificity, and clonal dominance was evaluated using clone frequency distributions (Figure 5i).

GLIPH2 clustering showed disease-enriched TCR clusters in both CD4+ and CD8+ T cells in BAL and PBMC compartments in post-COVID-19 RLA compared with healthy controls (Figure 5j, (Supplementary Figure 16c). In the blood, TCR repertoires were broadly polyclonal, with multiple clusters contributing at low frequencies. In contrast, BAL repertoires exhibited oligoclonal dominance, with a small number of highly expanded clones comprising a large fraction of the repertoire. Clone frequency distribution plots supported these findings (Figure 5k).

Integration of TCR repertoire with gene-expression data, showed that clonal expansion occurred across multiple T cell states (Supplementary Figure 16a, b). Clonal sharing between blood and BAL was minimal (Supplementary Figure 16d).

Together, these findings suggest compartment-specific T cell clonal dynamics in post-COVID-19 RLA, with oligoclonal T cell dominance in the lung and broader polyclonal responses in the circulation.

## Discussion

Our study presents a high-resolution, cross-compartment, multi-omic atlas of patients with post-COVID-19 RLA, proposing a tripartite model (Figure 6) of (i) Immune dysregulation involving myeloid alterations, whereby HLA-DR^lo^CD163^+^PDE4D^hi^ classical monocytes expand in peripheral blood, are recruited to the lung and differentiate into profibrotic SPP1⁺MoAMs, that are expanded in disease. In lymphoid populations, there is depletion of MAIT cells in both the lung and blood, decreased NK cells with oligoclonal T cell expansion in the lung, and expansion of regulatory and cytotoxic T cells in the blood; (ii) Immune-stromal cross-talk is mediated by alveolar myeloid cell-derived BALF proteins which engage fibroblast receptors and macrophage-fibroblast communication; (iii) Soluble mediators in BALF induce fibroblast proliferation, differentiation and collagen deposition, with proliferation attenuated by nintedanib *in vitro*. Together, these findings suggest that persistent fibroinflammatory immune–stromal remodelling in post-COVID-19 RLA is driven by a dysregulated monocyte–macrophage–fibroblast axis.

**Figure 6.**
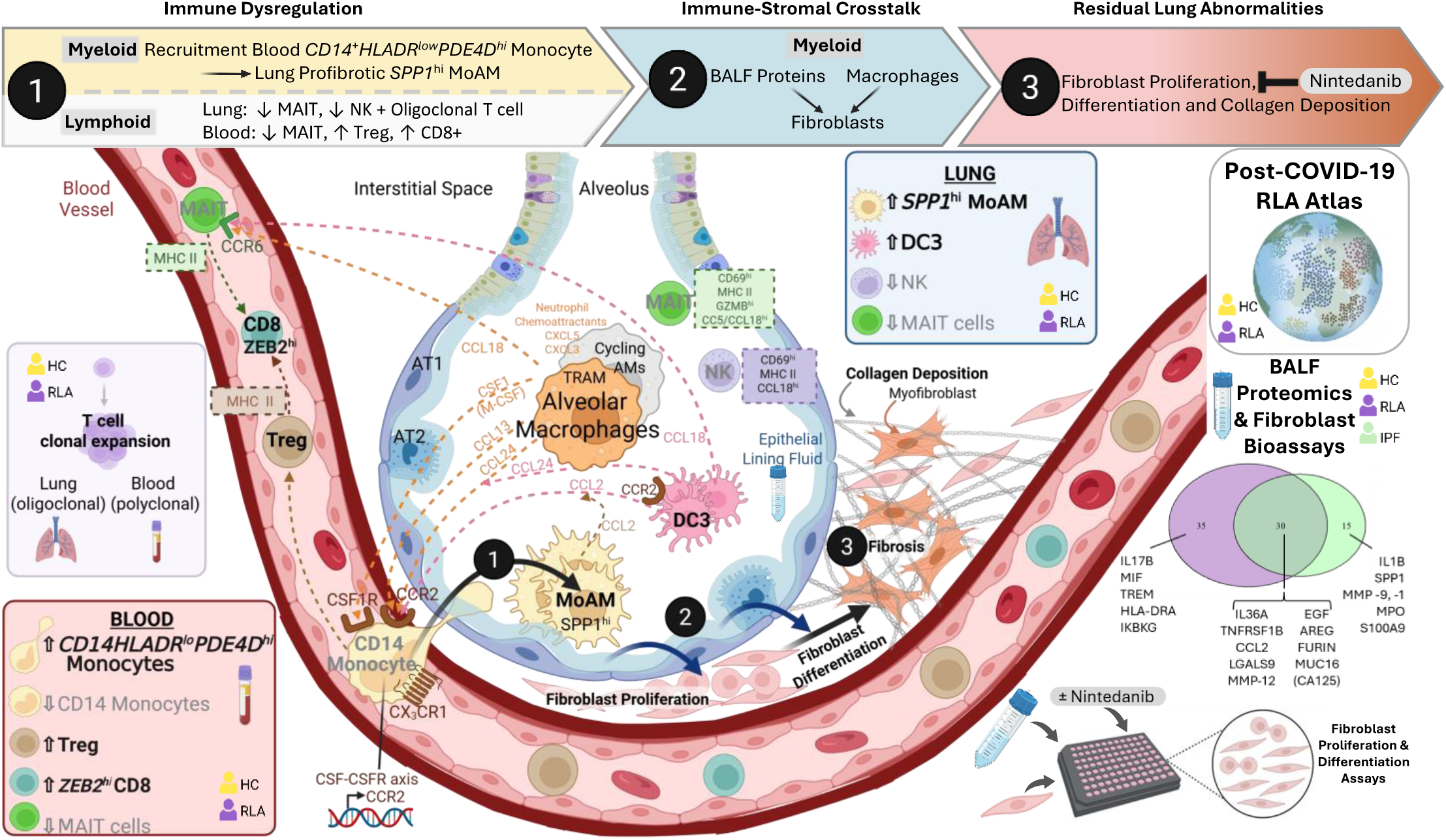
Proposed model of immune–stromal interactions driving post-COVID-19 residual lung abnormalities (RLA). A single-cell atlas of bronchoalveolar lavage (BAL) and peripheral blood mononuclear cells (PBMCs), combined with BALF proteomics and fibroblast functional bioassays, identifies mechanisms of lung remodelling in RLA. (1) Circulating CD14⁺ monocytes are recruited to the lung via CCL–CCR and CSF1–CSF1R signalling, where HLA-DR^lo^PDE4D^hi^ classical monocytes differentiate into profibrotic *SPP1*⁺ monocyte-derived alveolar macrophages (MoAMs). (2) Fibrogenic mediators produced by myeloid cells promote fibroblast proliferation and differentiation within the interstitial space. (3) Activated fibroblasts differentiate into myofibroblasts, driving collagen deposition and extracellular matrix remodelling that underlies post-COVID-19 RLA. RLA lungs exhibit increased MoAMs and DC3 cells with reduced NK and MAIT cells, while blood shows increased Tregs and CD8⁺ ZEB2^hi^ T cells. scTCR-seq reveals oligoclonal T cell expansion in the lung compared with polyclonal expansion in blood.

Our data supports a blood-lung myeloid axis. Prior reports identified an expanded and dysfunctional population of CD14⁺HLA-DR^lo^CD163^+^ monocytes in acute severe COVI D-19^8,9^, but not in post-COVID-19 lung disease (nor in IPF or healthy controls)^8^. SPP1^hi^ MoAMs have been described in acute COVID-19^4^ and post-COVID-19 lung disease^6^, where their abundance correlates with the severity of radiological fibrosis^6^. To our knowledge, our study is the first to detect these circulating monocytes in patients with post-COVID-19 RLA, and furthermore to implicate them as the cellular origin of the profibrotic SPP1+ MoAMs.

We also provide a comprehensive BALF proteomic analysis in post-COVID-19 RLA, and by integrating proteomics and single-cell data, we demonstrate that BALF mediators originate predominantly from airway myeloid populations and are predicted to engage fibroblast receptors. Furthermore, our bronchoalveolar proteomic characterisation in IPF using ultrasensitive Alamar NULISAseq technology represents a unique dataset, given that bronchoscopy is infrequently performed in IPF patient and proteomic data remain sparce^24^. The post-COVID-19 RLA BALF proteome partially overlaps with, but is distinct from IPF.

Shared mediators included factors associated with immune recruitment, epithelial injury and tissue remodelling (CCL2, MMP12, IL36A, EGF, AREG). Our findings align with prior reports in post-COVID-19 lungs, where monocyte-derived BALF CCL2 is linked to fibrosis severity^6^, and elevated EGF^6^ and AREG^25^ likely reflect ongoing epithelial repair and fibroproliferative signalling. Post-COVID-19 RLA BALF showed distinct enrichment of immune signalling mediators (IL17B, MIF, TREM2 pathways). In contrast, IPF BALF was uniquely enriched for markers of extracellular matrix turnover and neutrophilic inflammation (SPP1, MMP9, MPO, S100A9).

Our findings also support an immune-stromal axis in post-COVID-19 RLA, similar to IPF, as both myeloid-derived proteins and alveolar macrophages were predicted to engage fibroblast receptors. Ligand–receptor analysis identified multiple signalling axes (e.g., PDGF, IL6, MIF, integrin/CD44) capable of directly engaging fibroblasts, while macrophages expressed profibrotic ligands including PDGFC, FGF10, HBEGF and TGFB1. This landscape suggests a more dynamic and potentially reversible fibroproliferative niche in post-COVI D-19 lungs, in contrast to IPF where canonical SPP1–integrin signalling was dominant.

Functional bioassays demonstrated BALF-induced fibroblast proliferation, differentiation and collagen deposition. While BALF-induced fibroblast proliferation has been described in other fibrotic lung diseases^20,26^, to our knowledge, our study provides the first demonstration that nintedanib can attenuate this effect. The reduced inhibitory effect size of nintedanib in post-COVID-19 RLA compared with IPF, suggests that fibroproliferation in post-COVID-19 lungs may depend on a broader and less RTK-centric signalling network. Identified mitogens in BALF (e.g., EGF, AREG), signal via EGFR, which is relatively nintedanib-insensitive^27^, suggesting involvement of both RTK -dependent and -independent pathways.

Overall, post-COVID-19 RLA shares fibrotic mechanisms with IPF, but is distinguished by greater immune activation, signalling heterogeneity, and cellular plasticity. These features suggest a less consolidated and potentially reversible disease state. Although antifibrotic therapy likely modulates key stromal responses, the persistence of inflammatory signalling indicates that combination approaches incorporating immunomodulation may be required to restore tissue homeostasis.

Beyond the myeloid-stromal axis, we observed marked remodelling of lymphocyte populations. MAIT cells were significantly depleted in both lung and blood, in contrast to their peripheral egress^28^ and lung accumulation^29,30^ in acute severe COVID-19. Although MAIT cells repopulate during convalescence, they remain functionally impaired^31^, and in our cohort showed no correlation with time since acute infection (Supplementary Figure 15c). Given their role in promoting antifibrotic dendritic cell responses^32^, this depletion may represent a failure of immune regulation, supported by the expansion of DC3 cells in the post-COVID-19 lung, which are implicated in skin and lung fibrosis^33^. T cells exhibited a sustained, tissue-directed activation profile. BAL T cells upregulated cytotoxic mediators, antigen presentation pathways, and stress-response genes, consistent with a potentially tissue-damaging immune state. Circulating ZEB2⁺CD8⁺ T cells were expanded and enriched for cytotoxic and migratory programmes, suggesting ongoing antigen exposure. Regulatory T cells were increased in blood but displayed features of exhaustion in the lung, suggesting impaired local immunoregulation. TCR sequencing demonstrated compartmentalised clonal expansion with minimal overlap between blood and lung, supporting persistent, localised immune activation within the fibrotic niche. NK cells were depleted in BAL, despite evidence of migratory priming in circulation, suggesting impaired recruitment or retention. Residual NK cells were skewed towards a CD56^bright^CD16⁺ phenotype(Supplementary Figure 15a-b), contrasting with the CD56^dim^CD16^+^ phenotype observed in healthy lungs^34,35^. Similar depletion of NK cells has been observed in acute severe COVI D-19^36,37^ and I PF lungs^38,39^, and is associated with increased fibrosis in mouse models^38^. Together, these findings indicate that the lymphoid compartment in post-COVID-19 RLA is characterised by persistent cytotoxic T cell activation, impaired regulatory control, and loss of NK cell-mediated homeostasis, promoting chronic inflammation and potentially fibrotic remodelling.

Our study has some limitations. The modest cohort size may affect the generalisability of our findings. Sampling (BAL) and technical (single-cell genomics) constraints limited comprehensive cellular profiling, given the inherent challenges in detecting interstitial macrophage and neutrophils, respectively. Nevertheless, we were able to characterise the immune landscape at high resolution. Although computational communication and trajectory analyses are inherently predictive, our findings were supported using orthogonal approaches. Functional bioassays were restricted by BALF sample availability, however we were able to detect statistically significant differences between groups and dilutional effects were reproducible, supporting biological plausibility. Restricted BALF volumes precluded evaluation of nintedanib on fibroblast differentiation and collagen deposition, however we were able to demonstrate significant effects on fibroblast proliferation. IPF fibroblasts were used for bioinformatic analyses, whereas primary human lung fibroblasts from a healthy donor were used for bioassays. However, there is experimental evidence that differences between IPF and healthy fibroblasts may influence the magnitude, rather than the direction of observed effects^40^.

In summary, our findings offer mechanistic insight into the immunopathology of post-viral lung fibrosis and support further evaluation of potential combined immunomodulatory and antifibrotic strategies, for patients with persistent symptoms and radiological changes. Using post-COVID-19 RLA, as a human model of fibroinflammation, our cross-compartment, multi-omic integrated paired BAL and blood profiling, with single-cell and proteomic analyses, suggests a central monocyte-macrophage-fibroblast axis, and is supported by functional bioassays, with potential relevance across fibrotic lung diseases.

## Methods

### Study participants

The study cohort comprised participants of the <u>POST-COVID</u>-19 interstitial lung <u>D</u>iseas<u>E</u> (POSTCODE) study, a sub-study of the UK Interstitial Lung Disease (UK-ILD) Long COVID study, a prospective, multicentre evaluation of patients with suspected ILD following COVID-19^41^. Ethical approval was obtained from the North London Research Ethics Committee (13/LO/0900; 13/0186; LPR 143364, IRAS 297891, REC 21/HRA/3313, IRAS 297495, REC 21/SC/0180). Written informed consent was obtained from all participants. Participants were recruited from University College London Hospitals NHS Trust and Imperial College London NHS Trust (UK) between July 2020 and August 2022. Full eligibility criteria are outlined in Extended Data Table 3, Extended Data Table 4.

### Sample collection and cell isolation

BAL samples were obtained using flexible fibreoptic bronchoscopy into the right middle lobe or lingula. BALF was collected by aspiration and sent for microbiological testing and differential cell counts. Remaining BALF was immediately cooled to 4°C and processed within 30 minutes, filtered to remove debris and centrifuged to separate cells and supernatant. Supernatant was stored at −80 °C for downstream analyses, and cells were resuspended in PBS for viability assessment prior to single-cell capture using the Chromium Next GEM Single Cell 5⍰ platform (10x Genomics). 20,000 cells were loaded per channel.

Peripheral blood was collected in EDTA tubes and PBMCs isolated by Ficoll density-gradient centrifugation, as previously described^42^. PBMCs were cryopreserved until further processing.

### Single cell RNA sequencing and CITE-seq library preparation

PBMCs were thawed, pooled using stratified randomization across disease groups (four donors per pool) and stained with a TotalSeq-C antibody panel (BioLegend) according to the manufacturer’s instructions. Cells were then processed for single-cell capture using the Chromium Next GEM Single Cell 5⍰ platform with Feature Barcoding technology (10x Genomics). Two lanes of 25,000 cells were loaded per pool onto a 10X chip.

Single-cell RNA-seq libraries for BAL cells were generated using the Chromium Next GEM Single Cell 5⍰ v2 kit (10x Genomics). PBMC libraries were prepared using the Chromium 5⍰ V(D)J and Feature Barcoding kit. Libraries were sequenced on Illumina NovaSeq 6000 or NextSeq 2000 instruments.

### Single-cell RNAseq data processing

Raw sequencing data were processed using Cell Ranger (10x Genomics) and aligned to the GRCh38 reference genome. Downstream analyses were performed in R using Seurat. Low-quality cells and predicted doublets were removed based on transcript count, mitochondrial gene content and doublet detection algorithms. Data were normalised, integrated across donors and clustered using a Louvain algorithm

Pooled PBMC samples were demultiplexed using genotype-based assignment and verified using souporcell (v.2)^43^ and further filtered for doublets. Gene expression and antibody-derived tag counts were normalised and integrated using Harmony (v.1.2.3)^44^. Weighted nearest neighbour graphs were generated from gene expression and protein data.

Clusters were annotated using reference mapping^45^ and manual marker-based curation (Supplementary Methods).

### Cell composition, differential expression and gene signature analysis

Cell type proportions were compared across conditions using the Wilcoxon rank-sum test with Benjamini-Hochberg correction for multiple testing. Log2 fold changes were computed to quantify differences, and significance thresholds were set based on FDR-adjusted p-values.

Differential gene expression was assessed using Seurat with the Wilcoxon rank-sum test and Benjamini–Hochberg correction for multiple testing. Genes expressed in ≥25% of cells with adjusted *P*⍰<⍰0.05 were considered significant.

To assess generalisability of macrophage subset annotation, we performed gene signature over-representation analysis with curated gene sets from published single-cell studies of acute COVID-19 and IPF using Fisher’s exact test.

### Trajectory inference and cell-cell communication

To infer relationships between circulating monocytes and lung macrophages, PBMC and BAL myeloid compartments were integrated. RNA velocity, pseudotime analysis and partition-based graph abstraction were used to infer cellular trajectories^46-50^. Intercellular communication networks were predicted using CellChat^51^ and NicheNet^52^ to identify ligand–receptor interactions. To investigate interactions between BAL macrophages and the fibrotic niche (fibroblasts and myofibroblasts), we integrated our post-COVID-19 BAL myeloid dataset with a published 5’ single-cell dataset of dissociated lung tissue from IPF patients^19^ using Harmony. This integrated dataset only included macrophages, fibroblasts, and myofibroblasts. NicheNet^52^ and CellChat^51^ were then used to analyse macrophage-fibroblast communication.

### TCR repertoire analysis

T cell receptor V(D)J libraries were generated using the Chromium Single-Cell V(D)J platform (10x Genomics). Clonotypes were defined using shared V(D)J gene usage and CDR3 sequences and analysed using scRepertoire^53^ and STARTRAC^54^. TCR clustering and expansion analysis were performed using GLIPH2^54^, clustering CD4⁺ and CD8⁺ TCR sequences separately in PBMC and BAL.

### BALF Proteomics

Proteomics was performed on BALF samples from patients with post-COVID-19 RLA, IPF and healthy controls, using the Alamar Biosciences Inflammation 250 panel, using the Nucleic Acid Linked Immuno-Sandwich Assay sequencing (NULISA-seq) platform, according to the manufacturer’s protocol. Protein counts were normalised prior to principal component analysis. Differential protein abundance in BALF from patients with post-COVID-19 RLA and IPF was compared with healthy controls using linear regression modelling with Benjamini–Hochberg correction for multiple testing. Significantly differentially expressed proteins were analysed using Reactome pathway enrichment (g:Profiler2) and upstream regulator analysis (Ingenuity Pathway Analysis). Cellular source attribution analysis involved protein-gene mapping. NicheNet was used to infer candidate protein ligand–fibroblast receptor interactions.

### Fibroblast proliferation and differentiation bioassays

Primary human lung fibroblasts were cultured and exposed to concentrated and desalted BALF, to assess the effects of BALF on fibroblast proliferation and differentiation and collagen deposition. Fibroblast proliferation was assessed using MTS assays and DAPI cell counts, and Inhibition experiments were performed with nintedanib, a clinically approved antifibrotic agent for pulmonary fibrosis. Fibroblast differentiation experiments were performed under macromolecular crowding conditions and analysed using high-content imaging-based quantification of collagen I and αSMA immunostaining. Detailed experimental protocols are provided in the Extended Methods.

### Statistical analysis

Statistical analyses were performed in R (v4.4.0). Differential expression and cell composition analyses were conducted using Wilcoxon rank-sum tests with Benjamini–Hochberg correction^55^ for multiple testing. A significance threshold of *P*⍰<⍰0.05 was used unless otherwise stated. Investigators performing fibroblast bioassays were blinded to experimental condition.

## Supporting information

UK-ILD consortium

## Extended Data Figures with legends

**Supplementary Figure 1.**
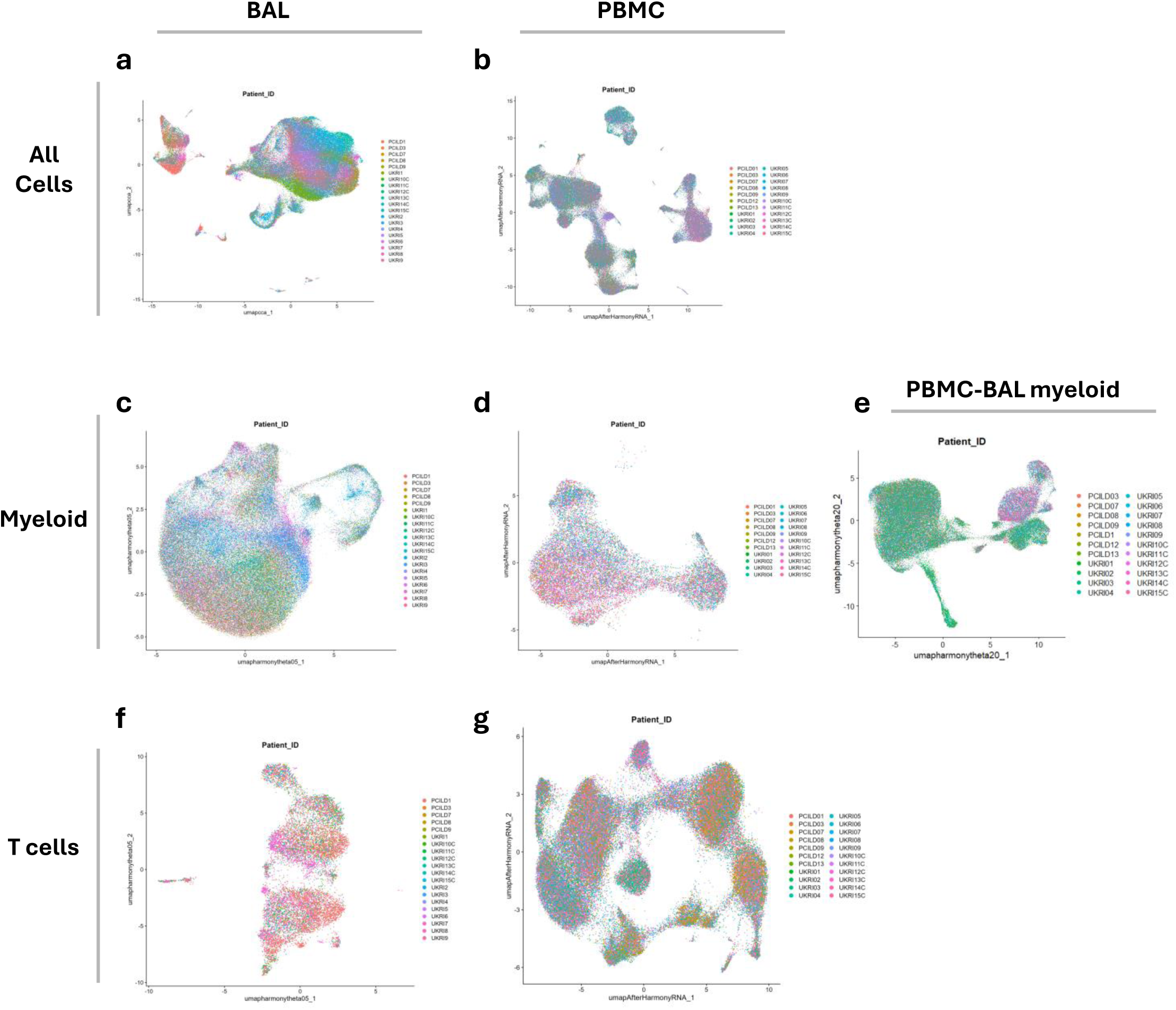
Patient-level UMAPs demonstrating integration across BAL and PBMC datasets. Each point represents a single cell coloured by originating patient. UMAPs are coloured by patient ID. Cells from different patients are well mixed across clusters, indicating effective integration and removal of patient-specific batch effects while preserving underlying cellular structure. **a** All bronchoalveolar lavage (BAL) cells **b** All peripheral blood mononuclear cells (PBMCs). **c** BAL myeloid cells. **d** PBMC myeloid cells. **e** integrated BAL–PBMC myeloid compartment. **f** BAL T cells. **g** PBMC T cells.

**Supplementary Figure 2.**
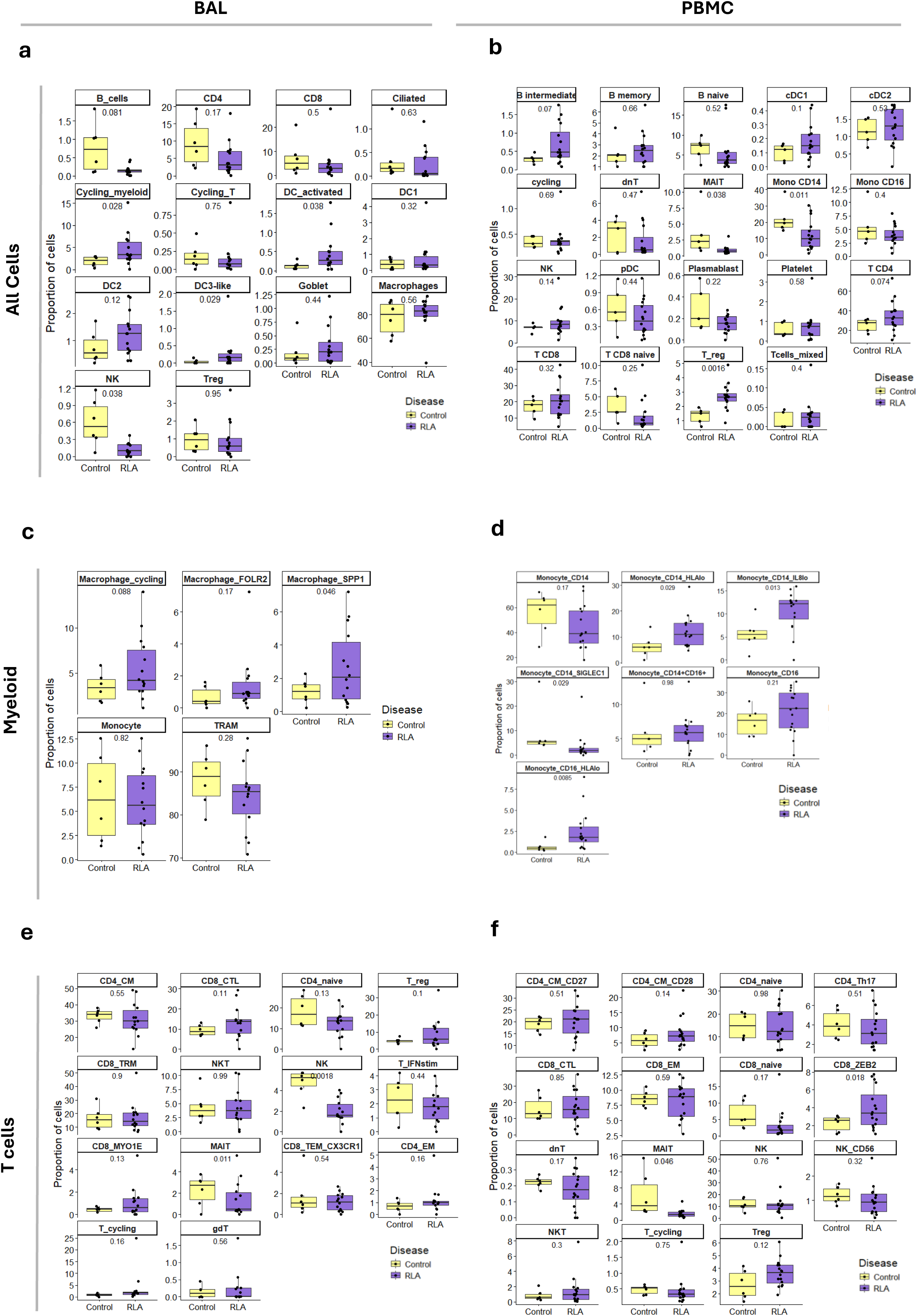
Patient-level cell type proportions across Control and RLA samples. Boxplots show the proportion of each myeloid cell type per patient, calculated as the percentage of total immune cells within each sample. Each point represents an individual patient. Boxes indicate the interquartile range with the median shown by the centre line; whiskers extend to 1.5× the interquartile range. Statistical comparisons between Control and RLA groups were performed using an unpaired two-sided t-test, and p-values were adjusted for multiple testing using the Benjamini–Hochberg (BH) false discovery rate correction. **a** All bronchoalveolar lavage (BAL) cells **b** All peripheral blood mononuclear cells (PBMCs). **c** BAL myeloid cells. **d** PBMC myeloid cells. **e** BAL T cells. **f** PBMC T cells.

**Supplementary Figure 3.**
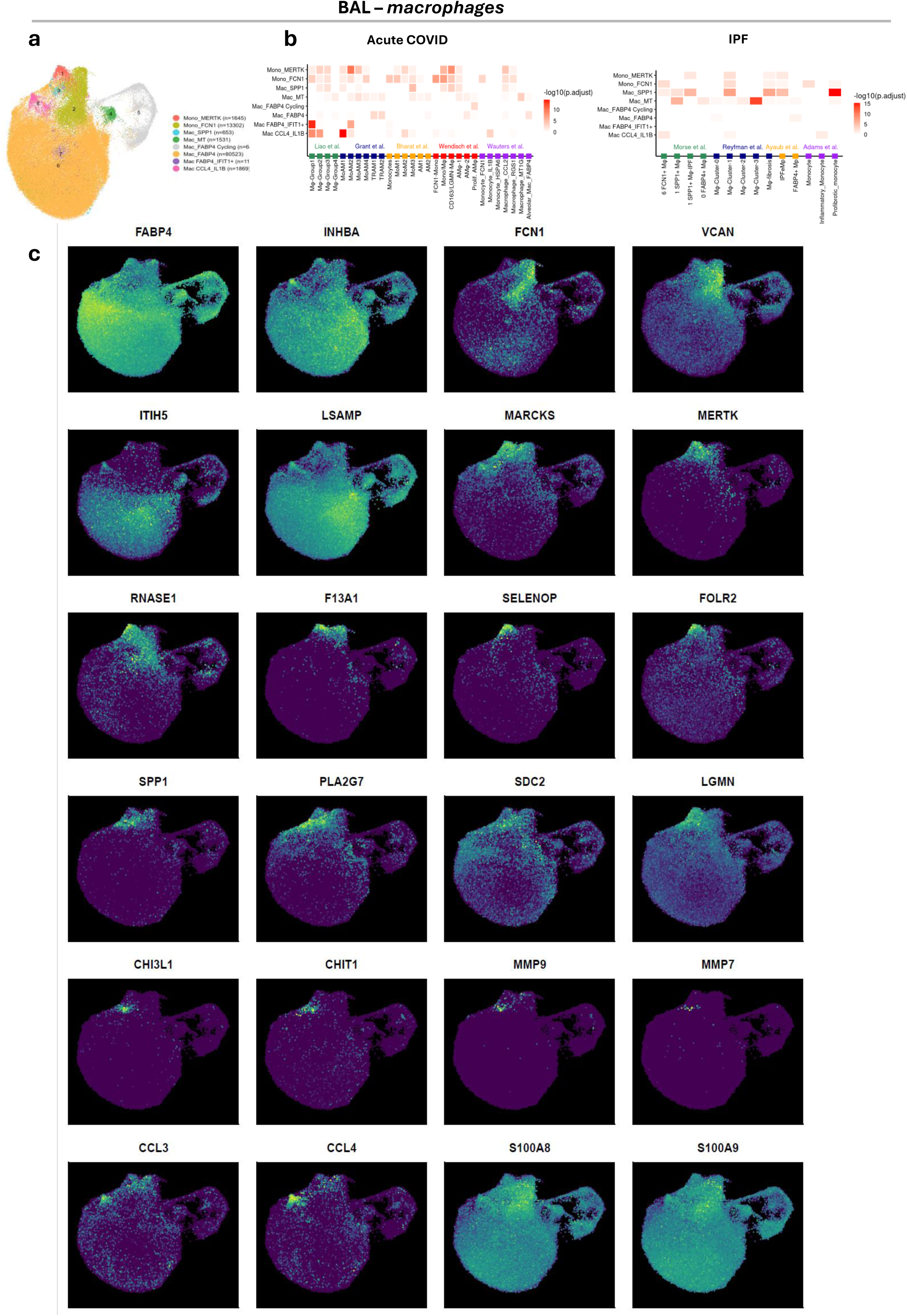
Single-cell analysis ofbronchoalveolar macrophages. **a** Uniform manifold approximation and projection (UMAP) visualization of annotated myeloid BAL cells labelled by major cell types with cell numbers shown in parentheses Mono= Monocyte; Mac = Macrophage **b** Heatmap showing –log₁₀ transformed adjusted p-values (one-sided Fisher’s exact test) for overlap between gene sets from BAL monocyte/macrophage clusters in RLA (y-axis) and published transcriptional signatures from acute COVID-19 (left) and IPF (right) datasets (x-axis). Colour intensity indicates statistical significance of overlap. **c** Feature plots show marker gene expression across BAL myeloid populations, including Macrophage_MERTK (MERTK, FOLR2), Monocytes (FCN1, VCAN), Profibrotic (SPP1, MMP9, MMP7, CHIT1, PLA2G7), Tissue residency (FABP4, INHBA, LSAMP), Interferon stimulation (IFIT1), and Inflammation (CCL3, CCL4).

**Supplementary Figure 4.**
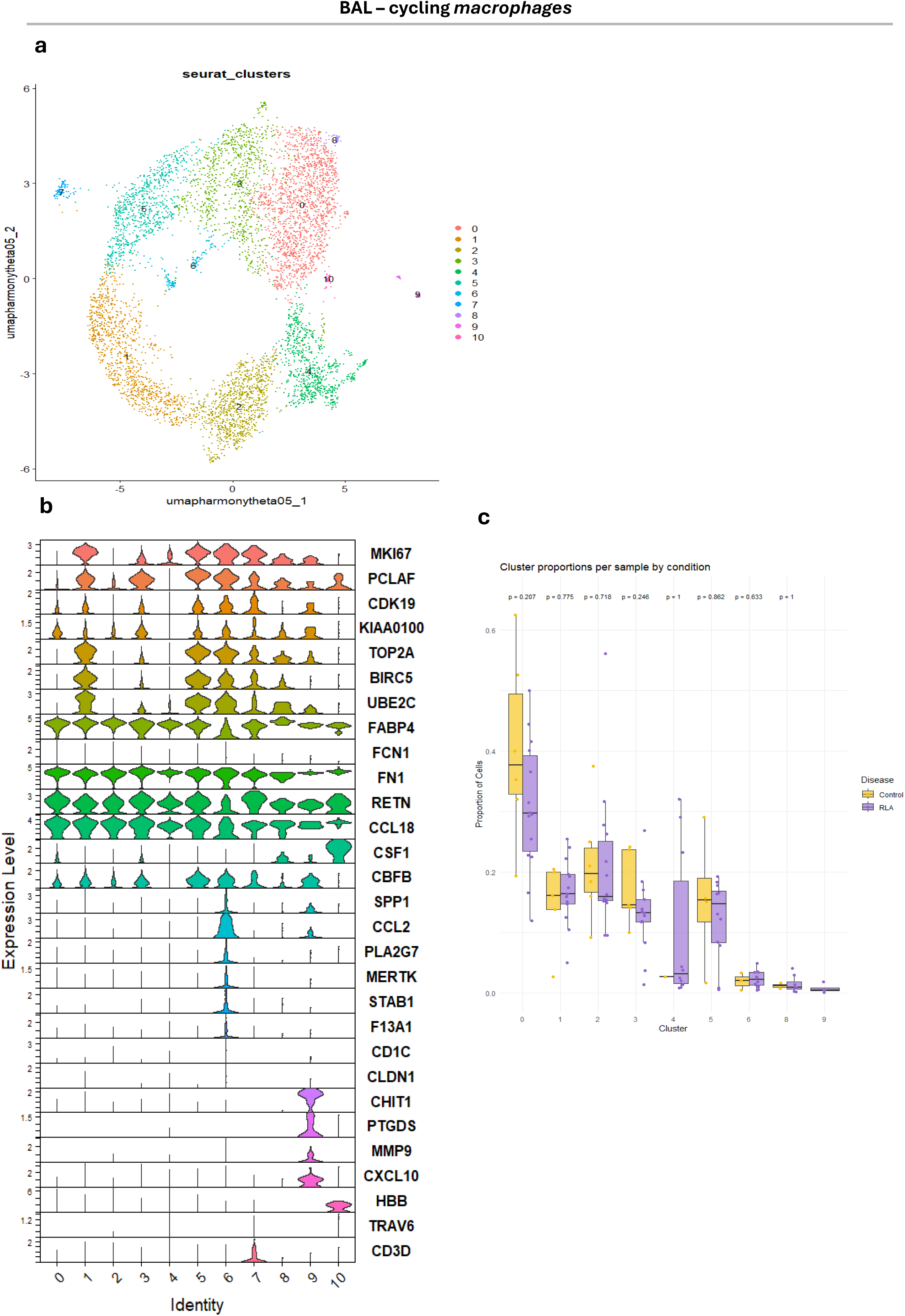
Single-cell analysis of cycling bronchoalveolar macrophages. **a** Uniform manifold approximation and projection (UMAP) visualization of cycling macrophages clustered by unsupervised graph-based clustering **b** Violin plots showing expression of proliferation (MKI67, PCLAF, CDK19), homeostatic (FABP4, FCN1), and profibrotic/remodelling (SPP1, MERTK, CCL2, FN1, RETN, PLA2G7) markers, alongside dendritic-like and contaminating population markers (HBB, TRAV6, CD3D). **c** Per-sample cluster proportions by condition (Control = yellow, RLA = purple); Wilcoxon rank-sum test shows no significant differences. One SPP1⁺ cycling macrophage cluster (cluster 9) was enriched in RLA and co-expressed fibrosis-associated genes (MMP9, CHIT1, PTGDS, CXCL10, CD1C), suggesting a disease-associated profibrotic population.

**Supplementary Figure 5.**
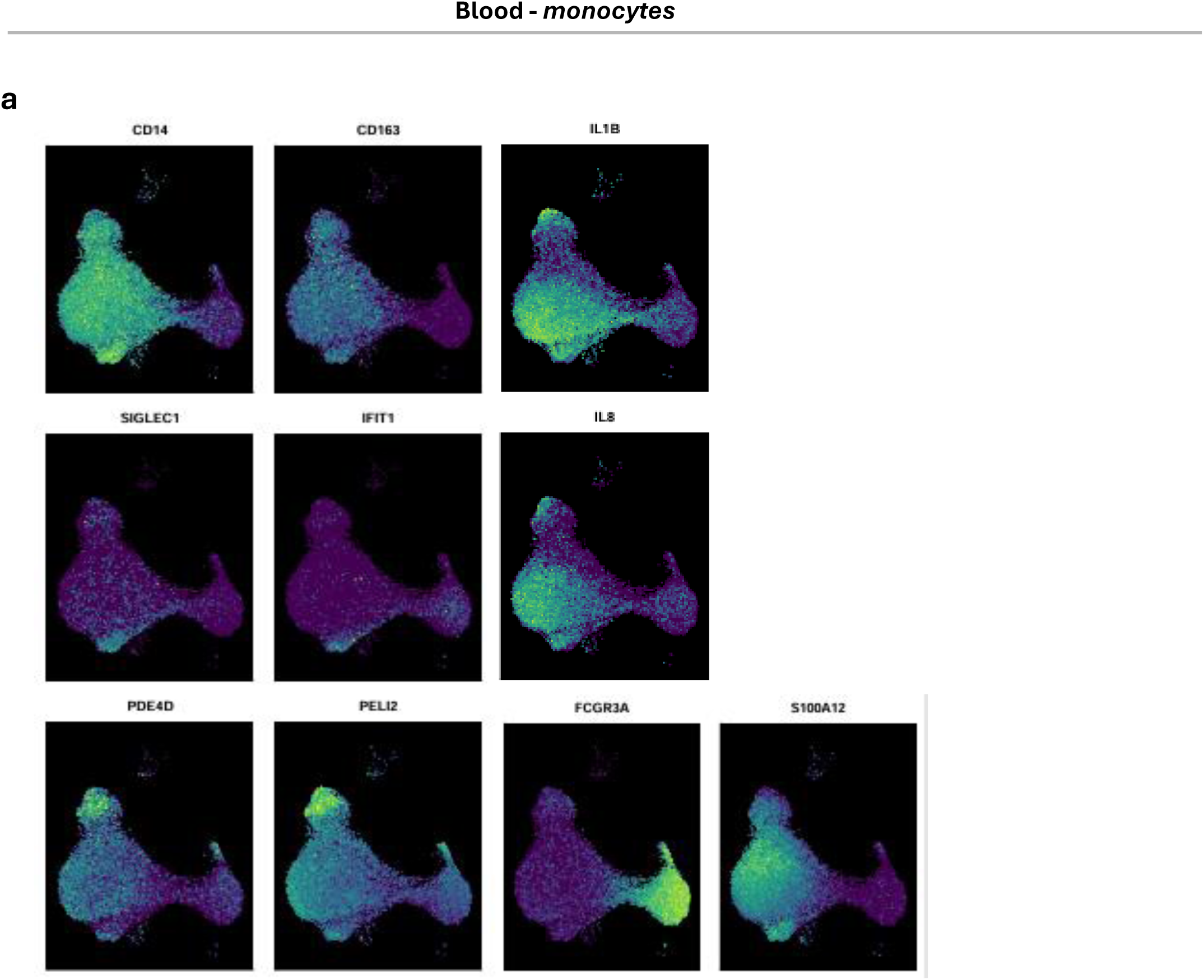
Single-cell analysis ofblood monocytes. **a** Feature plots showing expression of selected marker genes across PBMC-derived monocyte populations. Classical monocyte identity is marked by CD14, while FCGR3A highlights non-classical monocytes. Inflammatory gene expression is reflected by IL1B, IL8, and S100A12, whereas IFIT1 indicates interferon-stimulated responses. CD163 and SIGLEC1 represent immune regulatory/activation-associated programs, and PDE4D and PELI2 are involved in intracellular signaling pathways associated with innate immune activation. **b** Cellular origin of differentially expressed genes (DEGs) in PBMCs from RLA patients. Dot plots show the cell type origin of a 50-gene DEG signature in PBMCs (top) and subclustered monocytes (bottom). Dot size reflects the proportion of cells expressing each gene; colour indicates average expression. Genes associated with shorter transplant-free survival (TFS; dark green) were enriched in monocytes and dendritic cells, while genes decreased in RLA (light green) were primarily derived from T, B, and NK cells.

**Supplementary Figure 6.**
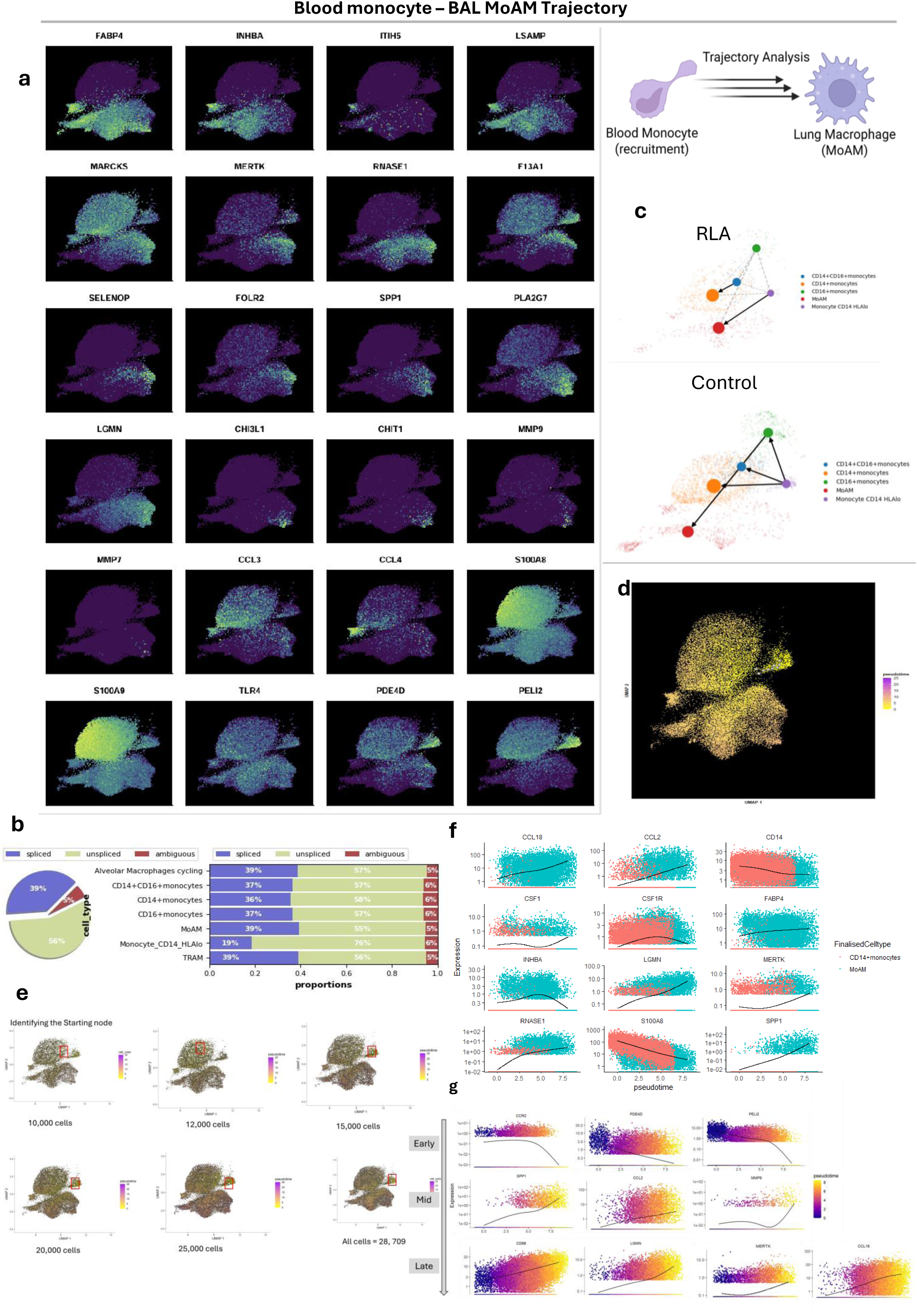
Blood monocyte-BAL macrophage trajectory analysis. **a** Feature plots showing expression of selected genes across the integrated myeloid landscape, including markers of tissue residency (*FABP4, INHBA, LSAMP, RNASE1*), immune regulation/remodelling (*ITIH5, MARCKS, F13A1, SELENOP, FOLR2*), profibrotic mediators (*SPP1, MERTK, LGMN, PLA2G7, CHI3L1, CHIT1, MMP7, MMP9*), inflammatory chemokines (*CCL3, CCL4, S100A8, S100A9*), and signalling/adaptor genes (PDE4D, PELI2). **b** Unspliced and spliced read proportions across blood and BAL myeloid cell types **c** Partition-based graph abstraction (PAGA) velocity graph showing directed connectivity between HLA-DR^lo^PDE4D^hi^ classical monocytes and MoAMs in RLA (top), absent in controls (bottom). **d** Pseudotime ordering of integrated blood CD14⁺ monocytes and BAL MoAMs coloured by pseudotime **e** Identification of root cells for pseudotime analysis using iterative bootstrapping of subsampled cells (10,000–28,709 cells). The most frequently selected high-connectivity node (red box) consistently localized to the *CD14⁺* monocyte compartment. **f.** Gene expression dynamics along the CD14⁺ monocyte (red) to monocyte-derived alveolar macrophage (MoAM; blue) pseudotime trajectory. Smoothed expression curves inferred using Monocle 3 illustrate dynamic regulation of genes associated with chemotaxis (*CCL2, CCL18*), macrophage differentiation (*CD14, CSF1, CSF1R, MERTK*), fibrotic remodeling (*SPP1, LGMN*), and immune regulation (*INHBA, FABP4, RNASE1, S100A8*) across pseudotime. **g.** Gene expression dynamics across integrated blood CD14⁺ monocytes and BAL monocyte-derived alveolar macrophages (MoAMs) colored by pseudotime, highlighting early, intermediate, and late transcriptional programs during monocyte-to-macrophage differentiation.

**Supplementary Figure 7.**
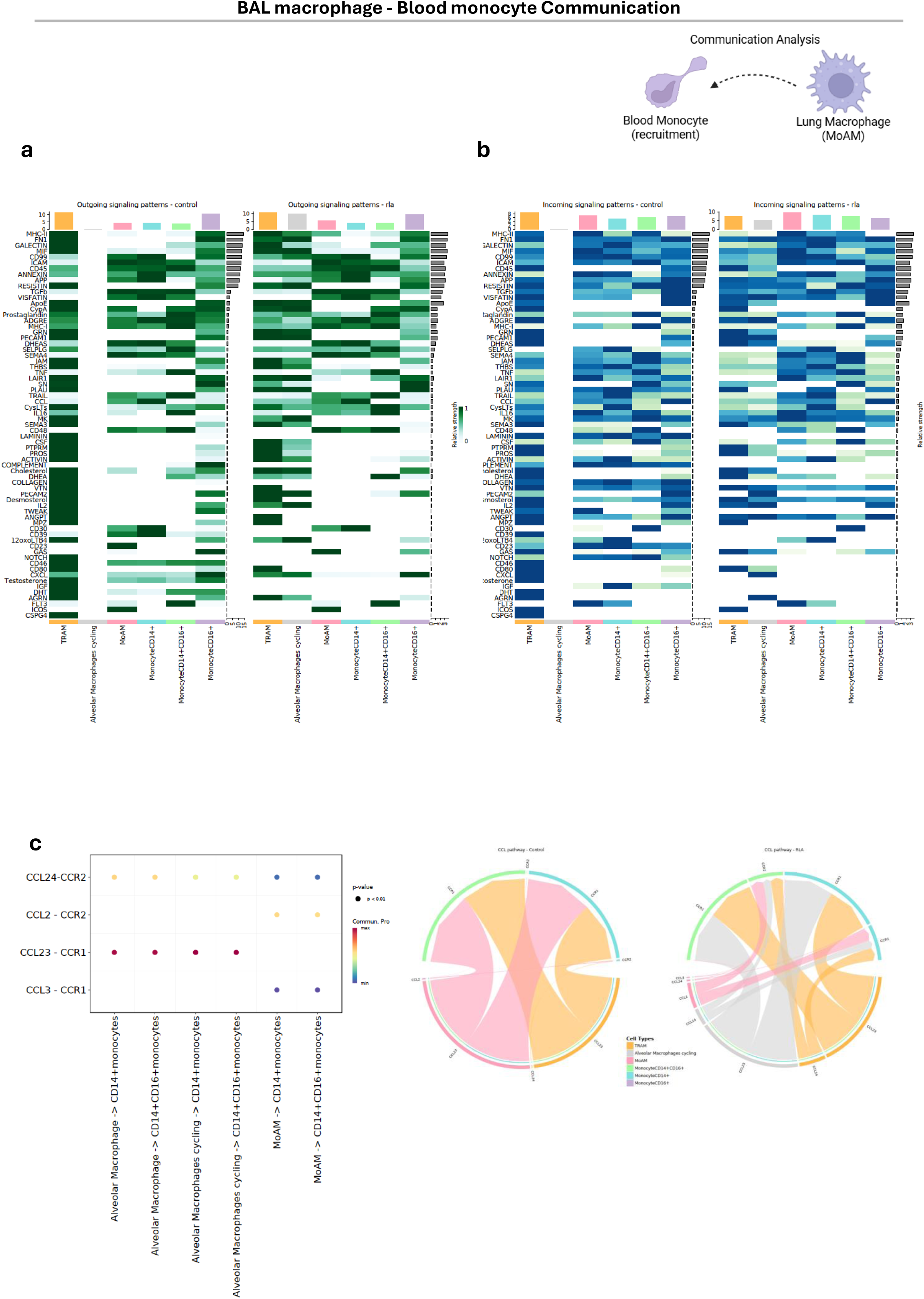
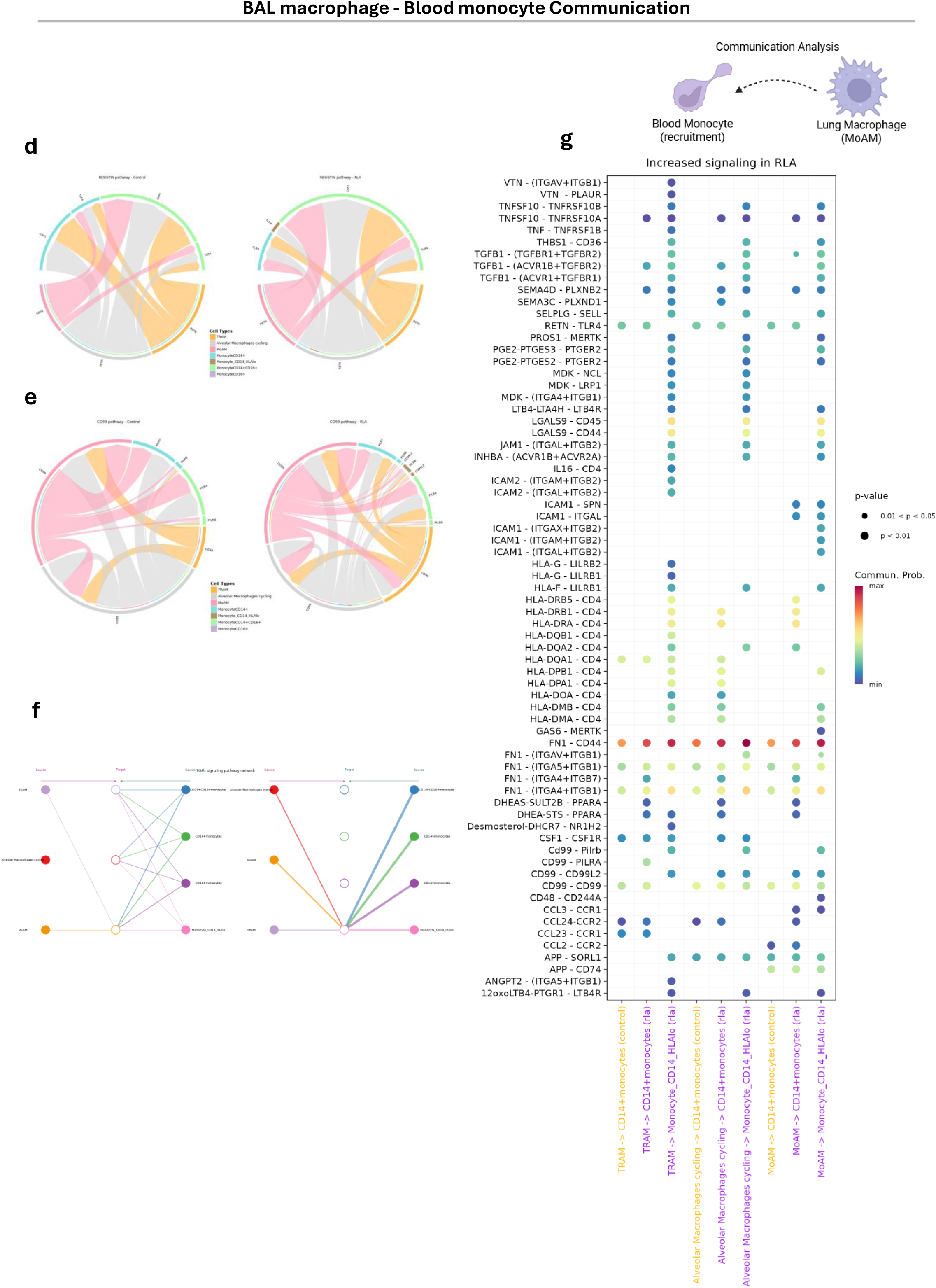
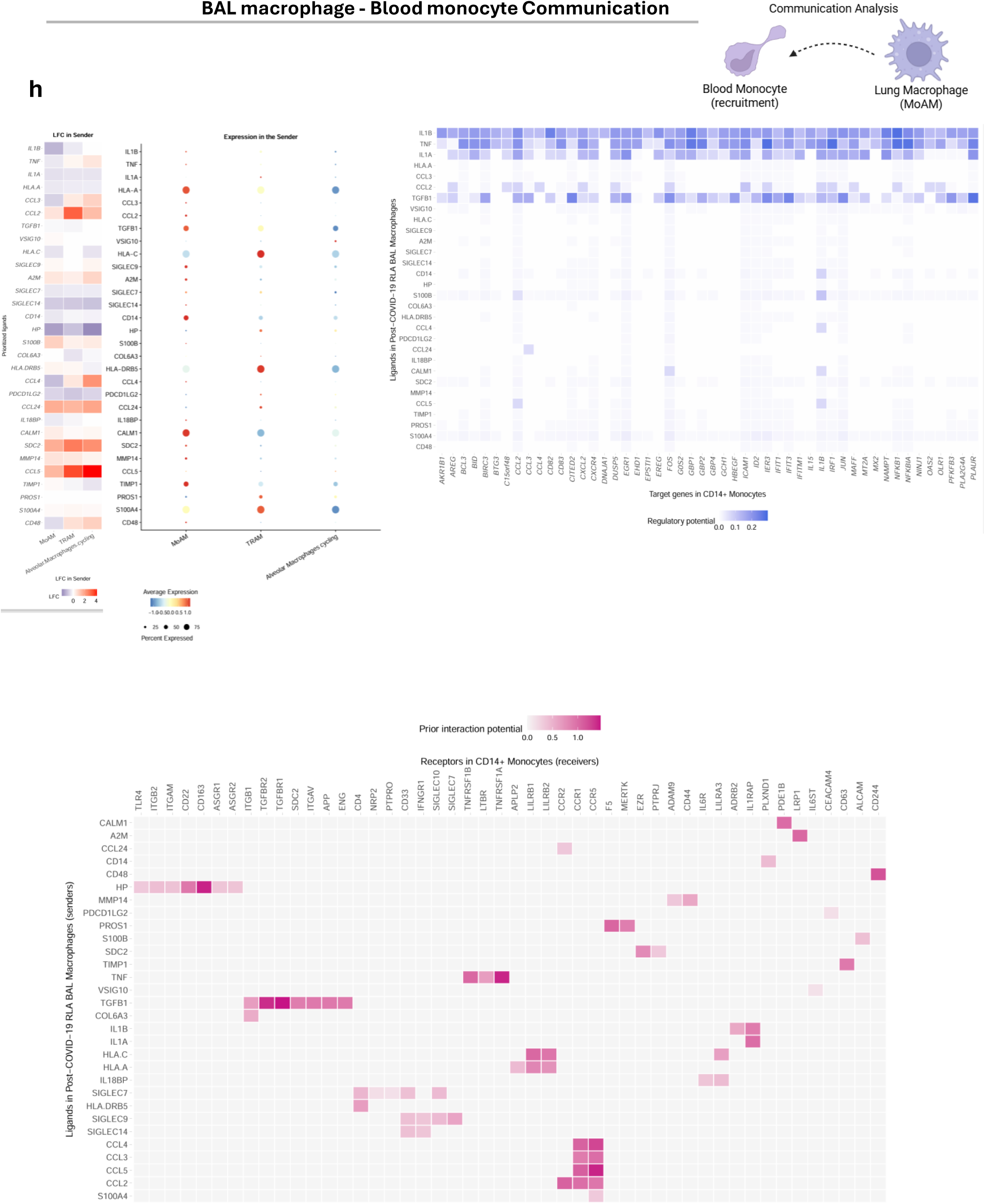
BAL macrophage – Blood monocyte communication analysis. **a–b** Heatmaps showing outgoing (**a**, BAL macrophage ligands) and incoming (**b**, blood monocyte receptors) signalling patterns in myeloid cells from healthy controls (left) and post-COVID-19 RLA patients (right), inferred using CellChat. Each row represents a signalling pathway; columns represent myeloid cell subsets. Colour intensity reflects communication probability, highlighting altered blood-BAL myeloid interactions in RLA. **c** Dot plot (left) and chord diagram (right) showing significant CCL-mediated ligand-receptor interactions between BAL macrophages (senders) and blood monocyte subsets (receivers), inferred using CellChat. Dot size represents significance (p < 0.01); colour indicates interaction probability. Interactions highlight key CCL-CCR axes mediating monocyte recruitment into the lung in RLA. TRAM = tissue-resident alveolar macrophage; MoAM = monocyte-derived alveolar macrophage. **d–f** Condition-specific intercellular signalling differences in RLA compared to controls. Chord diagrams show predicted ligand-receptor interactions within the resistin (*RETN)* (**d**) and *CD99* (**e**) signalling pathway for healthy controls (left) and RLA (right). Cell types are colour-coded, with arrows indicating the direction of communication from sender to receiver cells. Line thickness represents interaction strength. **f** Hierarchical network diagrams illustrate predicted autocrine and paracrine signalling mediated by TGFB and its receptors (TGFB1–TGFBR2) between BAL alveolar macrophage subsets (left) and between BAL macrophages and circulating monocytes (right). Nodes represent cell populations, and edges indicate the direction of ligand-receptor interactions from source to target cells. Edge thickness corresponds to interaction strength, with darker lines indicating stronger predicted signalling. **g** Dot plots showing ligand-receptor pairs with increased signalling from BAL macrophages to blood monocytes in RLA compared to controls, inferred using CellChat. Dot size reflects significance; colour represents interaction probability. Key RLA-enriched signalling pathways include TGFB1-TGFBR2 and FN1-ITGA5/ITGB1, associated with monocyte recruitment, differentiation, and fibrotic remodelling. **h** NicheNet Analysis of BAL Macrophage-to-CD14⁺ Monocyte Communication in post-COVID-19 RLA. Left: Ligand expression and log₂ fold change in BAL macrophages. Middle: Predicted ligand-target interactions in *CD14⁺* monocytes; regulatory potential indicates strength of association. Right: Heatmap of regulatory potential scores for each ligand in *CD14⁺* monocytes. Bottom: Heatmap showing prior interaction potential between ligands and receptors, identifying key interactions driving macrophage-monocyte communication in RLA

**Supplementary Figure 8.**
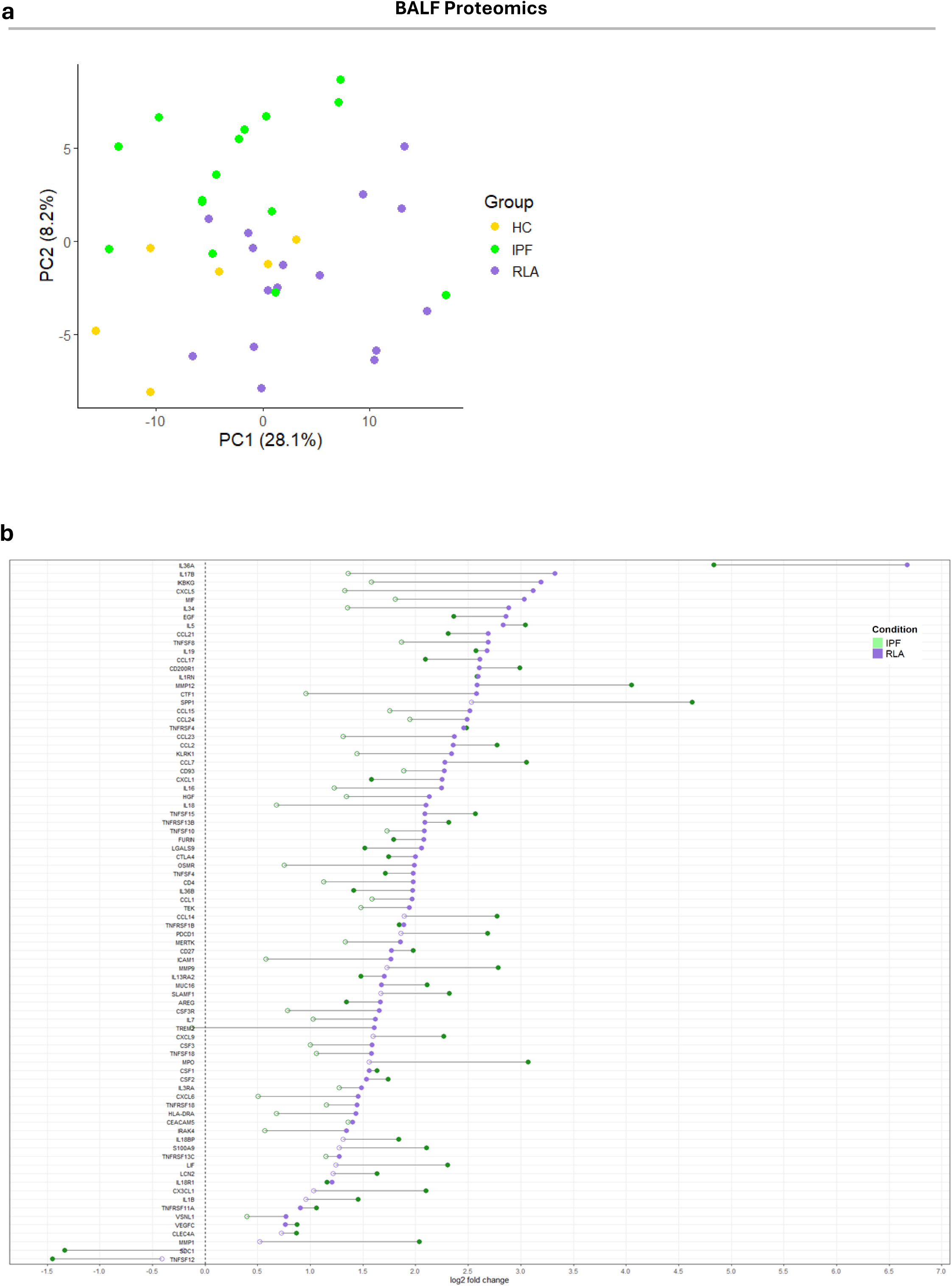
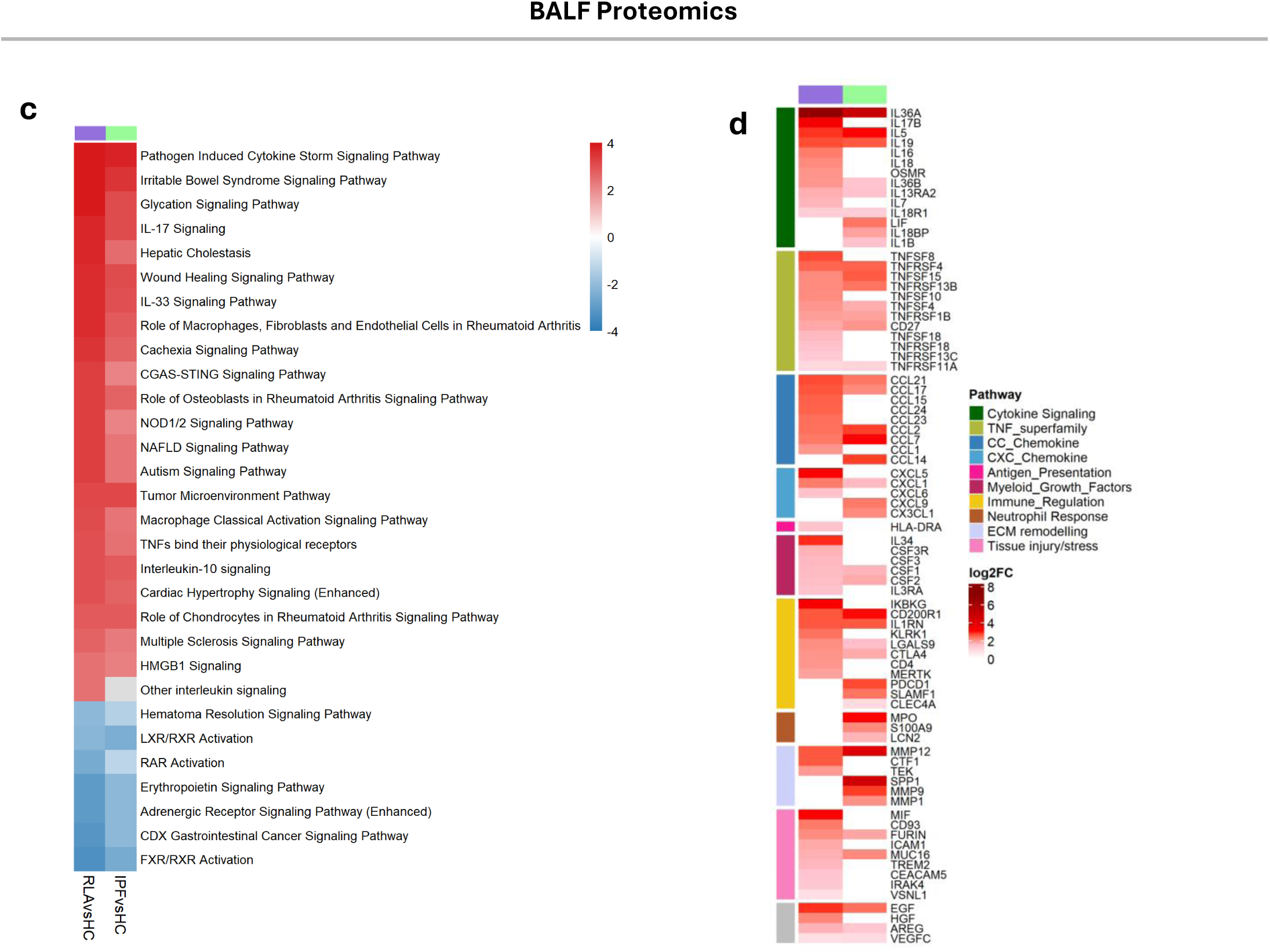
BALF proteomic alterations in post-COVID-19 residual lung abnormalities (RLA) and idiopathic pulmonary fibrosis (IPF). **a** Principal component analysis (PCA) was performed on normalized protein abundance values obtained from bronchoalveolar lavage fluid (BALF) proteomic datasets. Each point represents an individual sample and is coloured by clinical group: healthy controls (HC, yellow), idiopathic pulmonary fibrosis (IPF, green), and post-COVID-19 residual lung abnormalities (RLA, purple). The first two principal components (PC1 and PC2) account for 28.1% and 8.2% of the total variance, respectively. Distribution of samples across the PCA space indicates distinct BALF proteomic patterns among the three groups, with post-COVID-19 RLA samples showing partial overlap with both healthy controls and IPF. **b** Lollipop plot showing differential BALF protein abundance in RLA vs healthy controls (purple) and IPF vs healthy controls (green). Filled circles denote significantly differentially abundant proteins (linear regression, Benjamini–Hochberg FDR < 0.05); open circles denote non-significant proteins. Values are shown as log₂ fold change. **c** IPA canonical pathway analysis performed on significantly differentially abundant proteins from each comparison. Pathways shown met P < 0.05 and | z-score| ≥ 2 in at least one comparison. Heatmap colors indicate predicted pathway activation (red) or inhibition (blue). **d** Heatmap of manually curated functional pathway groups constructed from significantly differentially abundant proteins (FDR < 0.05). Color scale indicates log₂ fold change relative to healthy controls.

**Supplementary Figure 9.**
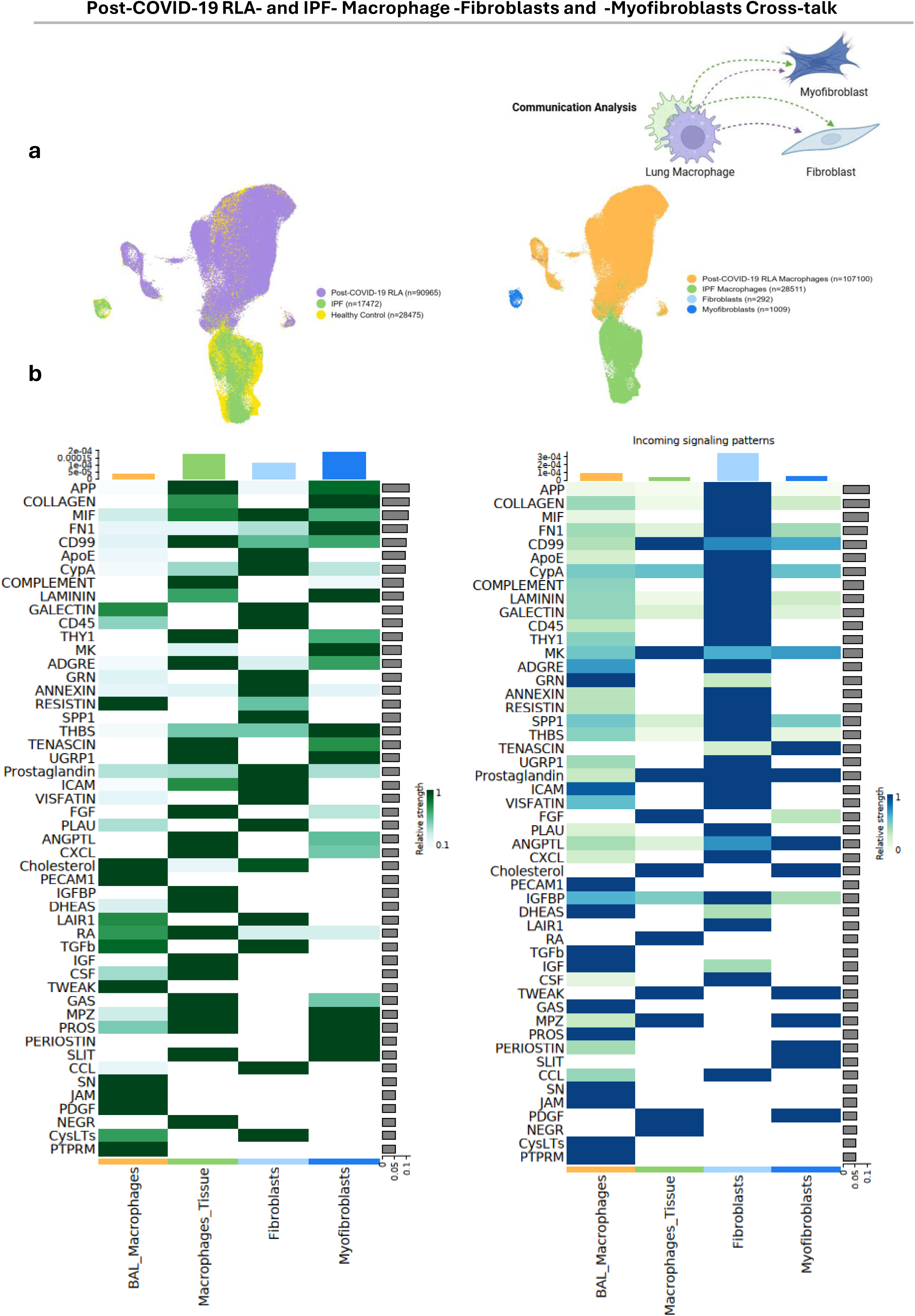
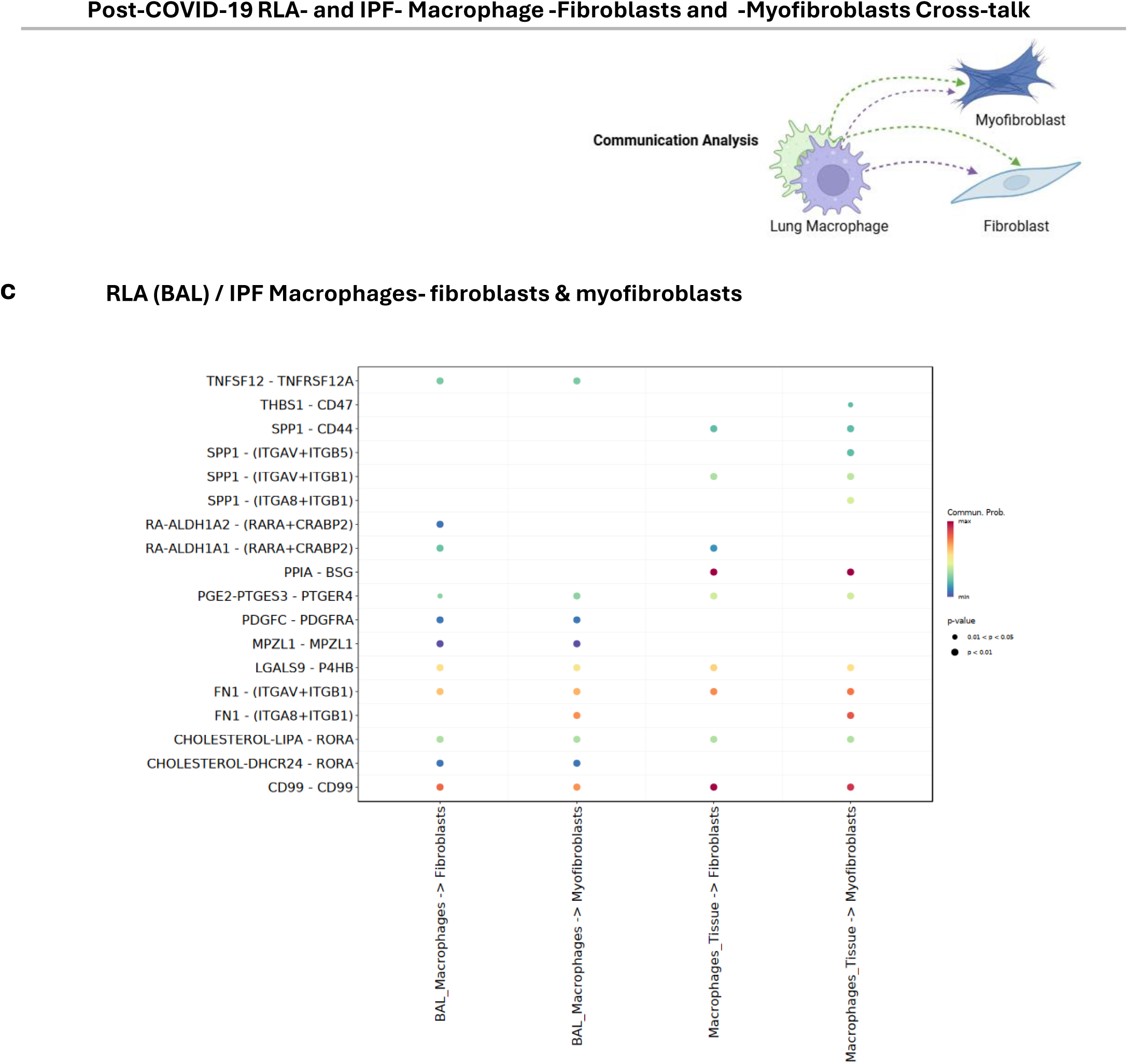
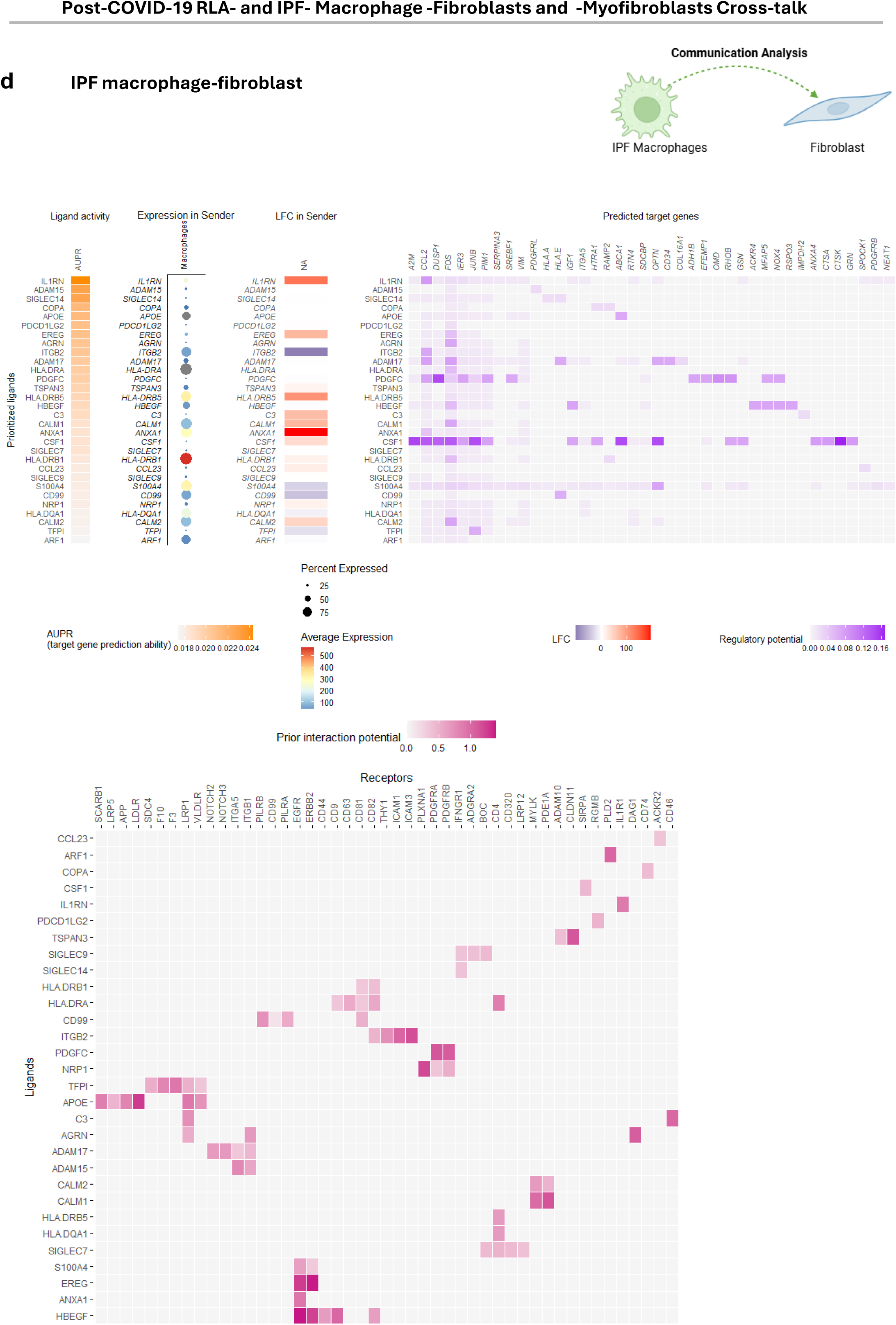
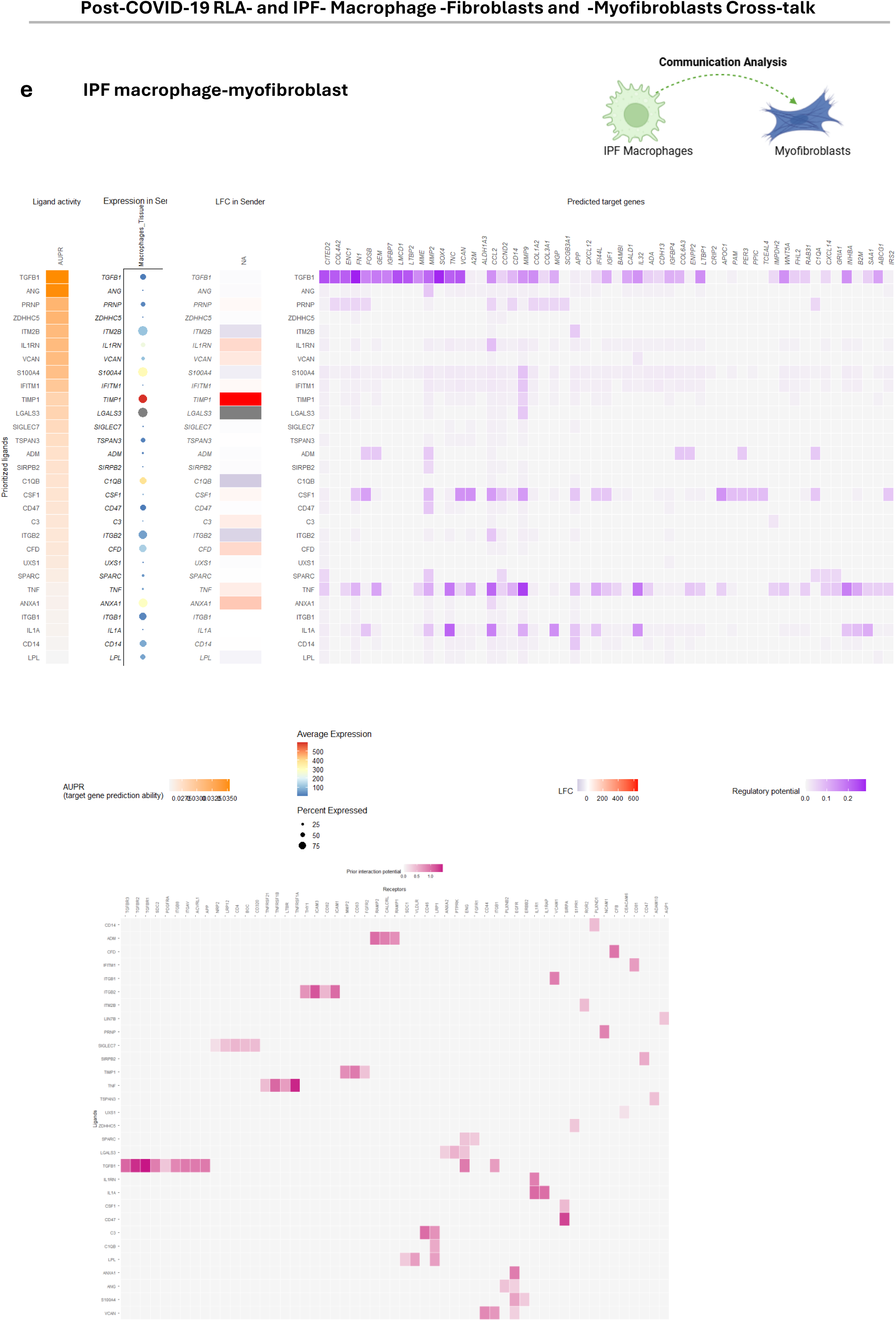
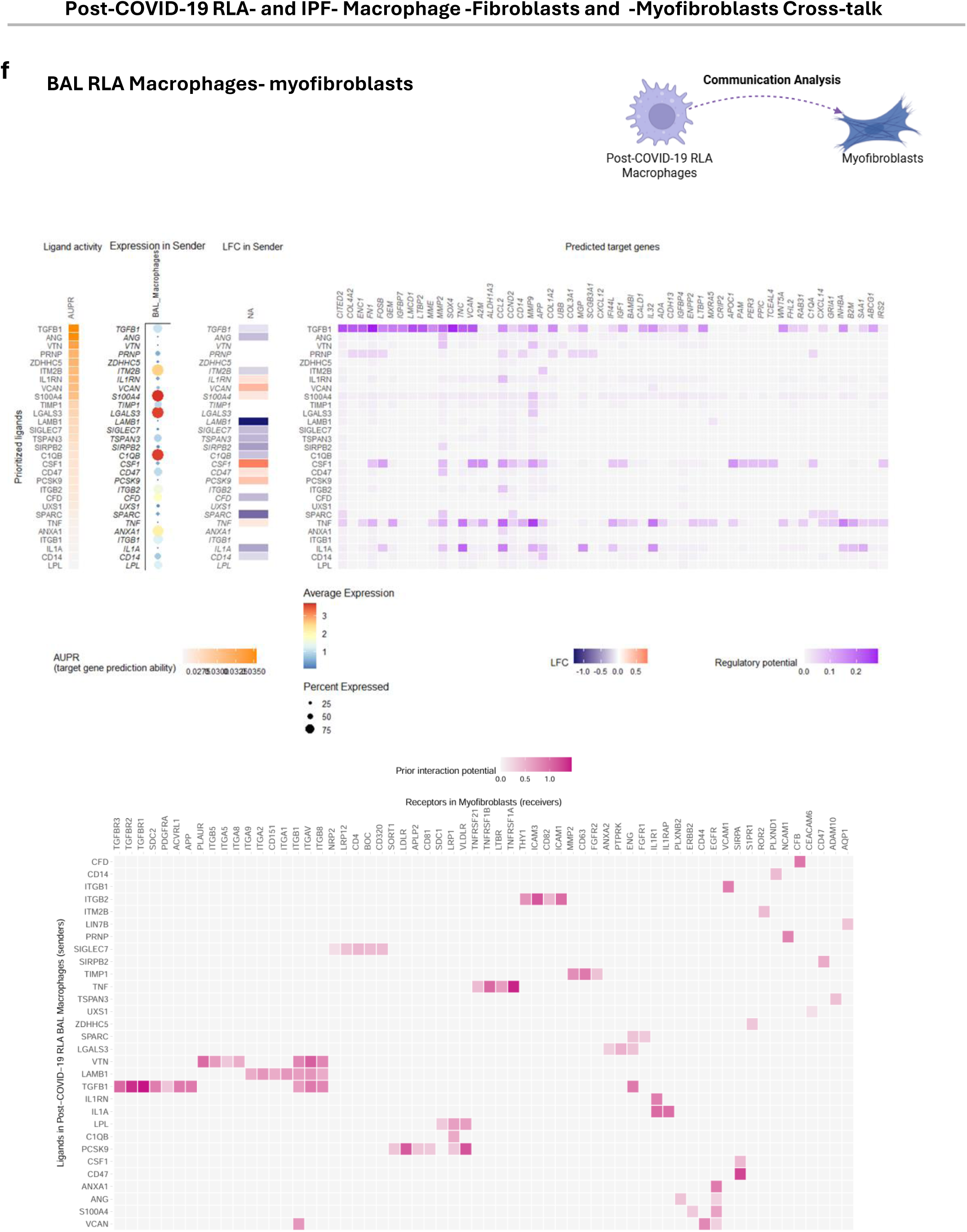
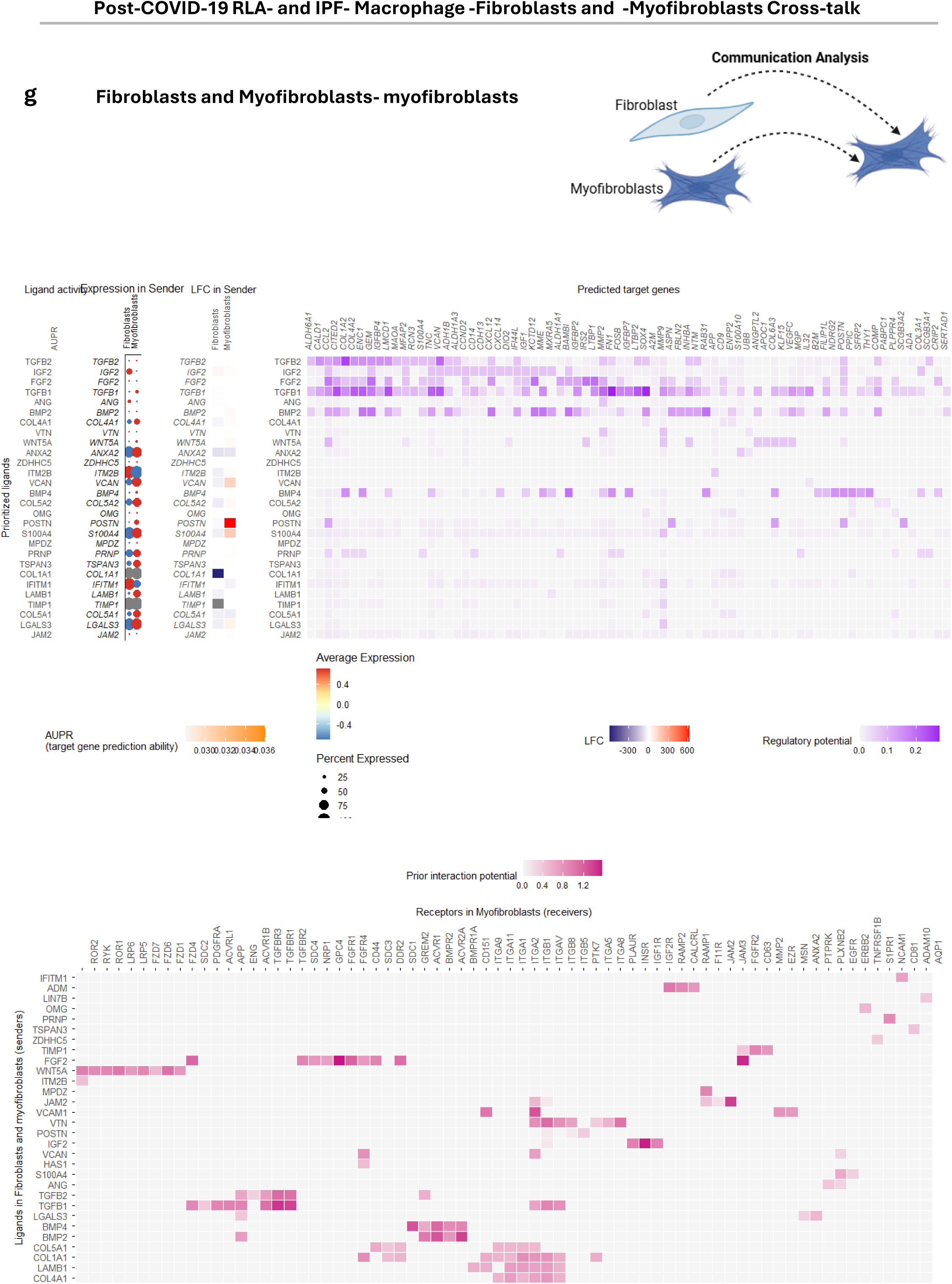
post-COVID-19 RLA- and IPF- Macrophage -Fibroblasts and -Myofibroblasts Cross-talk. post-COVID-19 BAL myeloid data (this study) were integrated with previously published IPF data, after subsetting IPF macrophages, fibroblasts and myofibroblasts. Data integration was performed using Harmony, followed by reclustering and re-annotation. **a** UMAP of integrated cells coloured by condition (RLA = purple, I PF = green, control = yellow; left) and by cell type (right). Cell numbers per group shown in parentheses. **b** Heatmaps showing outgoing (green, ligand-releasing; left) and incoming (blue, receptor-mediated; right) signalling patterns across RLA macrophages, IPF macrophages, fibroblasts, and myofibroblasts, inferred using CellChat. Rows represent signalling pathways; columns represent cell types. **c** Dot plot showing significant ligand-receptor interactions between BAL macrophages (senders) and fibroblasts/myofibroblasts (receivers) and between BAL or IPF tissue macrophages (senders) and fibroblasts/myofibroblasts (receivers). Dot size indicates significance; colour intensity indicates communication probability. Key interactions include FN1-ITGAV/ITGB1, PDGFC-PDGFRA, LGALS9-P4HB, and SPP1-ITGB. **d** NicheNet analysis of ligand-receptor interactions between BAL post-COVID-19 RLA macrophages (senders) and myofibroblasts (receivers). The AUPR score (area under the precision-recall curve) indicates predictive power for target gene regulation. Top heatmap shows predicted ligands and target genes in myofibroblasts ranked by regulatory potential. Dot size reflects ligand expression; colour intensity indicates log₂ fold change. Bottom heatmap shows ligand-receptor pairs; colour intensity reflects predicted interaction strength. **e–f** NicheNet analysis of ligand-receptor interactions between IPF tissue macrophages (senders) and fibroblasts (**f**) or myofibroblasts (**g**) (receivers). Top heatmap shows predicted ligands and target genes in myofibroblasts. Bottom heatmap shows ligand-receptor pairs. **g** NicheNet analysis of ligand-receptor interactions between fibroblasts and myofibroblasts (senders) and myofibroblasts (receivers). Top heatmap shows predicted ligands and target genes in myofibroblasts. Bottom heatmap shows ligand-receptor pairs.

**Supplementary Figure 10.**
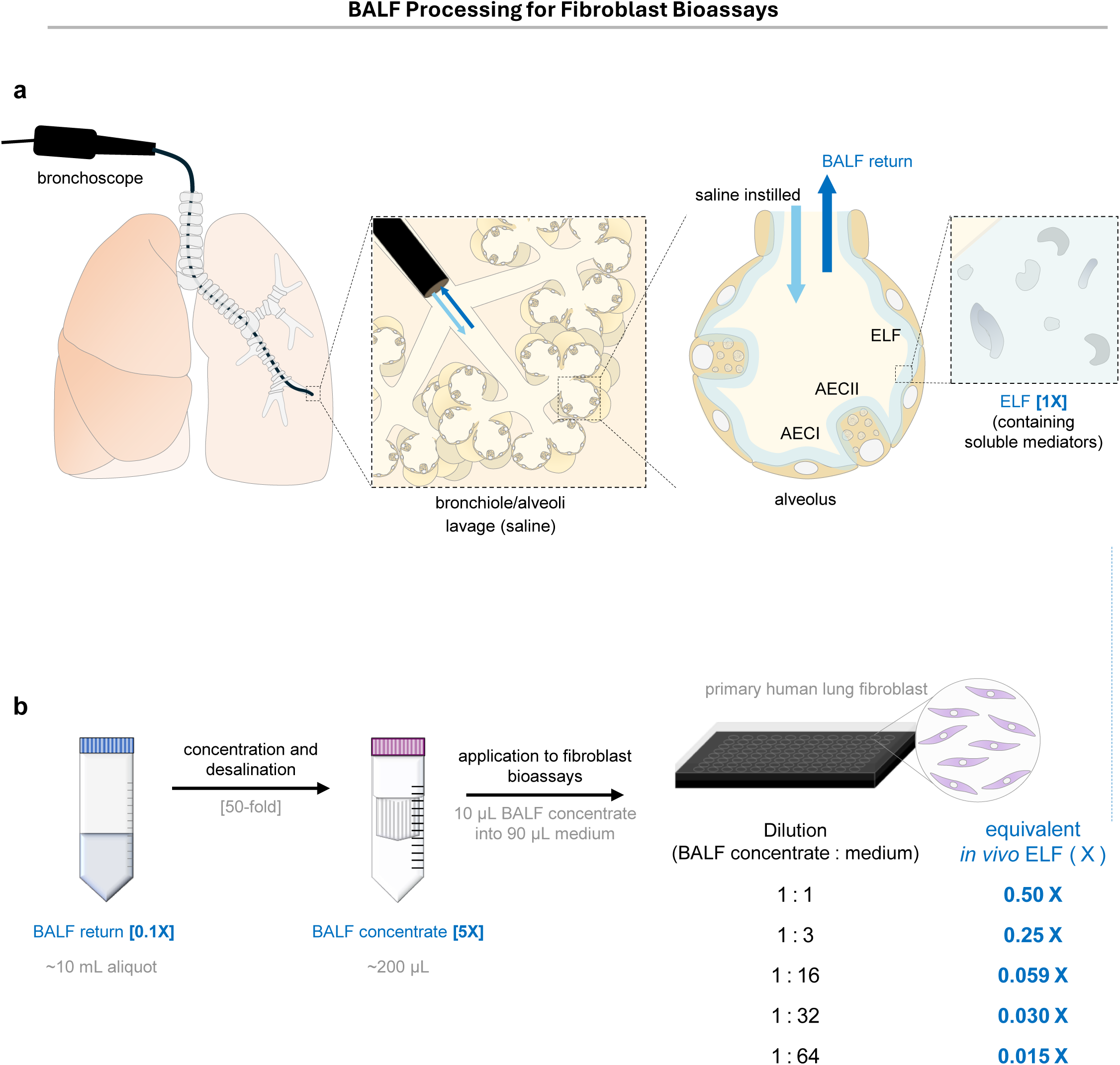
Workflow for BALF collection, processing, and application to fibroblast bioassays. **a** Bronchoalveolar lavage fluid (BALF) was collected by instilling 0.9% saline into the bronchoalveolar spaces via bronchoscopy. The instilled saline washes the bronchioles and alveoli, sampling the epithelial lining fluid (ELF) and recovering both cellular and soluble components. The recovered BALF return contains diluted ELF (∼0.1× the concentration of in vivo ELF). **b** Following collection, BALF samples were maintained on ice, transported to the laboratory, filtered to remove debris, and centrifuged to obtain the acellular supernatant. The BALF return (∼0.1× ELF concentration) was cryopreserved and subsequently concentrated and desalted using Amicon® filtration devices, generating a BALF concentrate corresponding to approximately 5× the in vivo ELF concentration. Aliquots of this concentrate were then diluted into cell culture medium at defined BALF:medium ratios and applied to primary human lung fibroblast bioassays. These dilutions correspond to approximate in vivo ELF equivalents of 0.50× (1:1), 0.25× (1:3), 0.059× (1:16), 0.030× (1:32), and 0.015× (1:64).

**Supplementary Figure 11.**
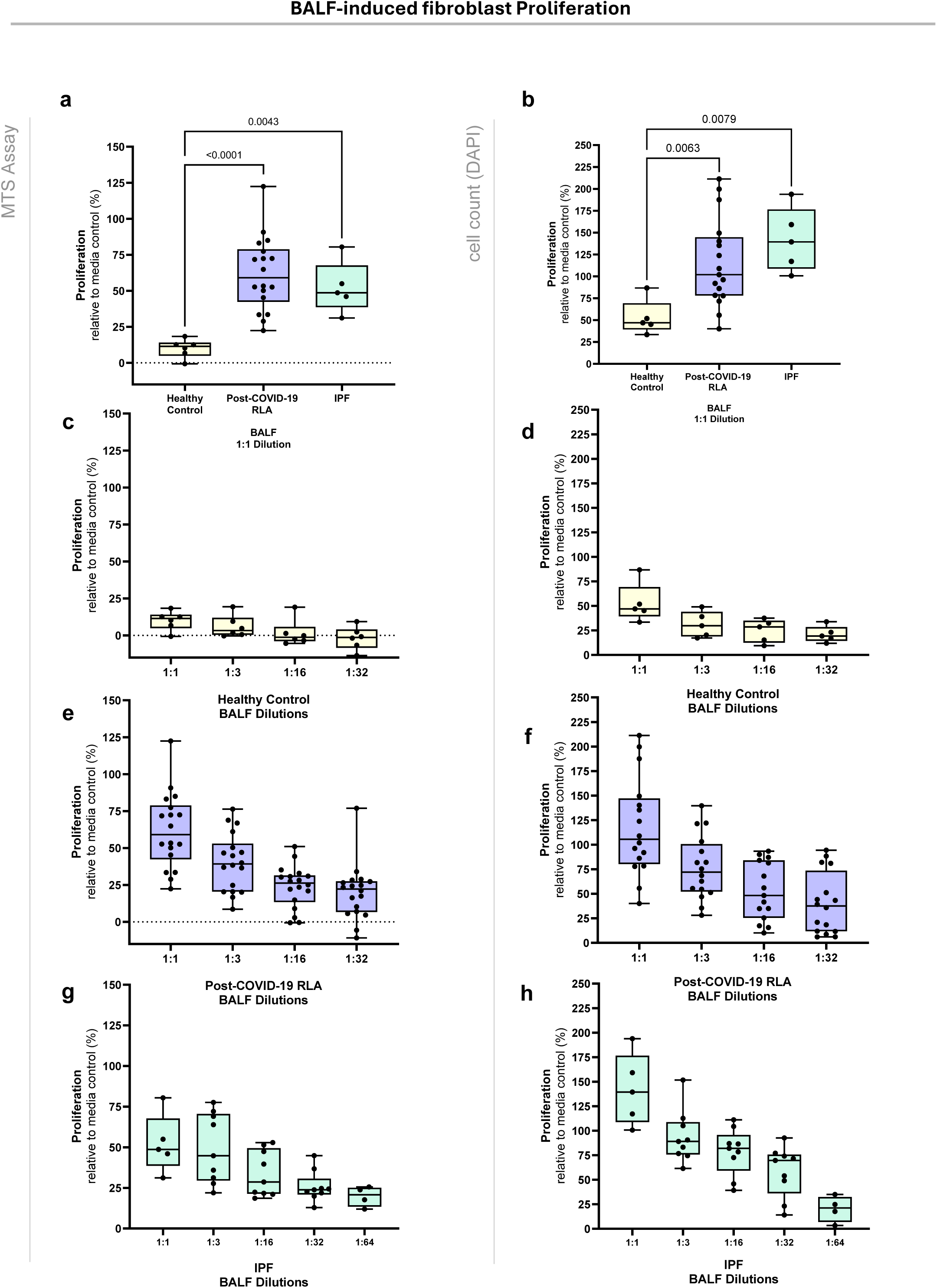
BALF-induced fibroblast proliferation. **a–h** Fibroblast proliferation relative to media control (Plasmax™ + 0.1% FBS; dotted line, y = 0) following exposure to BALF from healthy controls (yellow, n = 6), post-COVID-19 residual lung abnormality (RLA) patients (purple, n = 18), and IPF patients (green, n = 9). Proliferation was quantified by MTS assay (left panels: **a, c, e, g**) and DAPI cell count (right panels: **b, d, f, h**). Data are shown relative to medium control (Plasmax™ + 0.1% FBS; y = 0).Each box represents the interquartile range, the center line indicates the median, and whiskers extend to the minimum and maximum values. Each data point represents the mean of 3–4 technical replicates per biological sample (patient). Biological replicates are displayed with median ± IQR. Statistical comparisons were performed using the Mann–Whitney U test, with p-values indicated. **a–b** Fibroblast proliferation following exposure to BALF from all conditions at 1:1 dilution. **c–d** Fibroblast proliferation following exposure to BALF from healthy controls, at serial dilution **e–f** Fibroblast proliferation following exposure to BALF from post-COVID-19 RLA patients, at serial dilution **g–h** Fibroblast proliferation following exposure to BALF from IPF patients, at seri al dilution

**Supplementary Figure 12.**
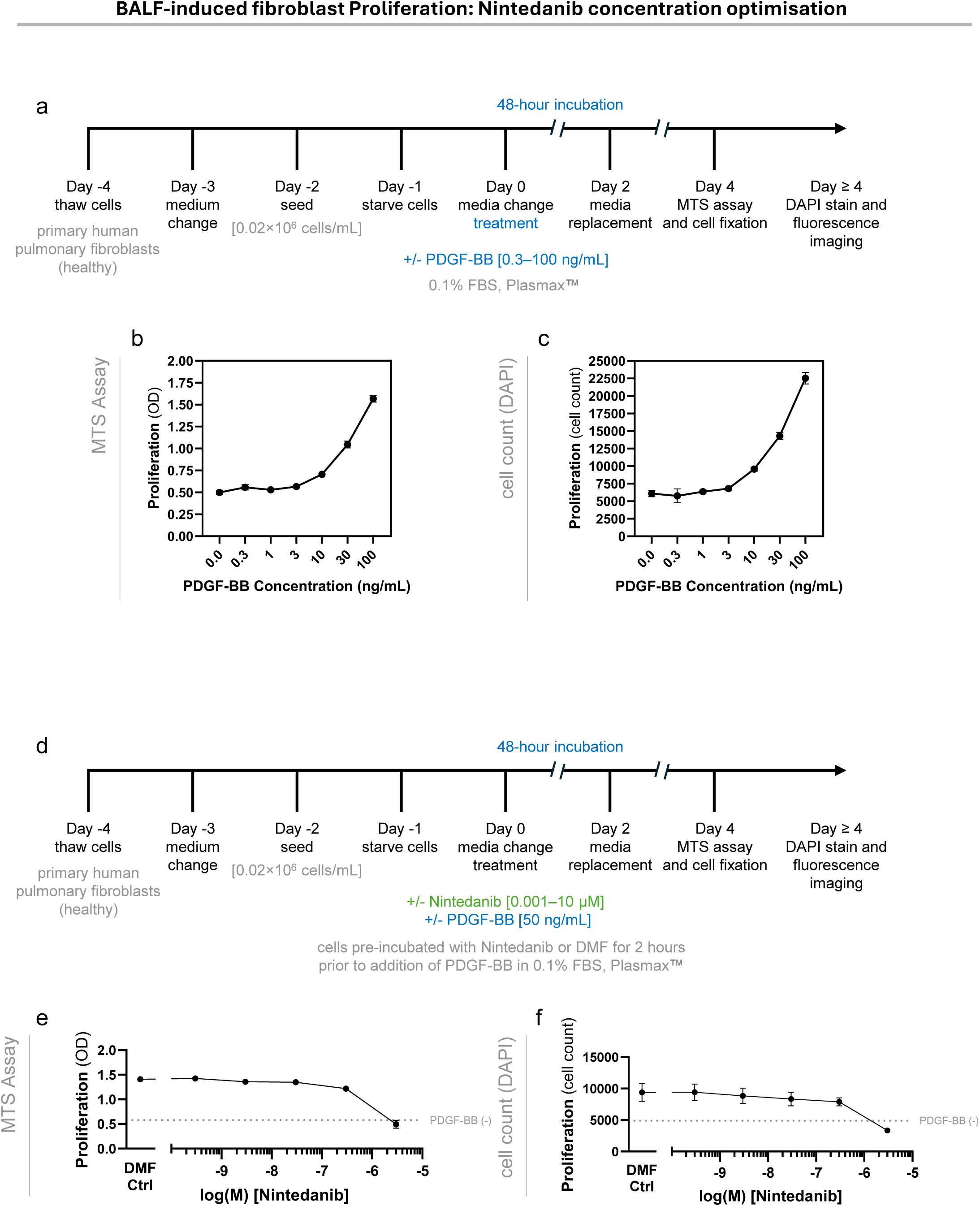
Optimisation of BALF-induced fibroblast proliferation and nintedanib inhibition assays. **a** Schematic illustrating the workflow used to assess fibroblast proliferation in response to recombinant PDGF-BB. Primary human lung fibroblasts were serum-starved and stimulated with PDGF-BB in 0.1% FBS Plasmax™, followed by proliferation measurements after 48 h. **b–c** PDGF-BB–induced fibroblast proliferation across concentrations ranging from 0.3–100 ng/mL, measured by **b** MTS assay (495 nm optical density) and **c** corresponding DAPI-based cell counts. **d** Schematic illustrating the workflow used to determine the nintedanib concentration–response curve in the presence of PDGF-BB. **e–f** Concentration-dependent inhibition of PDGF-BB–induced fibroblast proliferation by nintedanib (0.001–10 µM), measured by **e** MTS assay and f corresponding cell counts. Cells were pre-incubated with nintedanib or vehicle (DMF) for 2 h prior to PDGF-BB stimulation. Wells without PDGF-BB (grey dashed line) served as the negative control. Data represent n = 1 biological experiment with 5–6 technical replicates per condition.

**Supplementary Figure 13.**
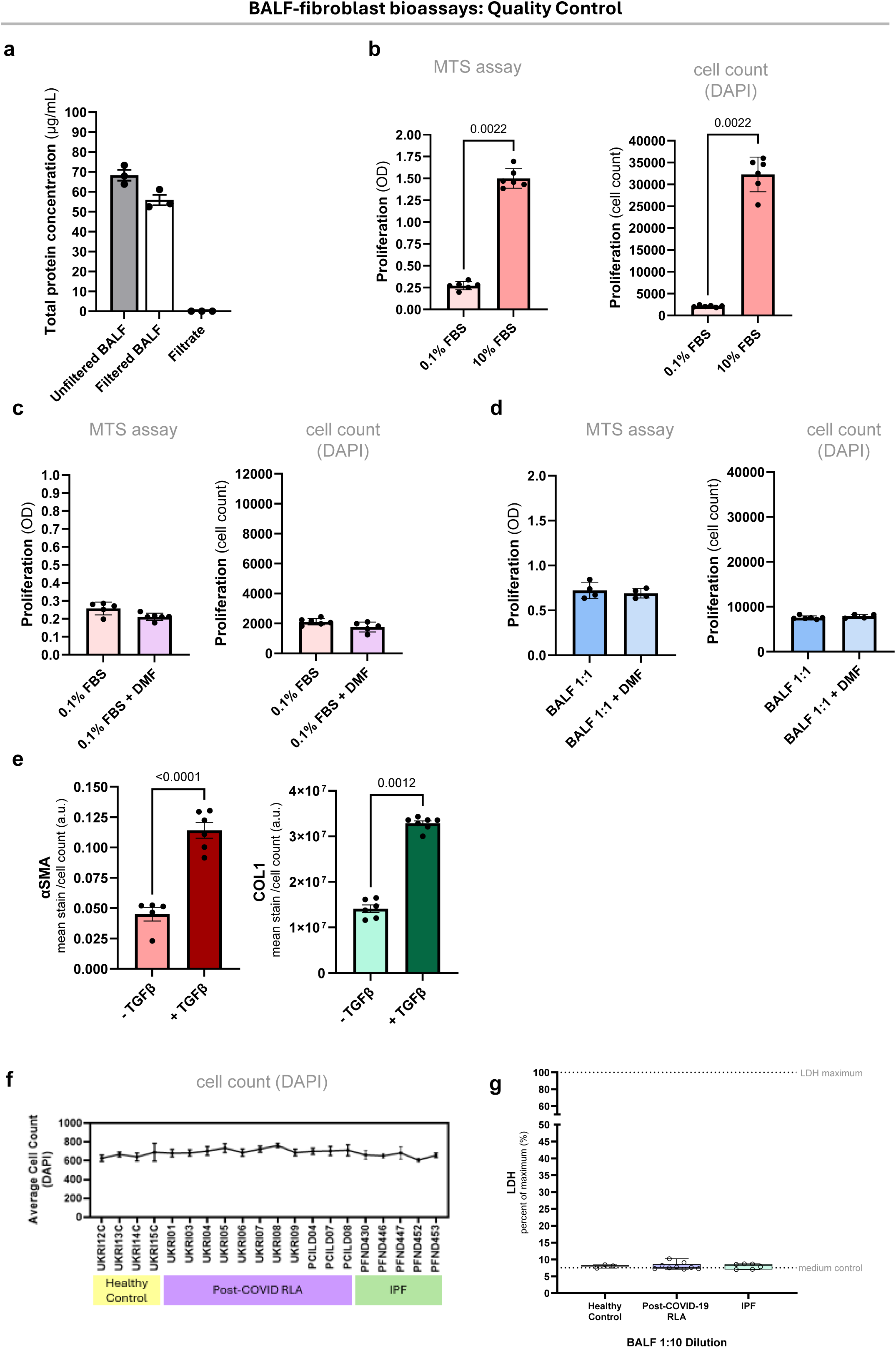
BALF-fibroblast bioassays: Quality Control. **a** Total protein concentration of bronchoalveolar lavage fluid (BALF) samples measured using a BSA protein assay. Protein levels were compared between unfiltered BALF, filtered BALF (retentate after 3 kDa Amicon® filtration), and filtrate. Protein recovery following filtration was ∼82%. **b** Control experiments for fibroblast proliferation assays, assessed by MTS assay (left) and DAPI-based cell count (right). 10% fetal bovine serum (FBS) as a positive control induced robust fibroblast proliferation compared to 0.1% FBS (p=0.0022). Each data point represents the mean of 3-4 technical replicates. Mann-Whitney U test; p-values indicated. **c** Assessment of the nintedanib vehicle control (DMF). Fibroblast proliferation in 0.1% FBS + DMF was compared with 0.1% FBS alone, measured by MTS assay (left) and DAPI-based cell count (right) (n = 1 experiment with 5–6 technical replicates). **d** Pilot experiment assessing the effect of DMF on BALF-induced proliferation. A representative BALF sample at 1:1 dilution was tested with or without DMF, measured by MTS assay (left) and DAPI-based cell count (right) (n = 1 experiment with 4 technical replicates). **e** Validation of fibroblast differentiation and collagen deposition assays performed in Plasmax™ with 0.4% FBS under macromolecular crowding conditions, with or without recombinant transforming growth factor-β (TGF-β, 1 ng/mL). Myofibroblast differentiation was quantified by αSMA expression (left), and collagen I deposition by COL1 staining (right), both normalized to cell count. Statistical comparisons were performed using the Wilcoxon matched-pairs signed-rank test with multiple-testing correcti on using the two-stage linear step-up procedure of Benjamini, Krieger, and Yekutieli. P-values indicate comparisons between conditions with and without TGF-β. **f** DAPI-positive cell counts demonstrating consistent fibroblast numbers across experiments treated with BALF from healthy controls, post-COVID-19 RLA patients, and IPF patients. **g** Cell viability assessment by lactate dehydrogenase (LDH) release in BALF-treated fibroblast cultures (BALF 1:10 dilution). LDH activity was measured using the CytoTox 96® Non-Radioactive Cytotoxicity Assay, with absorbance recorded at 490 nm. Maximum LDH release was determined by lysing cells with lysis solution to establish the assay upper limit.

**Supplementary Figure 14.**
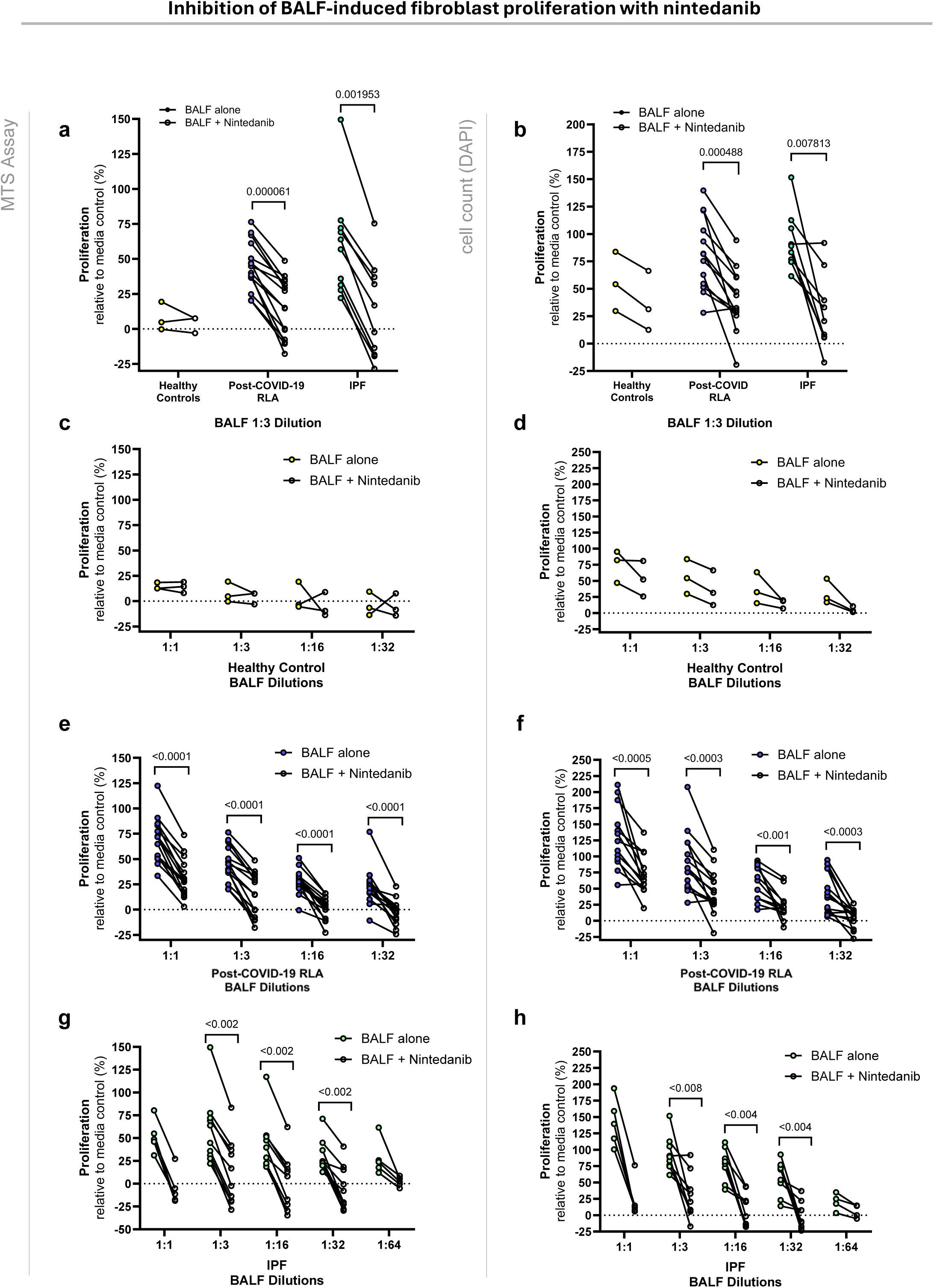
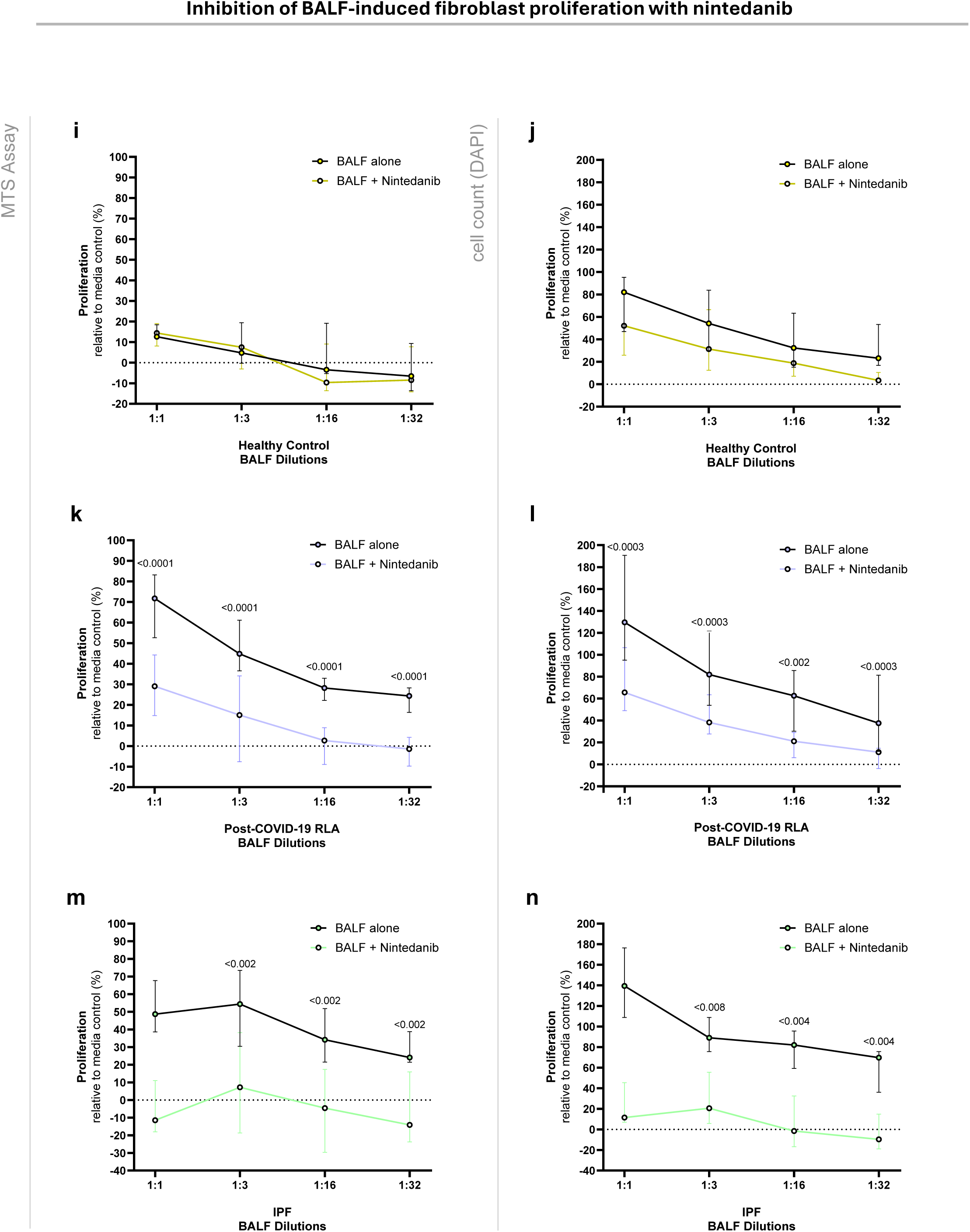
Inhibition of BALF-induced fibroblast proliferation with nintedanib. **a–b** Fibroblast proliferation in response to BALF (1:3 dilution) from healthy controls (n = 3, yellow), post-COVID-19 RLA patients (n = 15, purple), and IPF patients (n = 10, green), with or without nintedanib. Proliferation was assessed by MTS assay (left panels, **a,c,e,g,i,k,m**) and DAPI-based cell count (right panels, **b,d,f,h,j,l,n**). Proliferation was assessed relative to the Plasmax™ 0.1% FBS media control (dotted line, y = 0). **c–h** Serial dilution analysis of BALF from healthy controls (**c–d**), post-COVID-19 RLA patients (**e–f**), and IPF patients (**g–h**) demonstrating concentration-dependent effects on fibroblast proliferation in the presence or absence of nintedanib. Each line represents a single patient sample. Each data point represents the mean of 3-4 technical replicates per biological repeat. Statistical comparisons were performed with a Wilcoxon matched-pairs signed rank test, multiple testing correction was applied using the two-stage linear step-up procedure of Benjamini, Krieger, and Yekutieli to control the false-discovery rate at 5%.. **i–n** Fibroblast proliferation across serial BALF dilutions (1:1–1:32) from healthy controls (**i–j**), post-COVID-19 RLA patients (**k–l**), and IPF patients (**m–n**), measured by MTS assay (left panels) and DAPI-based cell count (right panels). Coloured lines represent BALF + nintedanib, whereas black lines represent BALF alone. Statistical significance was assessed using the Wilcoxon matched-pairs signed-rank test, with multiple testing correction using the Benjamini, Krieger, and Yekutieli procedure to control the false discovery rate at 5%. P-values are indicated.

**Supplementary Figure 15.**
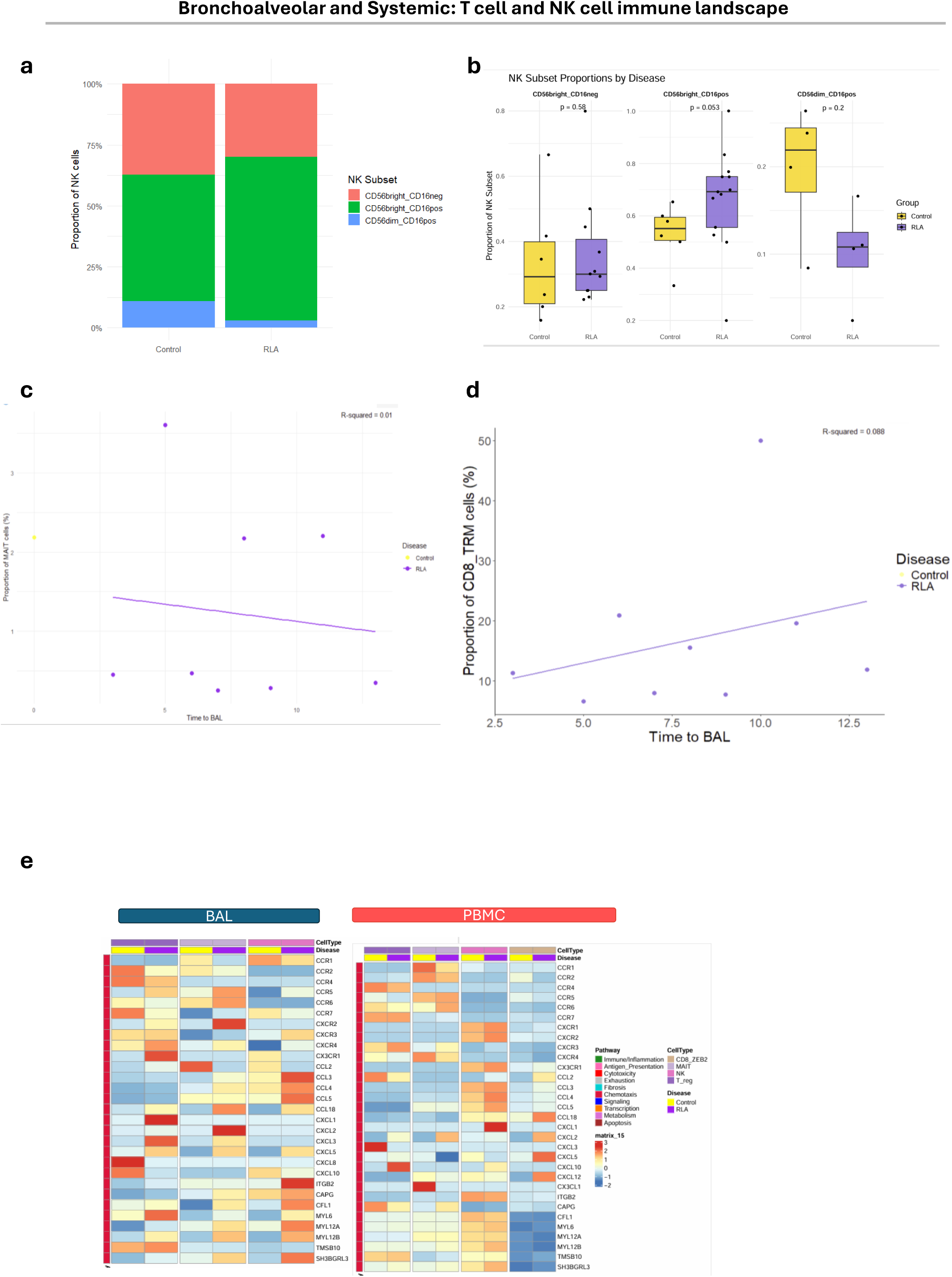
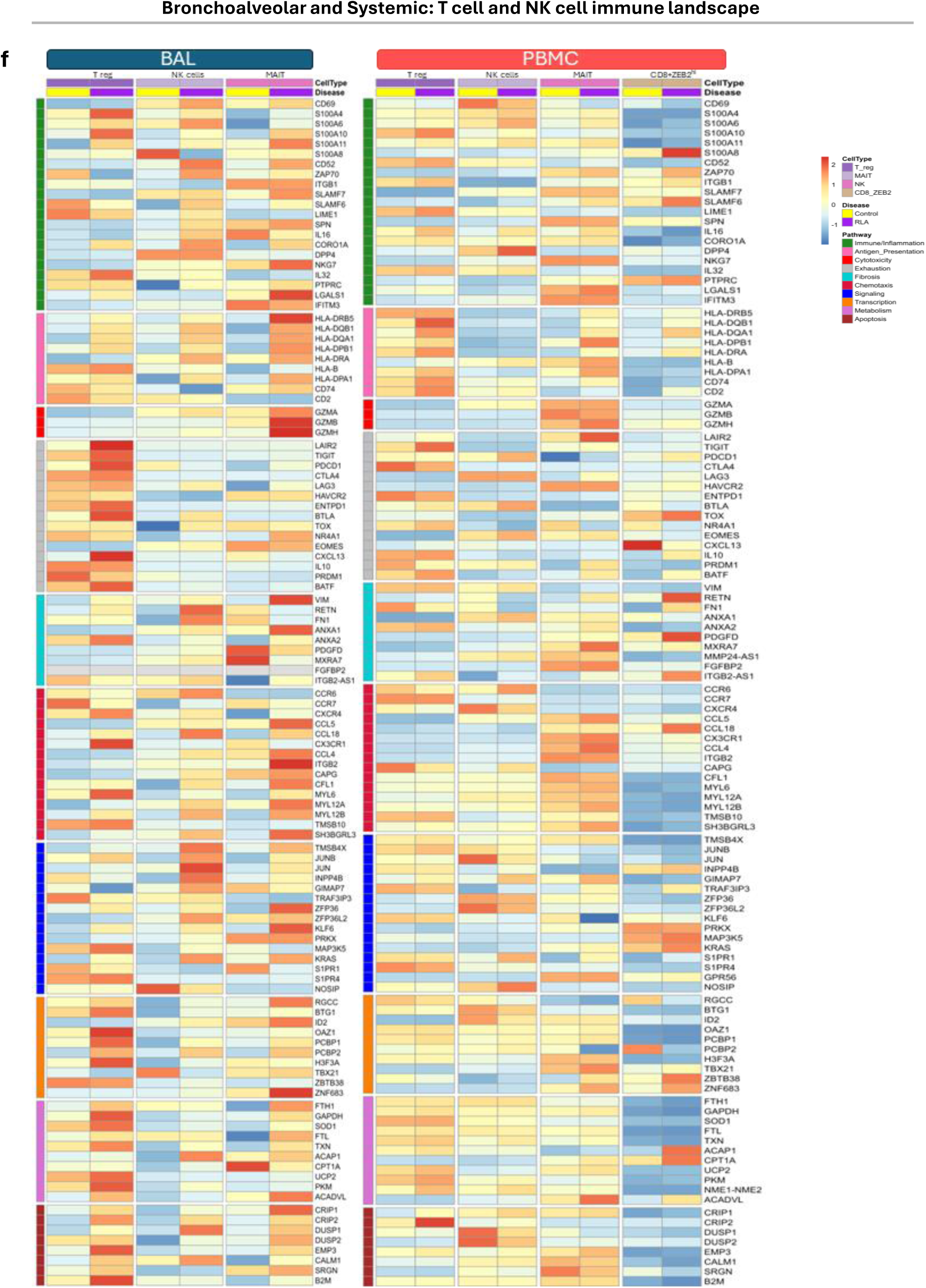
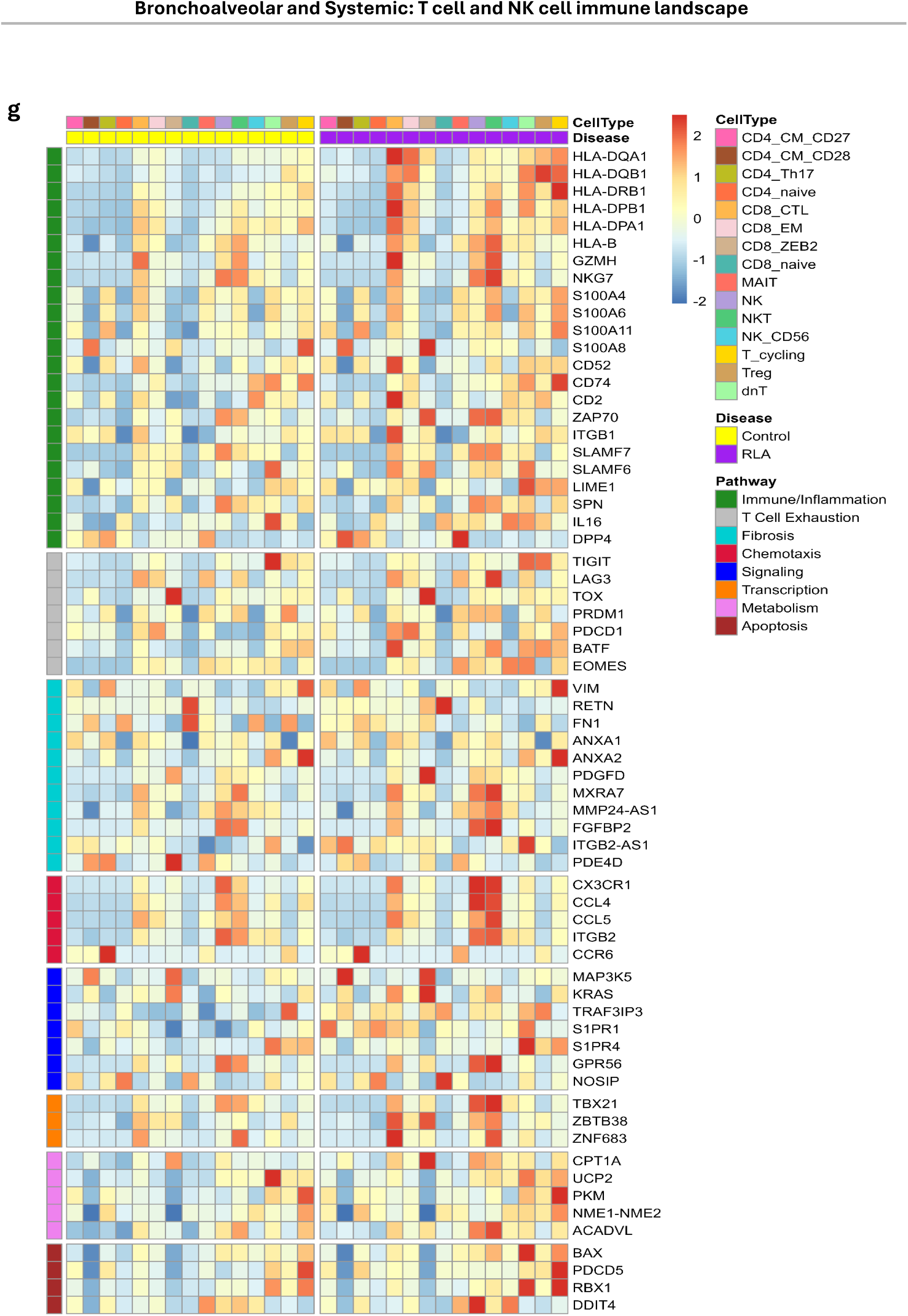
Bronchoalveolar and Systemic - Tcell and NK cell immune landscape. **a,b** Distribution of BAL NK cell subsets in healthy controls and post-COVID-19 RLA patients. a Proportional distribution of NK cell subsets based on pooled cell counts. A significant difference was observed (Chi-squared test, p = 0.0015), with enrichment of CD56^bright^CD16^+^ NK cells in RLA. b Per-patient proportions of each NK subset; p-values from Wilcoxon rank-sum tests were not statistically significant. **c,d** Scatter plots showing the proportion of (c) MAIT cells and (d) CD8⁺ tissue-resident memory T cells (TRM) in BAL plotted against time from acute infection to sampling (Time to BAL) in patients with post-COVID-19 RLA. R² values from linear regression models are shown. **e-g** Heatmaps showing differentially expressed genes in BAL and PBMC T and NK cells from post-COVID-19 RLA patients and healthy controls showing celltypes with compositional changes between conditions (Treg, MAIT, NK and CD8+ZEB2^hi^ cells), focusing on Chemotaxis pathways (e), several pathways of interest (f) and all celltypes in PBMCs (g). Gene expression values represent Z-score normalized log-normalized counts, where each gene’s expression is mean-centered and scaled across all samples. The colour scale represents row-wise Z-scored expression, with red indicating upregulation and blue indicating downregulation in post-COVID-19 RLA patients relative to controls. The top annotation denotes disease groups, with Control in yellow and RLA in purple, alongside cell type classifications. The right annotation categorizes genes by biological pathways. Differential expression analysis was performed using a Wilcoxon rank-sum test via Seurat’s FindMarkers function, with a minimum log₂ fold change of 0.25 and an FDR-adjusted p-value < 0.05 after Bonferroni correction. The heatmap includes the top DEGs per cell type, ranked by log₂ fold change, following the removal of ribosomal genes.

**Supplementary Figure 16.**
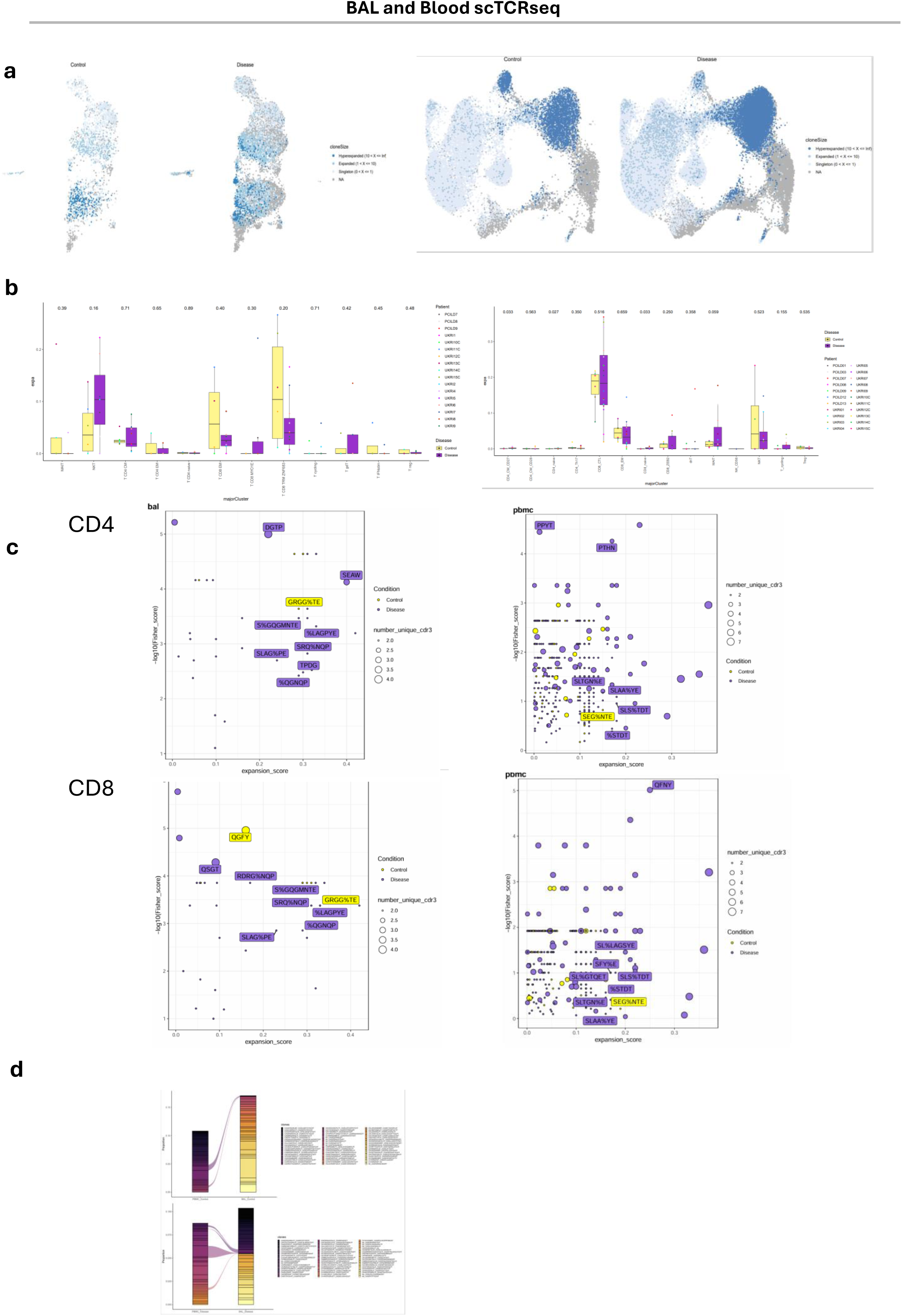
Bronchoalveolar and Systemic scTCRseq. **a** T Cell Clonal Expansion in Control and RLA Conditions. UMAP visualizations display T cell clonal expansion, split by control (left) and RLA (right) conditions and sample type – BAL (left), PBMC (right). Clonal expansion is categorized based on TCR sequence frequency: hyperexpanded clones (≥100 cells, dark blue), expanded clones (10–99 cells, medium blue), singleton clones (2–9 cells, light blue), and non-expanded (NA) clones (grey). **b** Proportion of expanded T cell clones across major T cell subsets in BAL (left) and PBMC (right), stratified by condition (Control, yellow; RLA, purple). Each point represents one patient. Expanded clones are defined by a TCR sequence frequency ≥2 cells. P-values from Wilcoxon rank-sum tests are indicated. **c** GLIPH2 clustering analysis of CD4⁺ (top) and CD8⁺ (bottom) T cell repertoires in BAL (left) and PBMC (right) in post-COVID-19 RLA (purple) and healthy controls (yellow). The x-axis represents the expansion score, indicating the degree of clonal expansion, while the y-axis (−log⁡10(Fisher Score) quantifies the statistical significance of each cluster’s enrichment, determined using Fisher’s exact test. Each point represents a TCR cluster, with the size corresponding to the number of unique CDR3 sequences within that cluster. Clusters exclusive to either disease or control were selected for visualization. The top-ranked clusters, ranked by GLIPH2’ s final_score metric, are labeled with their defining CDR3 motifs. These findings reveal distinct TCR repertoire dynamics between PBMC and BAL, highlighting potential disease-associated T cell responses in post-COVID-19 RLA. **d** Sankey plots showing clonal overlap between BAL and PBMC in Control (top) and RLA (bottom). Each bar represents a unique TCR clone; colour gradients indicating clone frequency within each compartment. Line width corresponds to the proportion of shared clones between compartments, with broader connections indicating higher clonal persistence across compartments. Annotations on the right list the most abundant TCR clones detected in each condition. **e** Expanded TCRα (red) and TCRβ (blue) sequences from BAL (top panels) and PBMC (bottom panels) samples were annotated based on overlap with known virus-specific TCR repertoires, including SARS-CoV-2, CMV, EBV, and HIV. The proportion of expanded clones matching each viral antigen is shown for each compartment. Expanded clones in the BAL exhibited a higher proportion of SARS-CoV-2-specific annotations compared to CMV, EBV, and HIV, suggesting persistent or cross-reactive antigenic stimulation within the lung microenvironment following COVID-19. Annotation was performed by matching CDR3 sequences against curated public databases (VDJ database^56^, Tao et al^57^., and Francis et al^58^.).

## Extended Data Tables

**Extended Data Table 1.**
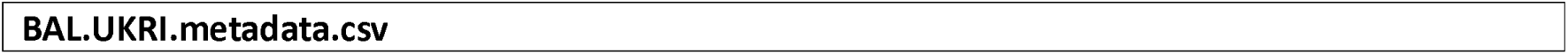
Patient Metadata - BAL.

**Extended Data Table 2.**
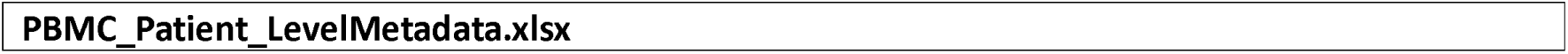
Patient Metadata - PBMC.

**Extended Data Table 3.**
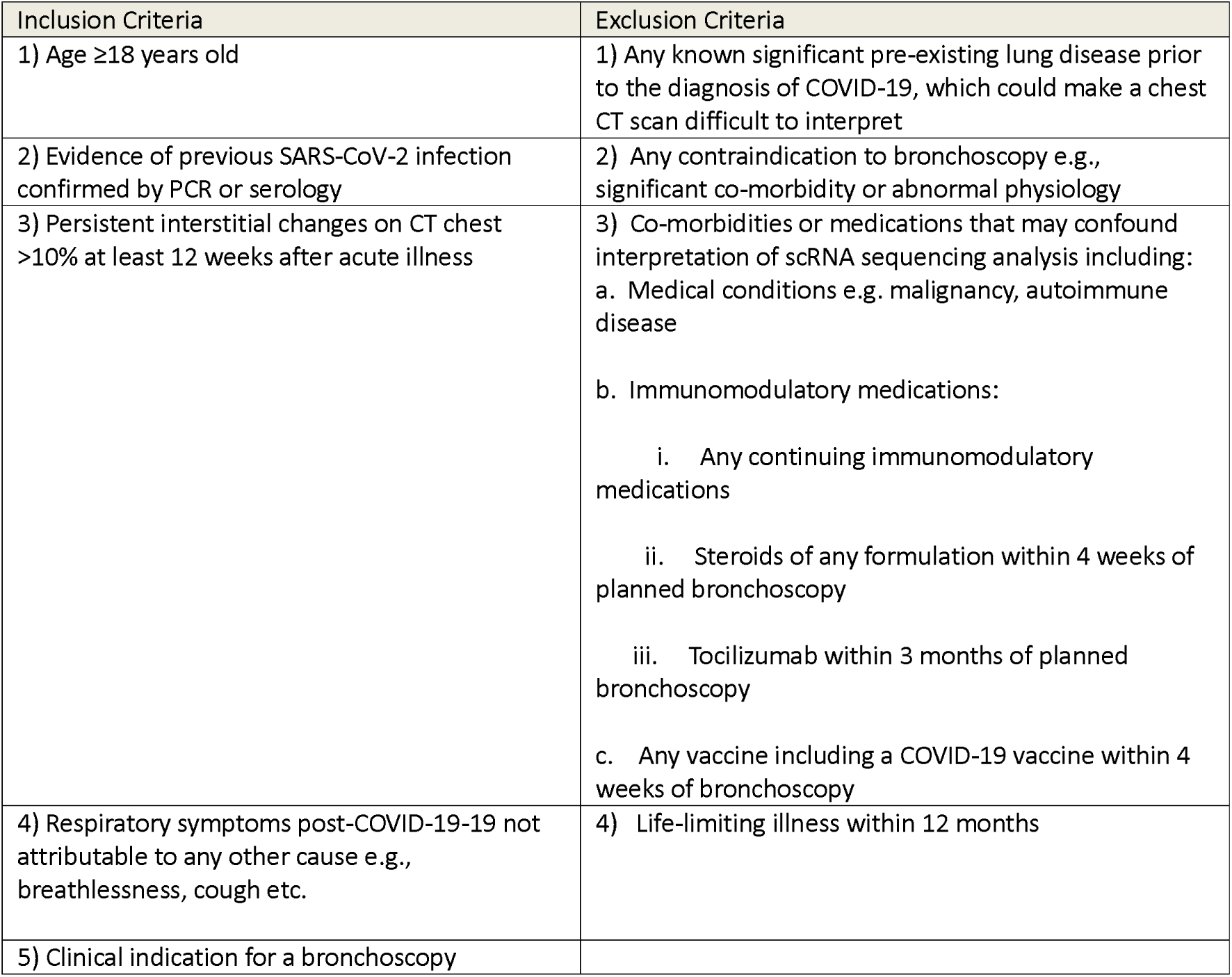
Eligibility Criteria for Patients with post-COVID-19-19 RLA.

**Extended Data Table 4.** Eligibility criteria for Control Cohort (Healthy Controls)

| Inclusion Criteria | Exclusion Criteria |
| --- | --- |
| 1) Age $\geq 18$ years old | 1) Any known significant pre-existing lung disease prior to the diagnosis of COVID-19, which could make a chest CT scan difficult to interpret |
| 2) Normal CT chest scan | 2) Any contraindication to bronchoscopy e.g., significant co-morbidity or abnormal physiology |
|  | 3) Co-morbidities or medications that may confound interpretation of scRNA sequencing analysis including:<br>a. Medical conditions e.g. malignancy, autoimmune disease<br>b. Immunomodulatory medications:<br>i. Any continuing immunomodulatory medications<br>ii. Steroids of any formulation within 4 weeks of planned bronchoscopy<br>iii. Tocilizumab within 3 months of planned bronchoscopy<br>c. Any vaccine including a COVID-19 vaccine within 4 weeks of bronchoscopy |
|  | 4) Life-limiting illness within 12 months |

**Extended Data Table 5.** Characteristics ofpatient groups with bronchoalveolar lavage samples suitable for single cell analysis. Patients PCILD 12 and 13 were excluded from the BAL analysis, with a remaining n=14 samples available for analysis. Data are median (range), n (%), or n/N (%). Percentages might not total 100 where expected owing to rounding. HV = Healthy volunteers; BAL = Broncholveolar lavage; BMI = Body Mass Index; ECMO = Extracorporeal Membrane Oxygenation; I&V = Intubated and ventilated; CPAP = Continuous Positive Airways Pressure; HFNO = High-flow nasal oxygen;

|  | post-COVID-19 RLA All<br>(n=16) | Healthy Controls<br>(n=6) | p |
| --- | --- | --- | --- |
| Age |  |  |  |
| Years | 58.5 (25-79) | 47.5 (28-71) | 0.149 |
| Sex |  |  |  |
| Male | 9 (64%) | 5 (83%) | 0.749 |
| Female | 5 (36%) | 1 (17%) |  |
| Race |  |  |  |
| White | 7 (50%) | 5 (83%) | 0.0471 |
| Asian | 7 (50%) | 0 |  |
| Black | 0 | 1 (17%) |  |
| BMI |  |  |  |
| kg/m<sup>2</sup> | 30 (21-49) | 27 (21-30) | 0.248 |
| Smoking status |  |  |  |
| Current | 1 (6%) | 1 (17%) | 0.442 |
| Former | 2 (13%) | 2 (33%) |  |
| Never | 13 (81%) | 3 (50%) |  |
| Respiratory support in acute COVID-19 |  |  |  |
| ECMO | 3 (21%) | NA |  |
| I&V | 8 (57%) | NA |  |
| CPAP | 1 (7%) | NA |  |
| HFNO | 1 (7%) | NA |  |
| Nil | 1 (7%) | NA |  |
| WHO Severity Score |  |  |  |
|  | 8 (3-9) | NA |  |
| Treatment for acute COVID-19 |  |  |  |
| Steroid | 12 (85%) | NA | 1 * |
| Tocilizumab | 4 (29%) | NA | 0 * |
| Timing of bronchoscopy post-acute COVID-19 |  |  |  |
| Months | 9.5 (3-13) | NA |  |

**Extended Data Table 6.** Metadata for patients with IPF, BAL fluid for bioassays.

**IPF.BALF.metadata.csv**
| Sample_ID | BALF Volume (ml) | Diagnosis | Sex | Date of BAL | Age at time of Bronchoscopy |
| --- | --- | --- | --- | --- | --- |
| L5 | 5 | IPF | Male | 2017-04-24 | 72 |
| L8 | 5 | IPF | Male | 2017-05-15 | 74 |
| L33 | 5 | IPF | Male | 2018-08-14 | 69 |
| L30 | 5 | IPF | Female | 2018-03-06 | 60 |
| L31 | 5 | IPF | Male | 2018-06-05 | 89 |
| 427 | 10 | IPF | Male | 2023-10-25 | 70 |
| 428 | 10 | IPF | Male | 2023-11-01 | 78 |
| 430 | 10 | IPF | Male | 2023-10-30 | 83 |
| 437 | 10 | IPF | Male | 2023-11-20 | 72 |
| 439 | 10 | IPF | Male | 2023-11-27 | 77 |
| 444 | 10 | IPF | Male | 2024-01-04 | 64 |
| 446 | 10 | IPF | Male | 2024-01-22 | 71 |
| 447 | 10 | IPF | Female | 2024-01-22 | 54 |
| 452 | 10 | IPF | Male | 2024-02-05 | 72 |
| 453 | 10 | IPF | Female | 2024-02-19 | 83 |

## Data Availability

The dataset from our study are deposited at the Gene Expression Omnibus (GEO) under accession number (X) and can be explored interactively through a web portal (<u>X</u>). As data are from living patients, these data are available under managed data access.

## Code Availability

The code used to generate the results presented in this manuscript is available as a public repository (TBA on publication)

## Acknowledgements

We thank patients and healthy volunteers and their families for their participation in this study. We acknowledge the members of the UK-ILD Consortium, listed in the Appendix, as collaborators/non-author contributors. We extend thanks to Professor Rick Bucala (Section of Rheumatology, Allergy & Immunology, Yale University) and Professor Erica Herzog (Secti on of Pulmonary, Critical Care & Sleep Medicine, Yale University) for their helpful review of the manuscript.

## Supplementary Data

**Supplementary Data 2**

MetaData for post-COVID-19-19 RLA, Healthy Control and IPF Cohort

**Supplementary Data 3**

QC

## References

1. Thaweethai T, Jolley SE, Karlson EW, et al. Development of a Definition of Postacute Sequelae of SARS-CoV-2 Infection. Jama 2023; 329(22): 1934–46.

2. Mehta P, Rosas IO, Singer M. Understanding post-COVID-19 interstitial lung disease (ILD): a new fibroinflammatory disease entity. Intensive Care Med 2022; 48(12): 1803–6.

3. Bharat A, Querrey M, Markov NS, et al. Lung transplantation for patients with severe COVID-19. Science translational medicine 2020; 12(574).

4. Wendisch D, Dietrich O, Mari T, et al. SARS-CoV-2 infection triggers profibrotic macrophage responses and lung fibrosis. Cell 2021; 184(26): 6243–61.e27.

5. Li C, Qian W, Wei X, et al. Comparative single-cell analysis reveals IFN-γ as a driver of respiratory sequelae after acute COVID-19. Science Translational Medicine; 16(756): eadn0136.

6. Bailey JI, Puritz CH, Senkow KJ, et al. Profibrotic monocyte-derived alveolar macrophages are expanded in patients with persistent respiratory symptoms and radiographic abnormalities after COVID-19. Nature Immunology 2024; 25(11): 2097–109.

7. Karampitsakos T, Jia M, Tourki B, et al. The transcriptome of CD14(+)CD163(-)HLA-DR(low) monocytes predicts mortality in Idiopathic Pulmonary Fibrosis. Eur Respir J 2025.

8. Tourki B, Jia M, Karampitsakos T, et al. Convergent and divergent immune aberrations in COVID-19, post-COVID-19-interstitial lung disease, and idiopathic pulmonary fibrosis. American Journal ofPhysiology-Cell Physiology 2024; 328(1): C199–C211.

9. Schulte-Schrepping J, Reusch N, Paclik D, et al. Severe COVID-19 Is Marked by a Dysregulated Myeloid Cell Compartment. Cell 2020; 182(6): 1419–40.e23.

10. Kumar S, Li C, Zhou L, et al. A distinct monocyte transcriptional state links systemic immune dysregulation to pulmonary impairment in long COVID. Nature Immunology 2026; 27(2): 200–12.

11. Reyfman PA, Walter JM, Joshi N, et al. Single-Cell Transcriptomic Analysis of Human Lung Provides Insights into the Pathobiology of Pulmonary Fibrosis. Am J Respir Crit Care Med 2019; 199(12): 1517–36.

12. Zhao AY, Unterman A, Abu Hussein NS, et al. Single-Cell Analysis Reveals Novel Immune Perturbations in Fibrotic Hypersensitivity Pneumonitis. Am J Respir Crit Care Med 2024; 210(10): 1252–66.

13. Liao M, Liu Y, Yuan J, et al. Single-cell landscape of bronchoalveolar immune cells in patients with COVID-19. Nature Medicine 2020; 26(6): 842–4.

14. Grant RA, Morales-Nebreda L, Markov NS, et al. Circuits between infected macrophages and T cells in SARS-CoV-2 pneumonia. Nature 2021; 590(7847): 635–41.

15. Wauters E, Van Mol P, Garg AD, et al. Discriminating mild from critical COVID-19 by innate and adaptive immune single-cell profiling of bronchoalveolar lavages. Cell Res 2021; 31(3): 272–90.

16. Morse C, Tabib T, Sembrat J, et al. Proliferating SPP1/MERTK-expressing macrophages in idiopathic pulmonary fibrosis. Eur Respir J 2019; 54(2).

17. Ayaub EA, Poli S, Ng J, et al. Single Cell RNA-seq and Mass Cytometry Reveals a Novel and a Targetable Population of Macrophages in Idiopathic Pulmonary Fibrosis. bioRxiv 2021: 2021.01.04.425268.

18. Adams TS, Schupp JC, Poli S, et al. Single-cell RNA-seq reveals ectopic and aberrant lung-resident cell populations in idiopathic pulmonary fibrosis. Sci Adv 2020; 6(28): eaba1983.

19. Habermann AC, Gutierrez AJ, Bui LT, et al. Single-cell RNA sequencing reveals profibrotic roles of distinct epithelial and mesenchymal lineages in pulmonary fibrosis. Science Advances; 6(28): eaba1972.

20. Cambrey AD, Harrison NK, Dawes KE, et al. Increased levels of endothelin-1 in bronchoalveolar lavage fluid from patients with systemic sclerosis contribute to fibroblast mitogenic activity in vitro. Am J Respir Cell Mol Biol 1994; 11(4): 439–45.

21. Foster MW, Thompson JW, Que LG, et al. Proteomic analysis of human bronchoalveolar lavage fluid after subsgemental exposure. J Proteome Res 2013; 12(5): 2194–205.

22. Mehta P, Sanz-Magallón Duque de Estrada B, Denneny EK, et al. Single-cell analysis of bronchoalveolar cells in inflammatory and fibrotic post-COVID lung disease. Front Immunol 2024; 15: 1372658.

23. Xing Y, Ye Y, Zuo H, Li Y. Progress on the Function and Application of Thymosin β4. Frontiers in Endocrinology 2021; Volume 12 - 2021.

24. Ngo LT, Rekowski MJ, Koestler DC, et al. Proteomic profiling of bronchoalveolar lavage fluid uncovers protein clusters linked to survival in idiopathic forms of interstitial lung disease. medRxiv 2024.

25. Vijayakumar B, Boustani K, Ogger PP, et al. Immuno-proteomic profiling reveals aberrant immune cell regulation in the airways of individuals with ongoing post-COVID-19 respiratory disease. Immunity 2022; 55(3): 542–56.e5.

26. Harri son NK, Cambrey AD, Myers AR, et al. Insulin—like Growth Factor-1 is Partially Responsible for Fibroblast Proliferation Induced by Bronchoalveolar Lavage Fluid from Patients with Systemic Sclerosis. Clinical Science 1994; 86(2): 141–8.

27. Roth GJ, Binder R, Colbatzky F, et al. Nintedanib: From Discovery to the Clinic. Journal of medicinal chemistry 2015; 58(3): 1053–63.

28. Flament H, Rouland M, Beaudoin L, et al. Outcome of SARS-CoV-2 infection is linked to MAIT cell activation and cytotoxicity. Nat Immunol 2021; 22(3): 322–35.

29. Jouan Y, Guillon A, Gonzalez L, et al. Phenotypical and functional alteration of unconventional T cells in severe COVID-19 patients. Journal ofExperimental Medicine 2020; 217(12): e20200872.

30. Huang X, Kantonen J, Nowlan K, et al. Mucosal-Associated Invariant T Cells are not susceptible in vitro to SARS-CoV-2 infection but accumulate into the lungs of COVID-19 patients. Virus Research 2024; 341: 199315.

31. Kammann T, Gorin JB, Parrot T, et al. Dynamic MAIT Cell Recovery after Severe COVID-19 Is Transient with Signs of Heterogeneous Functional Anomalies. Journal ofimmunology (Baltimore, Md : 1950) 2024; 212(3): 389–96.

32. Zhang X, Li S, Lason W, Greco M, Klenerman P, Hinks TSC. MAIT cells protect against sterile lung injury. Cell Reports 2025; 44(2): 115275.

33. Odell ID, Steach H, Gauld SB, et al. Epiregulin is a dendritic cell-derived EGFR ligand that maintains skin and lung fibrosis. Sci Immunol 2022; 7(78): eabq6691.

34. Cong J, Wei H. Natural Killer Cells in the Lungs. Front Immunol 2019; 10: 1416.

35. Ran Gh, Lin Yq, Tian L, et al. Natural killer cell homing and trafficking in tissues and tumors: from biology to application. Signal Transduction and Targeted Therapy 2022; 7(1): 205.

36. Maucourant C, Filipovic I, Ponzetta A, et al. Natural killer cell immunotypes related to COVID-19 disease severity. Science Immunology 2020; 5(50): eabd6832.

37. Krämer B, Knoll R, Bonaguro L, et al. Early IFN-α signatures and persistent dysfunction are distinguishing features of NK cells in severe COVID-19. Immunity 2021; 54(11): 2650–69.e14.

38. Cruz T, Agudelo Garcia PA, Chamucero-Millares JA, et al. End-Stage Idiopathic Pulmonary Fibrosis Lung Microenvironment Promotes Impaired NK Activity. The Journal ofImmunology 2023; 211(7): 1073–81.

39. Cruz T, Jia M, Sembrat J, et al. Reduced Proportion and Activity of Natural Killer Cells in the Lung of Patients with Idiopathic Pulmonary Fibrosis. Am J Respir Crit Care Med 2021; 204(5): 608–10.

40. Hostettler KE, Zhong J, Papakonstantinou E, et al. Anti-fibrotic effects of nintedanib in lung fibroblasts derived from patients with idiopathic pulmonary fibrosis. Respiratory Research 2014; 15(1): 157.

41. Wild JM, Porter JC, Molyneaux PL, et al. Understanding the burden of interstitial lung disease post-COVID-19: the UK Interstitial Lung Disease-Long COVID Study (UKILD-Long COVID). BMJ Open Respir Res 2021; 8(1).

42. Yoshida M, Worlock KB, Huang N, et al. Local and systemic responses to SARS-CoV-2 infection in children and adults. Nature 2022; 602(7896): 321–7.

43. Heaton H, Talman AM, Knights A, et al. Souporcell: robust clustering of single-cell RNA-seq data by genotype without reference genotypes. Nature Methods 2020; 17(6): 615–20.

44. Korsunsky I, Millard N, Fan J, et al. Fast, sensitive and accurate integration of single-cell data with Harmony. Nat Methods 2019; 16(12): 1289–96.

45. Michielsen L, Lotfollahi M, Strobl D, et al. Single-cell reference mapping to construct and extend cell-type hi erarchies. NAR Genomics and Bioinformatics 2023; 5(3): lqad070.

46. Bergen V, Lange M, Peidli S, Wolf FA, Theis FJ. Generalizing RNA velocity to transient cell states through dynamical modeling. Nature Biotechnology 2020; 38(12): 1408–14.

47. Lange M, Bergen V, Klein M, et al. CellRank for directed single-cell fate mapping. Nat Methods 2022; 19(2): 159–70.

48. La Manno G, Soldatov R, Zeisel A, et al. RNA velocity of single cells. Nature 2018; 560(7719): 494–8.

49. Trapnell C, Cacchiarelli D, Grimsby J, et al. The dynamics and regulators of cell fate decisions are revealed by pseudotemporal ordering of single cells. Nature Biotechnology 2014; 32(4): 381–6.

50. Wolf FA, Hamey FK, Plass M, et al. PAGA: graph abstraction reconciles clustering with trajectory inference through a topology preserving map of single cells. Genome Biology 2019; 20(1): 59.

51. Jin S, Plikus MV, Nie Q. CellChat for systematic analysis of cell–cell communication from single-cell transcriptomics. Nature Protocols 2025; 20(1): 180–219.

52. Browaeys R, Saelens W, Saeys Y. NicheNet: modeling intercellular communication by linking ligands to target genes. Nat Methods 2020; 17(2): 159–62.

53. Borcherding N, Bormann N, Kraus G. scRepertoire: An R-based toolkit for single-cell immune receptor analysis [version 2; peer review: 2 approved]. F1000Research 2020; 9(47).

54. Zhang L, Yu X, Zheng L, et al. Lineage tracking reveals dynamic relationships of T cells in colorectal cancer. Nature 2018; 564(7735): 268–72.

55. Benjamini Y, Hochberg Y. Controlling the False Discovery Rate: A Practical and Powerful Approach to Multiple Testing. Journal ofthe Royal Statistical Society: Series B (Methodological) 1995; 57(1): 289–300.

56. Shugay M, Bagaev DV, Zvyagin IV, et al. VDJdb: a curated database of T-cell receptor sequences with known antigen specificity. Nucleic Acids Res 2018; 46(D1): D419–d27.

57. Peng Y, Felce SL, Dong D, et al. An immunodominant NP(105-113)-B*07:02 cytotoxic T cell response controls viral replication and is associated with less severe COVID-19 disease. Nat Immunol 2022; 23(1): 50–61.

58. Francis JM, Leistritz-Edwards D, Dunn A, et al. Allelic variation in class I HLA determines CD8+ T cell repertoire shape and cross-reactive memory responses to SARS-CoV-2. Science Immunology; 7(67): eabk3070.

