## Supplementary material for "Multi-omics reveals a monocyte-macrophage-fibroblast axis in post-COVID-19 fibroinflammatory lung remodelling": Figure Legends

**Figure Legends: Main**

**Figure 1. Study design and single-cell clustering and cell compositional results of the bronchoalveolar and systemic cellular landscape in patients with Post-COVID-19 RLA**

**a,** Study overview. **Left panel**, paired bronchoalveolar lavage (BAL) and peripheral blood mononuclear cells (PBMCs) were obtained from patients with post-COVID-19 residual lung abnormalities (post-COVID-19 RLA; n=16) and healthy controls (HC; n=6). Single-cell RNA sequencing (scRNA-seq) with 5′ chemistry and T cell receptor (TCR) sequencing was performed on BAL and PBMC samples, with CITE-seq applied to PBMCs. Trajectory analysis was used to investigate blood monocyte transition to monocyte-derived alveolar macrophages (MoAM). Immune–stromal communication analyses were conducted using macrophage–fibroblast interactions from post-COVID-19 and idiopathic pulmonary fibrosis (IPF) data and differentially expressed BAL fluid (BALF) proteomics-fibroblast. **Right panel**, BALF proteomics was performed using the Alamar Nucleic Acid Linked Immuno-Sandwich Assay (NULISA)-seq platform. BALF was applied to primary human lung fibroblasts to assess proliferation (MTS assay and DAPI cell counts) and the effect of the anti-fibrotic agent, nintedanib, and fibroblast differentiation and collagen deposition using macromolecular crowding assays. **b-i,** BAL (**b-e**) and PBMC (**f-i**) single-cell clustering and compositional analysis. BAL (**b-c**) and PBMC (**f–g)** uniform manifold approximation and projection (UMAP) visualisation of transcriptomes, annotated by major cell types (**b, f**) and coloured by condition (**c, g**). Post-COVID-19 RLA = purple; HC = yellow. Cell plotting order was randomised between conditions to improve visualisation. Dot plots show canonical marker genes used for cell-type annotation for BAL (**d**) and PBMC (**h**). BAL (**e**) and PBMC (**i**) differential cell abundance analysis comparing post-COVID-19 RLA and HC samples Log₂ fold change (LFC) is shown, with red indicating increased and blue decreased abundance in post-COVID-19 RLA. Circle size reflects –log₁₀(FDR) from Wilcoxon rank-sum test with Bonferroni correction (FDR < 0.05, indicated by yellow border). Dashed line indicates LFC = 0.

BAL, bronchoalveolar lavage; BALF, bronchoalveolar lavage fluid; CITE-seq, cellular indexing of transcriptomes and epitopes by sequencing; DC1, dendritic cells (type 1); DC2, dendritic cells (type 2); DC activated, activated dendritic cells; dNT, double negative T cells; FDR, false discovery rate; HC, healthy controls; HSPC, haematopoietic progenitor cells; IPF, idiopathic pulmonary fibrosis; LFC, log₂ fold change; MAIT, mucosal-associated invariant T cells; MoAM, monocyte-derived alveolar macrophages; Mono, monocytes; NK, natural killer cells; pDCs, plasmacytoid dendritic cells; RLA, post-COVID-19 residual lung abnormalities; TCR, T cell receptor; and Treg, regulatory T cells.

**Figure 2. Myeloid single-cell analysis of bronchoalveolar and blood compartments in patients with Post-COVID-19 Residual Lung Abnormalities**

**a–j,** BAL (**a–e**) and PBMC (**f-j**) single-cell clustering and differential abundance analysis of myeloid cells. BAL (**a, b**) and PBMC (**d, e**) uniform manifold approximation and projection (UMAP) visualisation of myeloid labelled by cell type (**a, f**); cell numbers in parentheses) or condition (**b, e)**; HC = yellow, post-COVID-19 RLA = purple. Differential cell abundance analysis comparing post-COVID-19 RLA vs HC in BAL (**c**) and PBMC (**f**). Log₂ fold change (LFC) is shown, with red indicating increased and blue decreased abundance in post-COVID-19 RLA. Circle size reflects –log₁₀(FDR) from Wilcoxon rank-sum test with Bonferroni correction (FDR < 0.05, indicated by yellow border). Dashed line indicates LFC = 0. Heatmaps of macrophage/monocyte subset marker genes in BAL (**g**) and PBMC (**h**). Genes ranked by FDR-adjusted p-value from Wilcoxon rank-sum test (per subject average expression), comparing each subset against all others. Gene expression values are unity normalised from 0 to 1 across rows within each subset. Each column represents the average expression value for one individual. Differentially expressed genes in BAL (**i**) and PBMC (**j**) myeloid cells comparing post-COVID-19 RLA vs HCs. Gene expression values represent Z-score normalised log-normalised counts, where each gene’s expression is mean-centred and scaled across all samples. The colour scale reflects row-wise Z-scored expression, with red indicating upregulation and blue indicating downregulation in post-COVID-19 RLA relative to controls. DEGs were identified using Seurat's FindMarkers function (Wilcoxon test; log₂ fold change > 0.25; FDR < 0.05; Bonferroni correction), excluding ribosomal genes. Top annotation indicates myeloid cell type and disease group (HC = yellow, post-COVID-19 RLA = purple); right annotation categorises DEGs by pathway.

HC, healthy controls; RLA, post-COVID-19 residual lung abnormalities; LFC, log₂ fold change; and FDR, false discovery rate; DEGs, differentially expressed genes.

**Figure 3. Blood monocyte – bronchoalveolar macrophage trajectory analysis and Bronchoalveolar Lavage Fluid Proteomics**

**a–g,** Integrated analysis of myeloid cells from BAL and PBMC compartments. UMAP of Harmony-integrated BAL and PBMC myeloid cells, labelled by sample type (**a**), cell type (**b**; cell numbers in parentheses), and disease (**c**; HC = yellow; post-COVID-19 RLA = purple). The plotting order of cells in (**c**) was randomly shuffled to improve visual clarity. RNA velocity vector fields projected onto the UMAP embedding and coloured by latent time, demonstrating velocity vectors pointing from HLA-DR^lo^PDE4D^hi^ classical monocytes to profibrotic SPP1+ MoAMs (**d**). Partition-based graph abstraction (PAGA) velocity graph showing directed connectivity between HLA-DR^lo^PDE4D^hi^ classical monocytes and MoAMs (**e**). Pseudotime trajectories across integrated blood *CD14*+ monocytes and BAL MoAMs, coloured by pseudotime (**f**). Upregulated ligand–receptor signalling interactions in post-COVID-19 RLA using CellChat. BAL macrophages were defined as signalling senders and circulating monocytes as receivers. The chord diagram illustrates significantly enriched signalling pathways in post-COVID-19 RLA compared with HC (log₂ fold change > 0.2; FDR < 0.05). Interactions are grouped by cell type and colour-coded to indicate the direction and source of signalling, with ribbons representing ligand–receptor pairs contributing to enhanced macrophage–monocyte communication in post-COVID-19 RLA (**g**). **h-o,** Bronchoalveolar lavage fluid (BALF) proteomics across disease groups. Area-proportional Venn diagram showing the number of significantly differentially expressed proteins identified when comparing cases versus controls in the post-COVID-19 RLA and IPF cohorts using Alamar NULISAseq. Shared (n = 30) and disease-specific proteins (n = 35 unique to post-COVID-19 RLA; n = 15 unique to IPF) are shown (**h**). Differential protein abundance in BALF from post-COVID-19 RLA (**i**) and IPF (**j**) patients compared with HCs. Volcano plot showing log₂ fold change and statistical significance for proteins measured by Alamar NULISAseq. Differential expression was assessed using linear regression (lm) with Benjamini–Hochberg (BH) correction for multiple testing. Proteins meeting significance thresholds are highlighted (FDR < 0.01, pink; FDR < 0.05, blue) (**i-j**). Heatmap showing relative abundance of selected BALF proteins across HC, post-COVID-19 RLA, and IPF samples measured by Alamar NULISAseq. Data are shown at the patient level, with each column representing an individual patient and each row a protein. Proteins are organised into sections based on differential abundance patterns: increased in post-COVID-19 RLA compared with controls, increased in both post-COVID-19 RLA and IPF compared with controls, increased in IPF compared with controls and decreased in IPF compared with controls. Proteins are ordered by log₂ fold change in post-COVID-19 RLA (first and second sections) and IPF (third and fourth sections). Values are row-normalised and displayed as z-scores, with samples grouped by disease status (**k**). Reactome pathway enrichment of BALF proteins enriched in post-COVID-19 RLA and IPA, compared to HCs. Significantly increased proteins (FDR < 0.05; log₂FC > 0.5; linear regression with Benjamini–Hochberg correction) from post-COVID-19 RLA vs HCs and IPF vs HCs were analysed for pathway enrichment using g:Profiler2 (Reactome; with FDR correction). Bar plots show the top 15 enriched Reactome pathways per comparison, plotted as −log₁₀(FDR) (**l**). Ingenuity Pathway Analysis (IPA) upstream regulator analysis of BALF proteomic signatures. Proteins significantly differentially abundant (FDR < 0.05) in post-COVID-19 RLA versus HCs and IPF versus HCs (Alamar NULISAseq) were used as input. Only regulators predicted to be activated (positive activation z-score; |z| ≥ 2 with nominal p < 0.05) are shown, ordered by activation z-score in post-COVID-19 RLA. Dot colour denotes activation z-score and dot size denotes significance (−log₁₀(p)) (**m**). Cellular sources of differentially abundant BALF proteins in post-COVID-19 RLA mapped to BAL single-cell transcriptomic data, all BAL cells (**n**) and BAL myeloid (**o**). Proteins significantly differentially abundant in post-COVID-19 RLA versus HCs (FDR < 0.05; |log₂FC| > 0.5; Alamar NULISAseq) were mapped to gene expression in the BAL scRNA-seq atlas. Heatmaps show average log-normalised expression per cell type displayed as row-normalised z-scores with hierarchical clustering (Ward’s method).

HC, healthy controls; RLA, post-COVID-19 residual lung abnormalities; PAGA, partition-based graph abstraction; MoAMs, monocyte-derived alveolar macrophages; BALF, bronchoalveolar lavage fluid; IPF, idiopathic pulmonary fibrosis; FDR, false discovery rate; BH, Benjamini–Hochberg; and IPA, Ingenuity Pathway Analysis.

HC, healthy controls; RLA, post-COVID-19 residual lung abnormalities; IPF, idiopathic pulmonary fibrosis; COL1, collagen I; αSMA, alpha-smooth muscle actin; LFC, log₂ fold change; AUPR, area under the precision–recall curve; IQR, interquartile range; FBS, foetal bovine serum; and FDR, false-discovery rate.

**Figure 5. Single-cell and clonal analysis of T and NK cells in bronchoalveolar and blood compartments in post-COVID-19 RLA**

**a–i**, Single-cell transcriptomic analysis of T and NK cells in BAL (**a-e**) and blood (**f-i**). UMAP embedding of T and NK cells annotated by cell subtype (**a**, **f**) and (**b**, **g**) disease group (post-COVID-19 RLA = purple; HC = yellow). Cell plotting order was randomised between conditions to improve visualisation. Dot plot showing expression of canonical marker genes across T and NK cell subsets; colour indicates mean expression and dot size indicates the fraction of cells expressing each gene (**c, h**). Differential cell abundance analysis comparing post-COVID-19 RLA vs HC in BAL (**d**) and PBMC (**i**).Log₂ fold change (LFC) is shown, with red indicating increased and blue decreased abundance in post-COVID-19 RLA. Circle size reflects –log₁₀(FDR) from Wilcoxon rank-sum test with Bonferroni correction (FDR < 0.05, indicated by yellow border). Dashed line indicates LFC = 0 (**d-i**). Heatmap of differentially expressed genes across BAL T and NK cell subsets, grouped by functional pathways. Genes ranked by FDR-adjusted p-value from Wilcoxon rank-sum test (per subject average expression), comparing each subset against all others. Gene expression values represent Z-score normalised log-normalised counts, where each gene’s expression is mean-centred and scaled across all samples. The colour scale reflects row-wise Z-scored expression, with red indicating upregulation and blue indicating downregulation in post-COVID-19 RLA relative to controls. DEGs were identified using Seurat's FindMarkers function (Wilcoxon test; log₂ fold change > 0.25; FDR < 0.05; Bonferroni correction), excluding ribosomal genes. Top annotation indicates T cell type and disease group (HC = yellow, post-COVID-19 RLA = purple); right annotation categorises DEGs by pathway (**e**). **j-k,** scTCR seq analysis in BAL (**left**) and blood (**right**)**.** TCR repertoire analysis of CD4⁺ (top) and CD8⁺ (bottom) T cells in BAL (**left**) and PBMC (**right**) using GLIPH2 on each compartment separately. Each point represents a TCR cluster; point size indicates the number of unique CDR3 sequences per cluster. The x-axis denotes –log₁₀(Fisher score), indicating enrichment in HC (yellow) or post-COVID-19 RLA (purple). Clusters are ranked on the y-axis by Fisher score (**j**). TCR clone frequency distribution in BAL (**left**) and PBMC (**right**) from HC (yellow) and post-COVID-19 RLA (purple) samples. The x-axis ranks individual clones by abundance, and the y-axis shows the proportion of total TCR sequences occupied by each clone (prop). Therefore, each bar represents unique clones (**k**).

T, T cells; NK, natural killer cells; HC, health controls; RLA, post-COVID-19 residual lung abnormalities; LFC, log₂ fold change; FDR, false discovery rate; DEGs, differentially expressed genes; scTCR-seq, single-cell T cell receptor sequencing; TCR, T cell receptor; CD4, cluster of differentiation 4; CD8, cluster of differentiation 8; GLIPH2, Grouping of Lymphocyte Interactions by Paratope Hotspots 2; and CDR3, complementarity-determining region 3.

**Figure 6 Proposed model of immune–stromal interactions driving post-COVID-19 residual lung abnormalities (RLA).**

A single-cell atlas of bronchoalveolar lavage (BAL) and peripheral blood mononuclear cells (PBMCs), combined with BALF proteomics and fibroblast functional bioassays, identifies mechanisms of lung remodelling in RLA. (1) Circulating CD14⁺ monocytes are recruited to the lung via CCL–CCR and CSF1–CSF1R signalling, where HLA-DR^lo^PDE4D^hi^ classical monocytes differentiate into profibrotic *SPP1*⁺ monocyte-derived alveolar macrophages (MoAMs). (2) Immune-Stromal cross-talk: soluble fibrogenic mediators produced mainly by myeloid cells engage fibroblast receptors, compounded by direct macrophage-fibroblast communication (3) Activated fibroblasts, proliferate and differentiate into myofibroblasts, driving collagen deposition and extracellular matrix remodelling that underlies Post-COVID-19 RLA. Post-COVID-19 RLA lungs exhibit increased MoAMs and DC3 cells with reduced NK and MAIT cells, while blood shows increased Tregs and CD8⁺ ZEB2^hi^ T cells. scTCR-seq reveals oligoclonal T cell expansion in the lung compared with polyclonal expansion in blood. Proteomic analysis revealed shared and distinct proteomic signatures in post-COVID-19 RLA and Idiopathic pulmonary fibrosis.

**Supplementary Figure 8 BALF proteomic alterations in post-COVID-19 residual lung abnormalities (RLA) and idiopathic pulmonary fibrosis (IPF).**

**a**, Lollipop plot showing differential BALF protein abundance in post-COVID-19 RLA vs healthy controls (purple) and IPF vs healthy controls (green). Filled circles denote significantly differentially abundant proteins (linear regression, Benjamini–Hochberg FDR < 0.05); open circles denote non-significant proteins. Values are shown as log₂ fold change. **b**, IPA canonical pathway analysis performed on significantly differentially abundant proteins from each comparison. Pathways shown met P < 0.05 and |z-score| ≥ 2 in at least one comparison. Heatmap colours indicate predicted pathway activation (red) or inhibition (blue). **c**. Heatmap of manually curated functional pathway groups constructed from significantly differentially abundant proteins (FDR < 0.05). Colour scale indicates log₂ fold change relative to healthy controls.
