## Supplementary material for "Multi-omics reveals a monocyte-macrophage-fibroblast axis in post-COVID-19 fibroinflammatory lung remodelling": UK-ILD consortium

### UKILD Consortium

Roger Thompson, School of Medicine & Population Health, University of Sheffield; NIHR Sheffield Biomedical Research Centre, Sheffield, UK

John Simpson, Newcastle University Translational and Clinical Research Institute, Newcastle upon Tyne, UK; Respiratory Medicine Unit, Newcastle Upon Tyne Hospitals NHS Foundation Trust, Newcastle upon Tyne, UK

Beatriz Guillen Guio, Division of Public Health and Epidemiology, University of Leicester, UK; University Hospitals of Leicester NHS Trust, Leicester, UK; Centro de Investigación Biomédica en Red de Enfermedades Respiratorias (CIBERES), Instituto de Salud Carlos III, Madrid, Spain

Bibek Gooptu, NIHR Leicester Biomedical Research Centre, Division of Respiratory Sciences, Institute of Structural & Chemical Biology and Centre for Fibrosis Research, University of Leicester, Leicester, UK

Malcolm G Semple, Health Protection Research Unit in Emerging and Zoonotic Infections, Department of Clinical Infection, Microbiology and Immunology, Institute of Infection, Veterinary, and Ecological Sciences, University of Liverpool, Liverpool, UK; Department of Respiratory Medicine, Liverpool Institute of Child Health and Wellbeing, Alder Hey Children's NHS Foundation Trust, Liverpool, UK.

Emma K. Denneny, UCL Respiratory, University College London, London, UK; The Interstitial Lung Disease Service, University College London Hospitals NHS Foundation Trust, London, UK; NIHR University College London Hospitals Biomedical Research Centre

Fasihul Khan, University Hospitals of Leicester NHS Trust, Leicester, UK

Fergus Gleeson Oxford University Hospitals NHS Foundation Trust, Oxford, UK

Ian P. Hall, NIHR Nottingham Biomedical Research Centre, Nottingham, United Kingdom; Translational Medical Sciences, School of Medicine, University of Nottingham, Nottingham, United Kingdom; Respiratory Medicine, Nottingham University Hospitals Trust, Nottingham, United Kingdom

Jennifer K. Quint, School of Public Health, Imperial College London, London, UK

Jim M. Wild, School of Medicine & Population Health, University of Sheffield, Sheffield, UK

John E. Pearl, Division of Microbiology and Infection, College of Life Sciences, University of Leicester, Leicester, UK

John F. Blaikley, Manchester University NHS Foundation Trust, Manchester, UK; School of

Bioological Sciences, University of Manchester, Manchester, UK

J. Kenneth Baille, Intensive Care Unit, Royal Infirmary Edinburgh, Edinburgh; Roslin Institute, University of Edinburgh, Edinburgh, UK and Baillie Gifford Pandemic Science Hub, Centre for Inflammation Research, University of Edinburgh, 47 Little France Crescent, Edinburgh, UK, EH16 4TJ Roslin Institute, University of Edinburgh, Easter Bush, Midlothian EH25 9RG
Intensive Care Unit, Royal Infirmary of Edinburgh, Little France Crescent, Edinburgh EH16 4SA

Joseph Jacob, Satsuma Lab, UCL Hawkes Institute, University College London, UK; UCL Respiratory, University College London, UK

Karen Piper Hanley, Faculty of Biology, Medicine and Health, University of Manchester, Manchester, UK

Laura Fabbri, Exeter South West Peninsula Interstitial Lung Disease Service, Academic Department of Respiratory Medicine, Royal Devon University Healthcare NHS Foundation Trust, Exeter, UK

Laura Saunders, Division of Clinical Medicine, School of Medicine and Population Health, The University of Sheffield, NIHR Biomedical Research Centre, Sheffield, UK; Insigneo Institute for In Silico Medicine, The University of Sheffield, Sheffield, UK

Lisa G. Spencer Aintree Chest Centre, Aintree Hospital, Liverpool, UK

Louise V. Wain Division of Public Health and Epidemiology, University of Leicester, Leicester, UK; University Hospitals of Leicester NHS Trust, Leicester, UK

Mark G. Jones, NIHR Southampton Biomedical Research Centre, Clinical & Experimental Sciences, Faculty of Medicine, University of Southampton, UK

Mark Spears, Department of Respiratory Medicine, Perth Royal Infirmary and Ninewells Hospital, NHS Tayside & School of Medicine, University of Dundee, Dundee, UK

Michael A. Gibbons, NIHR Exeter Biomedical Research Centre, University of Exeter; Royal Devon University Healthcare Foundation NHS Trust, Exeter, UK

Nazia Chaudhuri, Faculty of Life and Health Sciences, School of Medicine, Ulster University, Magee Campus, Londonderry, UK

Neil A. Hanley, College of Medicine & Health, University of Birmingham, Edgbaston, Birmingham, B15 2TT, UK; University Hospitals Birmingham NHS Foundation Trust, Birmingham, B15 2GW, UK; Faculty of Biology, Medicine and Health, University of Manchester, Manchester, UK

Peter M. George, National Heart and Lung Institute, Imperial College London; Royal Brompton and Harefield Clinical Group, Guy’s and St Thomas’ NHS Foundation trust, London, UK

Pilar Rivera-Ortega, Southwest Interstitial Lung Disease Service, Royal Devon and Exeter Hospital,Royal Devon University Healthcare Foundation NHS Trust; NIHR Exeter Biomedical Research Centre, Exeter, UK

Richard J. Allen, Division of Public Health and Epidemiology, University of Leicester, Leicester, UK; University Hospitals of Leicester NHS Trust, Leicester, UK

Simon LF Walsh, Royal Brompton and Harefield Clinical Group, Guy’s and St Thomas’ NHS Foundation trust, London, UK

Simon R Johnson, Centre for Respiratory Research, School of Medicine, University of Nottingham, Nottingham, UK

Valerie Quinn, National Heart and Lung Institute, Imperial College, London, London, UK

Gisli Jenkins, National Heart and Lung Institute, Imperial College, London, London, UK
